# Quantifying Cell-State Densities in Single-Cell Phenotypic Landscapes using Mellon

**DOI:** 10.1101/2023.07.09.548272

**Authors:** Dominik Otto, Cailin Jordan, Brennan Dury, Christine Dien, Manu Setty

## Abstract

Cell-state density characterizes the distribution of cells along phenotypic landscapes and is crucial for unraveling the mechanisms that drive cellular differentiation, regeneration, and disease. Here, we present Mellon, a novel computational algorithm for high-resolution estimation of cell-state densities from single-cell data. We demonstrate Mellon’s efficacy by dissecting the density landscape of various differentiating systems, revealing a consistent pattern of high-density regions corresponding to major cell types intertwined with low-density, rare transitory states. Utilizing hematopoietic stem cell fate specification to B-cells as a case study, we present evidence implicating enhancer priming and the activation of master regulators in the emergence of these transitory states. Mellon offers the flexibility to perform temporal interpolation of time-series data, providing a detailed view of cell-state dynamics during the inherently continuous developmental processes. Scalable and adaptable, Mellon facilitates density estimation across various single-cell data modalities, scaling linearly with the number of cells. Our work underscores the importance of cell-state density in understanding the differentiation processes, and the potential of Mellon to provide new insights into the regulatory mechanisms guiding cellular fate decisions.

## Introduction

Cell differentiation is a dynamic process that underpins the development and function of all multicellular organisms. Understanding how cells are distributed along differentiation trajectories is critical for deciphering the mechanisms that drive cellular differentiation, pinpointing the key regulators and characterizing the dysregulation of these processes in disease. Cell-state density is a representation of this distribution of cells and is impacted by biological process spanning proliferation to apoptosis (**Fig. 1B**, **Supplementary Fig. 1A-C**). For instance, proliferation can increase the number of cells in a state, resulting in high cell-state density (**Fig. 1B**). Cells converge to checkpoints that ensure the fidelity of the differentiation process, also leading to high cell-state density (**Fig. 1B**). In contrast, transcriptional acceleration, as seen in rare transitory cells, lead to lower cell-state density (**Fig. 1B**). Finally, apoptosis decreases the number of cells in a state, also resulting in low cell-state density (**Fig. 1B).** As a result of these influences, cell-state density of differentiation landscapes is likely not uniform (**Fig. 1A**) but exhibit rich heterogeneity of high- and low-density regions (**Fig. 1B**).

**Figure 1:**
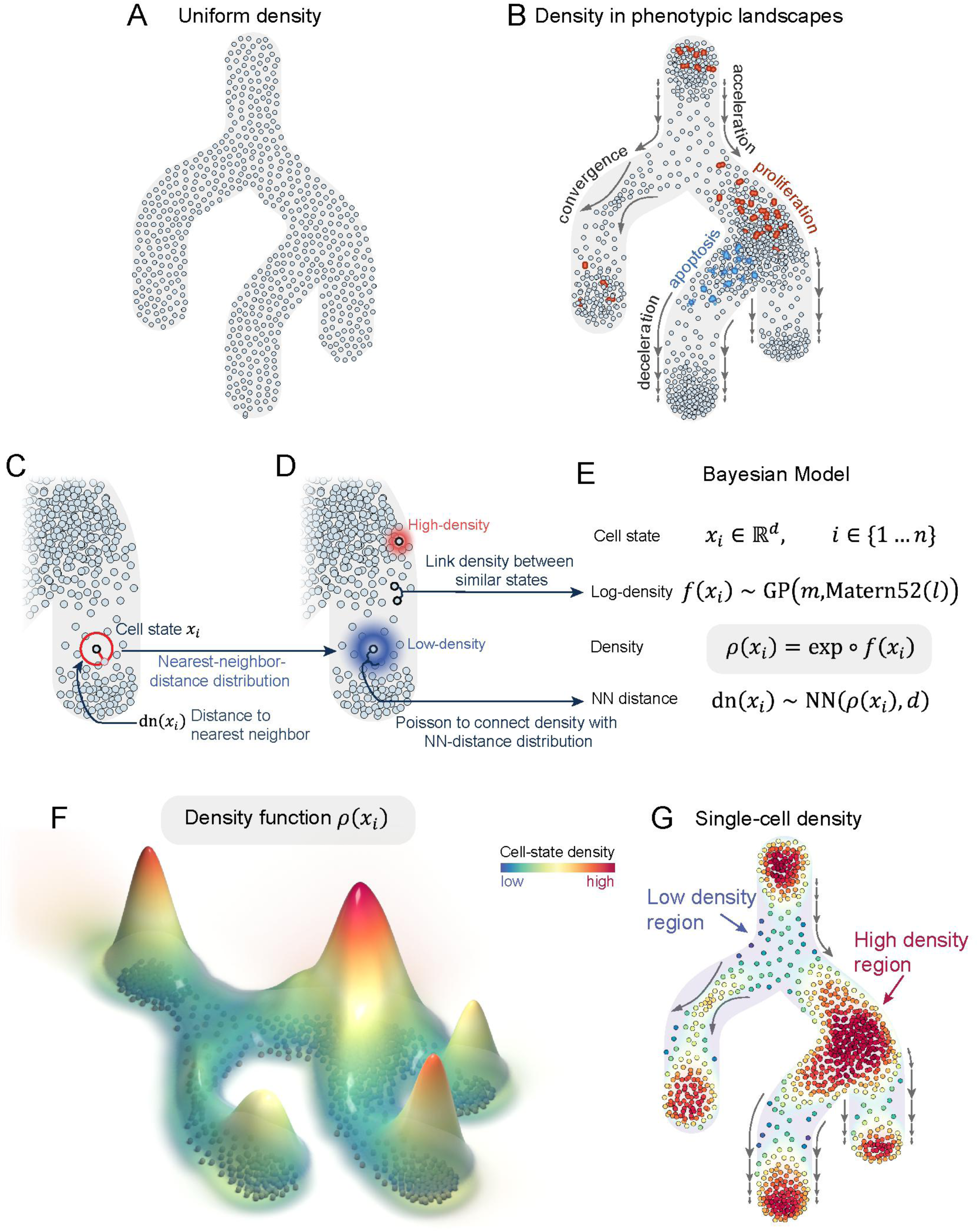
Illustrative diagram detailing the principles and processes of Mellon. A-B. An abstract depiction of a cellular differentiation landscape with cells uniformly distributed along its branches, representing a scenario not commonly found in biological systems. Diverse biological phenomena, as depicted in (B), impact cell-state density: apoptosis, acceleration, and divergence of cell-state changes lead to a decrease in density, while proliferation, deceleration, and convergence of cell-state changes increase density. Therefore, heterogeneity in cell-state densities is a norm rather than an exception in differentiation landscapes. C. Subset of cells with heterogeneous density are highlighted to illustrate the influence of biological factors in (B). Color gradient signifies the nearest-neighbor distribution around two example cells - one in a high-density state with a tighter distribution (red gradient) and another in a lower-density state with a broader distribution (blue gradient). D. Bayesian model employed by Mellon for density inference, underpinning the connection between the density estimation between neighboring cells using a Gaussian process and the log-density function as its random variable. Arrows relate the examples in panel C with their corresponding equations in D. E-F. Depict the resulting continuous density function from Mellon’s inference process over the set of cells in B. E: Density function is visualized as a 3D landscape, where the z-axis represents density, and individual cell states are illustrated as spheres at the base. F color-codes the cell states from B according to their inferred densities, overlaying these with a translucent representation of the continuous density in the background. Examples of high- and low-density regions are highlighted.

Single-cell studies have underscored the importance of the heterogeneous nature of cell-state density in single-cell phenotypic landscapes^1–3^. Rapid and coordinated transcriptional acceleration leading to low-density transitory states connecting higher-density regions have been demonstrated to be a hallmark of developmental progression in diverse biological contexts from plants to humans^4–7^. Rare transitory cells have also emerged as critical entities in the processes of differentiation^1^, reprogramming^8^, and the emergence of metastasis^9^. Despite the central importance of cell-state density, current approaches for density estimation in single-cell data often produce noisy results and struggle to provide biologically meaningful interpretation (**Supplementary Fig. 2**).

Here, we introduce Mellon, a novel computational algorithm to estimate cell-state density from single-cell data (**Fig. 1C-G**). The core principle of Mellon is based on the intrinsic relationship between neighbor distances and density, whereby distribution of nearest neighbor distances is linked with cell-state density using a Poisson distribution (**Fig. 1C**). Mellon then connects densities between highly similar cell-states using Gaussian processes to accurately and robustly compute cell-state densities that characterize single-cell phenotypic landscapes (**Fig. 1D-E**). Unlike existing approaches that interpret single-cell datasets solely as a collection of discrete cell states, Mellon infers a *continuous density function* across the high-dimensional cell-state space (**Fig. 1F**), capturing the essential characteristics of the cell population in its entirety. The density function can also be used to determine cell-state densities at single-cell resolution (**Fig. 1G**). Mellon is designed to efficiently scale to increasingly prevalent atlas-scale single-cell datasets and can be employed to infer cell-state density from diverse single-cell data modalities.

We applied Mellon to dissect the density landscape of human hematopoiesis, revealing numerous high-density regions corresponding to major cell types, intertwined with low-density, rare transitory cells. We discovered a strong correlation between low-density regions and cell-fate specification, suggesting that that lineage specification in hematopoiesis is driven by accelerated transcriptional changes. Exploration of the open chromatin landscape during lineage specification hinted at the role of enhancer priming in facilitating these transcriptional changes. Furthermore, extending Mellon’s framework to time-series datasets enabled us to compute time-continuous cell-state densities and interpolate cell-state densities between observed timepoints, providing a high-resolution view of the cell-state dynamics during erythroid differentiation in mouse gastrulation. Mellon, a scalable and user-friendly open-source software package, complete with documentation and tutorials, is available at **<u>github.com/settylab/Mellon</u>**.

## Results

### The Mellon modeling approach

Mellon aims to compute cell-state densities within the intricate, high-dimensional single-cell phenotypic landscapes. Two major challenges need to be resolved to estimate cell-state densities: First, the high-dimensionality of single-cell data is an inherent computational obstacle, which Mellon overcomes by leveraging the relationship between density and neighbor distances (**Methods**). The second challenge lies in ensuring the precise and reliable density estimation in low-density states, which often represent rare, transitory cells that play critical roles in a range of biological systems^1, 8–10^. To address this, Mellon employs a strategy of estimating a continuous density function over the entire single-cell landscape. This approach enhances both the accuracy and robustness of density estimation (**Methods**). Moreover, the density function encapsulates a smooth and continuous portrayal of the high-dimensional phenotypic landscape, enabling density estimation not only for individual measured cells—thus achieving single-cell resolution—but also for unobserved cell-states, offering a comprehensive depiction of the entire cell population (**Fig 1, Supplementary Fig. 1D**).

Mellon’s utilization of neighbor distances and inference of continuous density function is underpinned by two well-established principles of single-cell analysis. First, Mellon assumes that distances between cells in the chosen representation of the phenotypic landscape are biologically meaningful and thus represent a valid measure of cell-to-cell similarity. We refer to such a space as *cell-state space,* where each point signifies a distinct cell state. To construct such a representation, we employ diffusion maps^11^, a non-linear dimensionality reduction technique that has been demonstrated to reliably and robustly represent the single-cell phenotypic landscape^12, 13^. Moreover, distances within diffusion space are considered more biologically informative than relying on gene expression-based distances (such as PCA)^12–14^ due to its consideration of potential cell-state transition trajectories.

The second assumption Mellon relies on is that density changes from cell-to-cell are smooth and continuous in nature i.e., Mellon assumes that cells with high degrees of similarity possess similar densities. The inherent molecular heterogeneity of cells, primarily due to the subtle differences in gene expression, supports these smooth density transitions. Further, single-cell studies have revealed that cells experience gradual, rather than abrupt, changes in gene expression, providing empirical support for this assumption^1, 15–17^.

Mellon first computes distance to the nearest neighbor for each cell in the cell-state space. We then capitalize on the stochastic relationship between density and neighbor distances, where cells in higher density states tend to exhibit shorter distances to their nearest neighbors, whereas cells in lower density states tend to have longer distances (**Supplementary Fig 3A**). Formally, Mellon relates the nearest neighbor distance to local density through the nearest neighbor distribution by employing a Poisson point process (**Fig. 1C-D**, **Methods**). Nearest neighbor distribution describes the probability of another cell-state existing within some distance of a given cell-state. Intuitively, regions with higher density of cell-states correspond to tighter nearest-neighbor distributions, while lower densities result in broader distributions (**Fig. 1D**).

Mellon then connects densities between highly related cells to estimate a continuous density function. The true density function can be arbitrarily complex depending on the biological system. Mellon therefore employs Gaussian Process (GP) in a Bayesian model to approximate this function without assuming a specific functional form (**Fig. 1D**). GPs are a mathematical framework to model the patterns and relationships among data points and, are highly effective for scenarios where the true functional form is intricate or unknown and where observations are limited^18, 19^. GPs are thus ideally suited for density estimation from noisy single-cell data. They achieve their robustness by incorporating the smoothness assumption through a covariance kernel, facilitating sharing of information between adjacent observations. In Mellon, the covariance kernel of the GP encodes cell-state similarity and determines the influence of nearby cells on density estimates at a specific state (**Methods**). This covariance kernel is effectively computed for all pairs of cells and thus ensures the appropriate weightage of nearby cells in both high- and low-density states (**Supplementary Fig 3B-F)**. Finally, Mellon adopts a scalable Bayesian inference approach, tailored for atlas-scale single-cell datasets. The scalability is in large part achieved by employing a sparse Gaussian Process that approximates the full covariance structure using a set of landmark points (**Methods**).

The density function derived by Mellon is a continuous representation of the single-cell phenotypic landscape (**Fig. 1E, Supplementary Fig. 1D**). This function enables density estimation at single-cell resolution (**Fig. 1F)**. Visualizing cell-state densities with methods such as UMAPs (**Fig**. **1F**) simplifies the exploration of high- and low-density cell states in differentiation landscapes. Within the cell-state density landscape, we discerned what we term *regions* – connected subsets within the cell-state space with similar density characteristics. Such regions represent collections of closely related cell states that cells appear to inhabit (*high-density* regions) or traverse (*low-density* regions) during their differentiation journey **(Fig. 1F)**.

To assess Mellon’s accuracy, we generated simulated datasets composed of either discrete clusters or continuous trajectories, using mixtures of Gaussians in ten to twenty dimensions (**Supplementary Fig. 4A, D, G**). Comparing the ground truth density from the Gaussian mixtures to Mellon-inferred density demonstrated strong agreement, showcasing Mellon’s capability to accurately estimate cell-state densities in high-dimensional spaces (**Supplementary Fig. 4**).

### Density landscape of hematopoiesis with Mellon

Hematopoiesis, the process through which the blood and immune cells differentiate from hematopoietic stem cells (HSCs), provides an ideal paradigm to understand and model differentiation^20^. We therefore utilized a previously generated single-cell multiome dataset of T-cell depleted bone marrow^21^ representing human hematopoietic differentiation (**Fig. 2A**) to evaluate the performance of Mellon and interpret the inferred cell-state densities.

**Figure 2:**
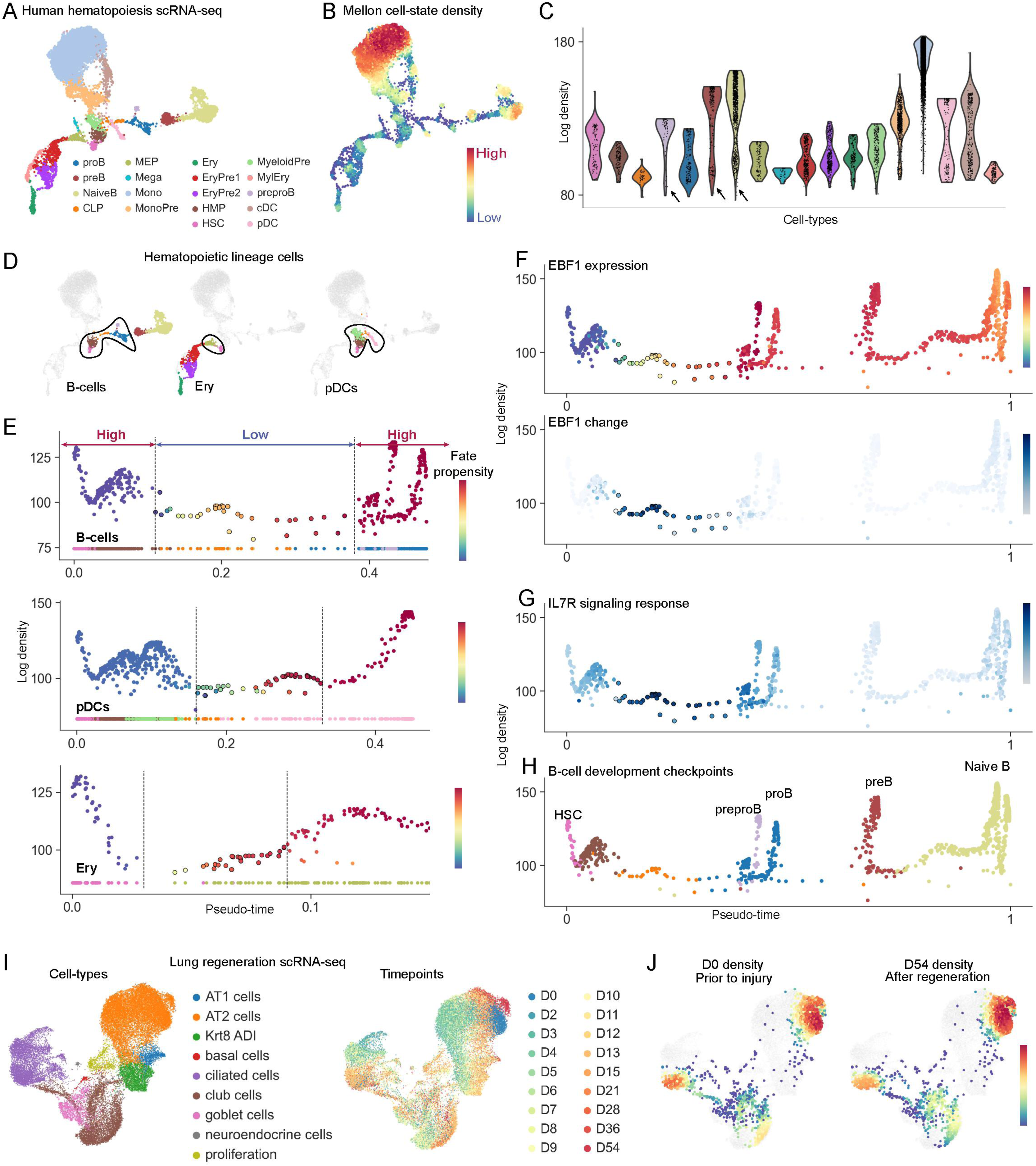
Mellon reveals the density landscape of human hematopoietic differentiation. A. UMAP representation of the scRNA-seq dataset of T-cell depleted bone marrow^21^ colored by cell-types. B. Same UMAP as (A), colored by Mellon cell-state density C. Violin plots to compare cell-state densities among different hematopoietic cell-types. Arrowheads indicate example cell-types with high variability in density. D. UMAPs as in (A), highlighted by cells of the different lineages, left to right: B-cells, Erythroid lineage cells and plasmacytoid dendritic cells (pDCs). Lineage cells were selected based on cell-fate propensities. Cells spanning hematopoietic stem-cells to fate committed cells along each lineage. E. Plots comparing pseudotime ordering and log density during the fate specification of each lineage. Top to bottom: B-cells, pDCs and Erythroid lineages. Cells are colored by Palantir fate propensities, which represent the probability of each cell differentiating to the corresponding lineage. Points at the bottom of each plot are colored by cell-type. Subset of cells along each lineage spanning hematopoietic stem cells to fate committed cells are shown. Dotted lines indicate the low-density region within which fate specification takes place and were added manually. F. Plots comparing pseudotime and log density for all cells of the B-cell trajectory colored by EBF1 MAGIC^2^ imputed expression (top) and EBF1 local variability in gene expression (bottom). G. Same as (F), with cells colored by signature scores for IL7R response genes. H. Same as (F), with cells colored by cell-types. Density peaks correspond to well-characterized checkpoints during B-cell differentiation. I. UMAP representation of the scRNA-seq dataset of lung regeneration^28^. Cells are colored by cell-type (left) and by timepoint of measurement (right). D0 is prior to injury and all subsequent timepoints show recovery from injury. J. UMAPs colored by density at D0 (left) and density at D54 (right). Cells from D0 and D54 are colored by density with cells from other timepoints in grey.

We used diffusion maps to derive a representation of hematopoietic cell-states and applied Mellon to infer density in this high-dimensional cell-state space (**Fig. 2B**). The resulting density landscape exhibited considerable heterogeneity, with numerous high-density regions, corresponding to major cell types, interconnected by low-density regions indicative of rare transitory cells (**Fig. 2B**). Monocytes, for instance, exhibited the highest cell-state density (**Fig. 2C**), which is consistent with their status as the most prevalent leukocyte in hematopoiesis and their emergence from bone marrow in a naïve state^22^. Intriguingly, we observed noticeable fluctuation in density within several cell-type clusters, suggesting an inherent heterogeneity, a nuance often masked when cells are grouped together by clustering methods (**Fig. 2C**).

For a more comprehensive understanding of the hematopoietic density landscape, we utilized our trajectory detection algorithm Palantir^14^ to determine a pseudo-temporal ordering of cells representing developmental progression and cell-fate propensities that quantify the probability of each cell to differentiate to a terminal cell-type (**Supplementary Fig. 5A-B**). We compared cell-state density along pseudotime for each lineage and observed that the increase in fate propensity towards the lineage is strongly correlated with the first low-density region in each lineage (**Fig. 2D-E, Supplementary Fig. 5C-D**). Low-density regions therefore appear to be a hallmark of cell-fate specification in hematopoiesis. These low-density regions from HSCs to fate-committed cells encompasses <0.4% of the data and under <0.01% of bone marrow cells, demonstrating the ability of Mellon to identify low-frequency rare transitory cells (**Fig. 2E**).

The occurrence of low-density regions in density landscapes can be attributed to accelerated gene expression changes, divergence, or apoptosis (**Supplementary Fig. 1A-C**). Apoptosis during hematopoietic cell-fate commitment has been shown to be minimal^23^. Further, divergence or spread of cell states, while theoretically possible, would likely result in a broader distribution rather than the observed tight trajectories. Therefore, our results strongly suggest that hematopoietic lineage specification events occur through low-density regions induced by rapid and accelerated gene expression changes.

We next devised a gene change analysis procedure to identify genes with high expression change in low-density regions (**Methods**). Our procedure consists of two steps: (1) We first determine local variability for each gene, which represents the change in expression of the gene in a cell-state. Local variability for a gene is determined as follows: For each state, we computed the absolute difference in gene expression to its neighbors. The differences are normalized by distance between states and the maximum of these normalized differences is nominated as the local variability of the gene. (2) Genes are then ranked by the weighted average of local variability across cells spanning a low-density region and the flanking high-density regions. Inverse of density are used as weights to ensure genes with higher expression change in low-density regions are ranked higher. Thus, gene change analysis quantifies the influence of a gene in driving state transitions in low-density regions (**Methods**).

We applied the gene change analysis procedure to identify genes that drive hematopoietic fate specifications by selecting cells spanning hematopoietic stem-cells to fate committed cells along each lineage (**Supplementary Fig. 6A**). Upregulated genes in each lineage transition were enriched for lineage identity genes whereas downregulated genes across lineages were associated with stem cell programs (**Supplementary Fig. 6B-C)**, indicating that changes that underlie cell-fate specification in hematopoiesis occur in low-density regions. Notably, we observed genes with higher expression levels specifically in low-density states, suggesting that despite their transitory nature, certain gene regulatory programs are uniquely adapted to facilitate these transitions (**Supplementary Fig. 6**).

We next utilized Mellon densities and associated genes to investigate B-cell fate specification. Genes with high change scores in this low-density region were enriched for modulators of B-cell lineage specification with their roles traversing transcriptional regulation, intracellular signaling and cell migration. Transcription factor EBF1 had the highest change score (**Supplementary Fig. 7A**), aligning with its role as the master regulator of B-cell differentiation^24^. In fact, the upregulation of EBF1 is exquisitely localized to the low-density region between stem and B-lineage committed cells (**Fig. 2F**), with similar dynamics observed in other critical B-cell commitment regulators such as PAX5 and IL7R (**Supplementary Fig. 7B-C**). From a signaling point of view, we observed an upregulation of IL-7 responsive Stat signaling targets in the same low-density cells concurrent with IL7R upregulation (**Fig. 2G, Supplementary Fig. 7C**). These observations are consistent with previous studies that have illustrated the vital role of IL-7 driven activation of STAT5 in a rare precursor population for B-cell specification^1^. Finally, genes such as NEGR1, with documented roles in cell adhesion and migration^25^, also score high (**Supplementary Fig. 7B)**, demonstrating that the spatio-temporal continuum of B-cell differentiation within the bone marrow is executed as rapid transcriptional changes through low-density cell-states.

These findings underscore the potential of Mellon to uncover rare, biologically significant cell populations. They also demonstrate that rapid transcriptional changes that drive state transitions in low-density regions are shaped by an intricate interplay of cell-autonomous and extrinsic factors, highlighting how Mellon can help unravel this complexity.

Following fate specification, B-cell development is a highly orchestrated process where cells transition through checkpoints as they gain functional and non-self-reactive B-cell receptors^26^. We analyzed Mellon densities along pseudotime and observed that B-cell differentiation is defined by alternating high- and low-density regions (**Fig. 2H**). Using marker gene expression and gene change score analysis, we inferred that every high-density peak represents a well-characterized checkpoint, and every checkpoint corresponds to a high-density peak (**Fig. 2H**, **Supplementary Fig 7D**). This also suggests that checkpoint releases manifest as low-density states. Since apoptosis has only been extensively observed in the transition from Pre-Pro B-cells to Pre-B-cells^1, 26^, our results suggest that cells rapidly change their state upon checkpoint release until they reach the next checkpoint, where they converge to create high-density regions.

As a test of robustness of these results, we assessed Mellon’s reproducibility by computing cell-state densities for single-cell datasets of bone marrow cells from eight independent donors from the Human Cell Atlas^27^. Densities were highly consistent across the donors, demonstrating consistent observation of high- and low-density regions across the hematopoietic landscape (**Supplementary Fig. 8A-B**). Moreover, density patterns along the B-cell differentiation trajectories were also consistent between the samples, reinforcing the reliability and reproducibility of Mellon density estimates (**Supplementary Fig. 8C**).

### Versatility of Mellon cell-state densities

We investigated whether cell-state density is a fundamental property of the homeostatic system by investigating whether cell-state density is restored upon regeneration. We utilized a single-cell dataset of lung regeneration where lungs were profiled using scRNA-seq following injuries induced with bleomycin (**Fig. 2I**)^28^. We applied Mellon to compute cell-state densities before injury and upon recovery. Remarkably, we observed that the density landscape reverts to the homeostatic state upon regeneration from injury (**Fig. 2J, Fig., Supplementary Fig. 9**). This observation suggests that cell-state density, while fundamental to tissue homeostasis, is also reflective of the tissue regenerate state. As the tissue recovers from injury, the restoration of the original cell-state density landscape could serve as an indicator of successful tissue regeneration.

We further explored Mellon’s versatility by applying it to a variety of homeostatic biological systems such as pancreatic development^29^, endoderm differentiation^30^ and spatial organization of intestinal tissues ^31^. The recurring observation of high- and low-density regions across these diverse systems suggests that these patterns are a ubiquitous feature of single-cell phenotypic landscapes (**Supplementary Fig. 10**). These density variations supply a wealth of biological insight beyond abstract quantities: High-density regions across systems typically correspond to key developmental checkpoints or bottlenecks, while low-density regions often represent rare transitory cells undergoing rapid transcriptional changes to bridge the denser areas (**Supplementary Fig. 10**).

These findings emphasize the effectiveness of Mellon for accurately characterizing differentiation landscapes and highlight the importance of scrutinizing both high- and low-density regions for a holistic understanding of the differentiation processes. Mellon’s fine-grained resolution also aids the identification of rare transitory cells, a critical element of diverse biological phenomena.

### Mellon produces robust cell-state densities

We next assessed the robustness of Mellon cell-state densities across different parameters. The number of cells measured in a dataset is a crucial factor affecting the accuracy and reliability of density estimates. We performed subsampling experiments and compared the results to those obtained using the full dataset by leveraging the continuous nature of Mellon. Our subsampling experiments show that Mellon’s density estimates are highly robust to subsampling, even when reducing the number of cells by an order of magnitude across different datasets (**Supplementary Fig. 11,12**). Density estimates are also robust to variations in the number of diffusion components (**Supplementary Fig. 13**), dimensionality (**Supplementary Fig. 14**), the number of landmarks (**Supplementary Fig. 15**), and the length-scale heuristic employed for scalability (**Supplementary Fig. 16**). These findings underscore the reliability of Mellon’s density estimation approach, which can provide accurate and robust results even with limited data.

Finally, we compared Mellon to existing approaches for cell-state density estimation. Densities have been approximated as the inverse of distance to kth nearest neighbor^2, 14^ due to computational complexity. However, due to the inherent noise and sparsity of scRNA-seq data, these approaches often fail to generate robust density estimates (**Supplementary Fig. 2A-B**). The characteristic high- and low-density regions identified by Mellon could not be demarcated by densities estimated solely from nearest neighbor distances (**Supplementary Fig. 2B-D**). Given this noise, 2D embeddings, such as UMAPs, have been widely utilized for density computation. While such embeddings are effective for visualization, the low-dimensionality restricts their capacity to encapsulate all biologically significant variability. The UMAP density estimates for the T-cell depleted bone marrow data are dominated by the most dominant cell-type, i.e., monocytes (**Supplementary Fig. 2C**) with no discernable variability in the other lineages (**Supplementary Fig. 2D**). Thus cell-state density estimation using Mellon substantially outperforms existing approaches in accuracy, and biological interpretability.

### Enhancer priming as a catalyst for rapid transcriptional changes in low density cell-states

We next turned our attention to the mechanisms that regulate the rapid transcriptional changes that generate rare transitory cells during lineage specification. Previous studies have identified extensive priming of lineage-specifying genes in hematopoietic stem cells, where gene loci are maintained in an open chromatin state through pre-established enhancers, even in the absence of gene expression^32–34^. Moreover, enhancer priming has been implicated to play a role in rapid transcriptional responses to stimuli in hematopoietic cells^35^.

Building on these findings, we hypothesize that the rapid upregulation of lineage specifying genes as cells transition from HSCs (a high-density region) to fate committed cells (another high-density region) is in part driven by enhancer priming. In this scenario, the loci of lineage-specifying genes are maintained in an accessible state in HSCs through pre-established enhancers. A combination of cell-autonomous and extrinsic factors trigger the upregulation of a small set of master regulators, which in turn rapidly upregulate the expression of lineage-specifying genes in a coordinated manner. Thus, the combined activity of pre-established enhancers in HSCs and lineage-specific enhancers established by master regulators could produce the rapid transcriptional changes that underpin rare transitory cells in low-density states (**Supplementary Fig. 17**).

We used the transition from HSC to B-cells as the case-study (**Fig. 3C**) to test our hypothesis. We leveraged the multiomic nature of our T cell depleted bone marrow dataset, with measurements of both expression (RNA) and chromatin accessibility (ATAC) in the same single cells (**Supplementary Fig. 18**). The first step is to delineate the primed and lineage-specific peaks associated with a gene. The noise and sparsity of scATAC data means that determination of individual peak accessibility at single-cell level is extremely unreliable^36^. Therefore, we devised a procedure to disentangle primed and lineage-specific peaks associated with a gene using different levels of abstractions (**Supplementary Fig. 19A, Methods**): First, we used our SEACells algorithm^21^ to aggregate highly-related cells into metacells and identified the set of peaks with accessibility that significantly correlate with gene expression (**Fig. 3A**). We then identified the subset of these peaks with greater accessibility in B-cells compared to other lineages by comparing accessibility between cell-types at the metacell resolution. This approach ensures the exclusion of ubiquitous and low-signal peaks while retaining peaks that are important for B-cell fate specification. Finally, we classified each peak as primed if it was accessible in HSCs, and as lineage-specific if its accessibility was B-cell restricted (**Fig. 3B**). We verified that the accessibility of lineage-specific peaks was near exclusive to B-cells and that of primed peaks were higher in HSCs and B cells (**Supplementary Fig. 19B-C**).

**Figure 3:**
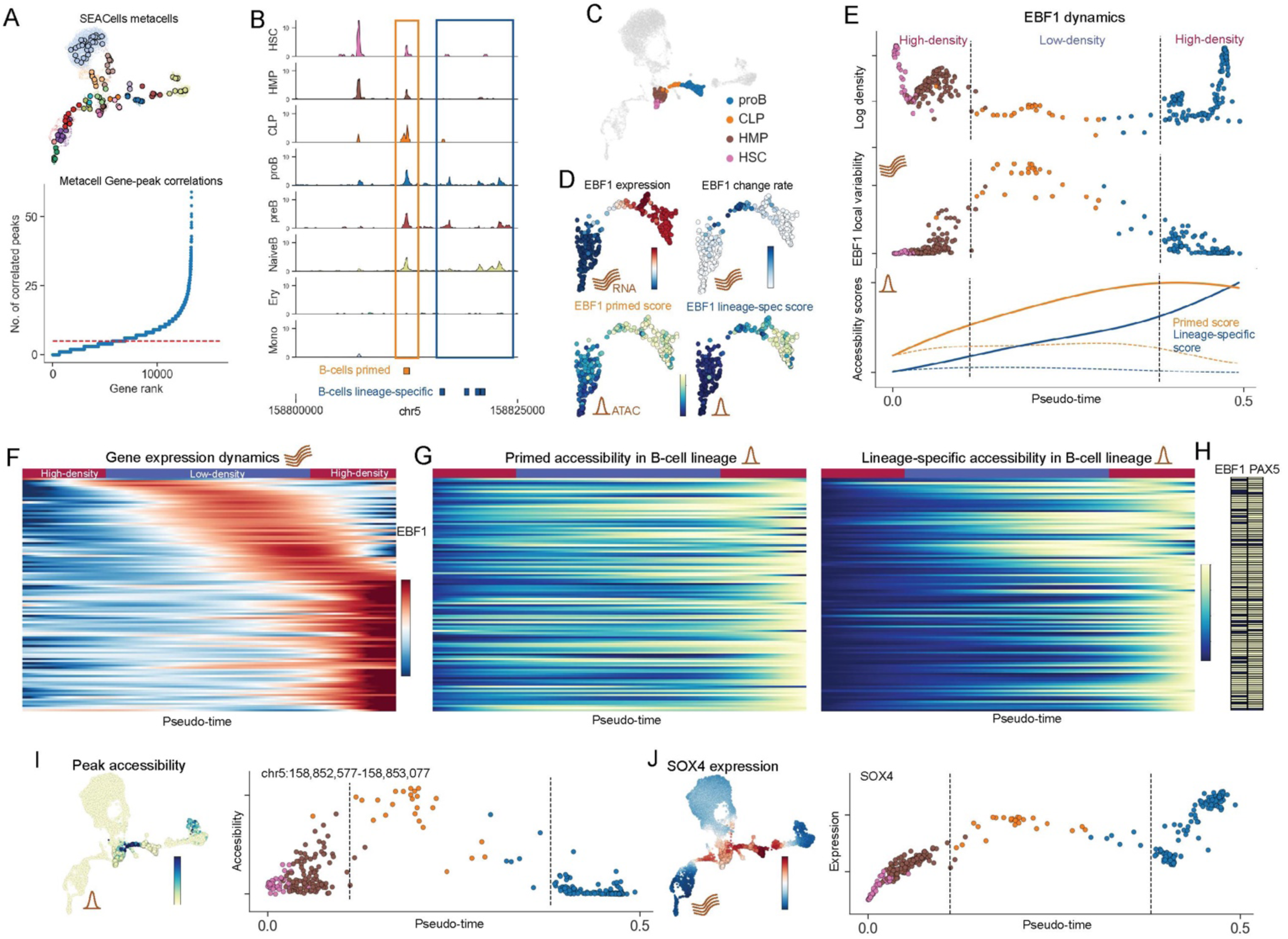
Dynamics of chromatin accessibility and gene expression during B-cell fate specification. A. Top: UMAP colored by cell-types and highlighted by SEACells^21^ metacells. Bottom: Plots showing the number of peaks significantly correlated with each gene. The correlations were computed using SEACells^21^ metacells. B. Coverage plots highlighting examples of B-cell primed (in orange) and B-cell lineage specific peaks (in blue). The genomic region is part of the EBF1 gene locus C. UMAP colored by cell types included in B-cell specification. The full dataset is shown in grey. D. UMAP colored by EBF1 MAGIC imputed expression, EBF1 local variability, EBF1 primed accessibility scores and EBF1 lineage-specific accessibility scores. The subset of cells involved in B-cell specification (C) are shown. E. Top: Plots comparing pseudotime and Mellon density for the B-lineage cells, colored by cell-type. Middle: Plots comparing pseudotime and EBF1 local change for the B-lineage cells, colored by cell-type. Bottom: Solid lines show the trend of primed and lineage-specific accessibility scores for EBF1 in B-cell lineage. Dotted lines show the corresponding trends in the erythroid lineage. Vertical dotted lines show high- and low-density regions selected manually. F. Heatmaps with z-score expression of genes with high change scores and upregulation during B-cell specification. Genes are sorted based on their expression along pseudotime. Genes with at least 1 primed and at least 1 lineage-specific peak from **Supplementary** Fig. 20A were used. G. Heatmaps of primed (left) and lineage-specific (right) accessibility scores for genes in (F) in the same order. Scores were scaled to maximum of 1 along the trend. H. Matrix indicating whether the genes in (F) are predicted targets of EBF1 or PAX5 using Insilico-ChIP^37^. I. Left: UMAP colored by MAGIC imputed accessibility of the single ATAC peak (chr5:158,852,577-158,853,077) with highest change score in EBF1 correlated peaks. Right: Plot comparing pseudotime to peak accessibility for cells during B-cell specification in (C). J. Left: UMAP colored by SOX4 MAGIC imputed expression. Right: Plot comparing pseudotime to gene expression for cells during B-cell specification in (C).

We identified the set of genes with high change scores in B-cell specification using our gene change analysis procedure (**Supplementary Fig. 20A**). We then used the subset of these genes with upregulation in B-cell lineage and those with at least 5 peaks correlated with expression to test our hypothesis. >80% of these genes were associated with at least one primed peak and one lineage-specific peak (**Supplementary Fig. 20A**), implicating enhancer priming as vital to their upregulation. In contrast, none of the genes associated with the erythroid fate specification demonstrated B-cell primed peaks. To characterize the dynamics of these peaks during lineage specification, we computed two accessibility scores for each gene at single-cell resolution: (i) primed score, defined as the aggregated accessibility of all primed peaks correlated with the gene and (ii)lineage-specific score, defined as the aggregated accessibility of all lineage-specific peaks correlated with the gene (**Methods**). We first used these scores to examine the dynamics of EBF1, the gene with the highest change score in the low-density region of B-cell fate specification (**Fig. 3D**) and the master regulator of B-cell differentiation^24^. We observed that primed peaks were open in stem cells as expected and increased in accessibility as B-cell fate was specified (**Fig. 3E**, orange line). This was followed by the establishment and stabilization of lineage-specific peaks (**Fig. 3E**, blue line) and finally lineage-specific upregulation of EBF1, highlighting the role played by enhancer priming in the upregulation of EBF1. We next examined the genes upregulated in B-cell lineage specification with primed and lineage-specific peaks, and observed a similar pattern to EBF1, along with a coordinated upregulation that follows EBF1 expression (**Fig. 3E-F, Supplementary Fig. 20C**). Finally, we used in-silico ChIP^37^ to identify that almost every gene in our gene set is a predicted target of either EBF1 or PAX5 (**Fig. 3H**), consistent with our hypothesis and the proposed role of EBF1 as a trigger for a “big-bang” of B-cell development^38^.

Our results support a mechanism where enhancer priming and subsequent activation of master regulators lead to a rapid and coordinated upregulation of genes, resulting in the emergence of rare transitory cells that confer lineage identity. These results highlight the importance of taking cell-state density into consideration for understanding gene regulatory networks that drive cell-fate specification. Our approach to determine primed and lineage-specific accessibility scores for each gene utilizes the history of peak establishment, a feature unaccounted for by most current techniques, which tend to aggregate all peaks in proximity of a gene to derive a single gene score^32, 36^ (**Supplementary Fig. 20D-E**). Finally, the expression and accessibility trends were determined using Gaussian process with the function estimator implemented in Mellon, highlighting another utility of the Mellon framework (**Supplementary Fig. 21**, Methods).

### Identification of master regulators with Mellon

While master regulators have been identified for several hematopoietic lineages, the mechanisms controlling lineage-specific upregulation of these master regulators remain largely elusive. To investigate whether the regulation of EBF1 could be clarified through cell-state density, we adapted our approach to compute gene-change scores to rank the EBF1 correlated peaks by their accessibility change score in the low-density region of B-cell specification (**Methods**). Interestingly, the top peak in this analysis was almost exclusively accessible in the low-density region (**Fig. 3I**). We employed in silico ChIP to identify transcription factors with a strong predicted signal to bind this peak and observed that top 10 enriched motifs were exclusively comprised of IRF and SOX motifs. Interestingly, the increase in accessibility in the peaks is concurrent with upregulation of the transcription factor SOX4 (**Fig. 3J**), a known regulator of EBF1 during B-cell development^39^. These results clarify the temporal order of transcriptional events where upregulation of SOX4 leads to lineage-specific expression of EBF1 to establish B-cell fate and also suggest the specific set of regulatory elements that drive this mechanism.

The strong association of EBF1 expression with low-density transition (**Supplementary Fig. 20A**) and the high number of expression-correlated peaks (**Supplementary Fig. 19E**), coupled with the coordinated upregulation of its targets (**Fig. 3H**), suggests a paradigm for identifying master regulators via Mellon densities. Additionally, identifying peaks whose accessibility changes are strongly associated with cell-state density can offer insights into the regulation of the master regulators themselves.

### Exploring Time-Series Single-Cell Datasets with Mellon to Understand Mouse Gastrulation

Time-series single-cell datasets are invaluable for understanding the intricate dynamic processes driving development, as they provide snapshots of the changes in cell-type and cell-state compositions during a fast-changing process. Although various computational methods exist to model trajectories using time-series data^8, 40–43^, they typically represent these changes as discrete steps between measured timepoints and thus are limiting when studying inherently continuous processes like embryonic development. To better represent these processes, we investigated if we could utilize Mellon’s continuous density functions to construct a *time-continuous* cell-state density function. This function will span not just the observed timepoints, but can also interpolate densities at unobserved times, enabling a truly continuous view of the shifting cell-state density landscape during development.

We used the mouse gastrulation atlas^44^, a single-cell dataset of 116,312 cells spanning gastrulation and early organogenesis (E6.5-E8.5) (**Fig. 4A**) for exploration of time-continuous densities. We first applied Mellon to each timepoint individually and observed considerable variability in cell-state densities over time (**Fig. 4B, Supplementary Fig. 22**). Interestingly, we observed that the emergence of new cell types or lineages was often marked by a low-density transition (**Fig. 4B**), highlighting the “fits and starts” nature of developmental progression^5^.

**Figure 4:**
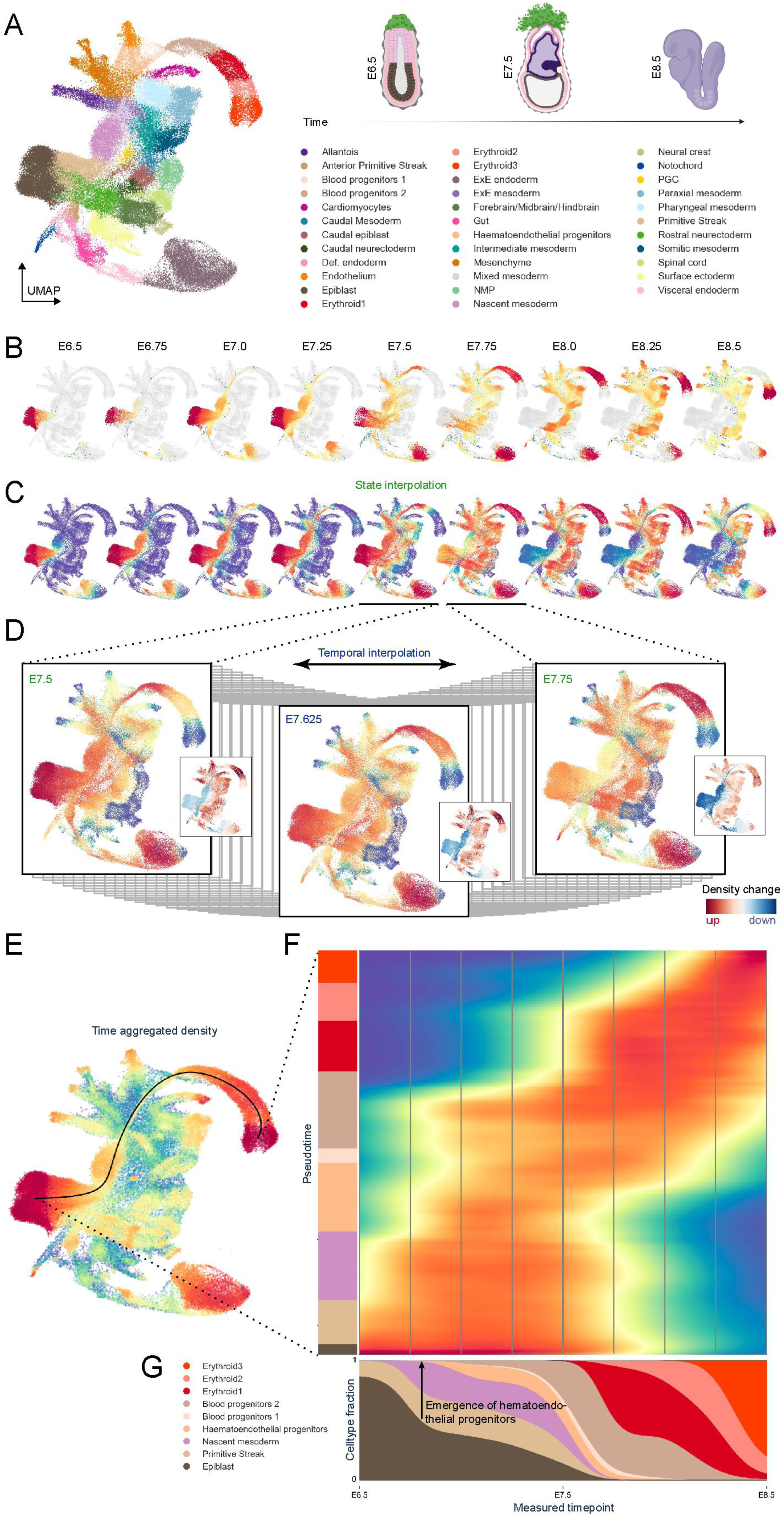
Depiction of time-continuous cell-state density estimation during mouse gastrulation using Mellon. A. UMAP representation of the mouse gastrulation dataset^44^. Illustrations on the right show a diagrammatic overview of the mouse embryo during gastrulation from E6.5 to E8.5, providing context to the developmental progression. *Created using BioRender*. B. UMAPs colored by Mellon cell-state density at each measured timepoint, demonstrating variability in cell-state densities within each observed timepoint. C. UMAPs colored by state-interpolated densities, derived from densities from (B), but evaluated across all cells. This showcases the potential of Mellon for extrapolating cell-state densities beyond directly sampled cell states. D. Illustration of time-continuous density on UMAP for measured (E7.5, E7.75) and interpolated (E7.25) timepoints, further demonstrating the application of Mellon in interpolating cell-state densities beyond measured timepoints. Smaller accompanying UMAPs denote the temporal rate of change in cell-state density, with red signifying increasing density (enrichment) and blue indicating decreasing density (depletion). E. UMAP colored by cell-state density inferred using all-cells without using temporal information. Trend highlights the erythroid trajectory. F. Heatmap displays the time-dependent cell-state densities along the trajectory (pseudotime on the y-axis and real-time on the x-axis), with vertical grey lines signifying the measured timepoints. G. Marginal plot illustrating the proportional composition of cell-types along the erythroid trajectory at each timepoint, derived by integrating density in F across the trajectory segment associated with each specific cell type.

Mellon’s capacity to generate a continuous density function enables it to estimate densities for cell states that were not part of the training data (**Fig. 1**). To demonstrate this, we used the density function associated with each specific time point to calculate densities across cells of all timepoints (**Fig. 4C**). In essence, we estimated the likelihood of each cell state being observed at a different time point. This unique feature of Mellon allows for comparison of cell-state densities across various developmental stages by calculating the correlation between the pair of time-point densities within the same cell state (**Supplementary Fig. 23**). Interestingly, embryonic stage E7.75 was least similar to neighboring timepoints, indicating the completion of gastrulation and onset of organogenesis at E7.75 (**Supplementary Fig. 23A-D**).

We next constructed a time-continuous cell-state density of mouse gastrulation by incorporating measurement time as a covariate. We devised a procedure to ensure that the covariance of measurement times between cells reflects the empirically observed correlation between timepoint densities (**Methods**, **Supplementary Fig. 23E-F**) to construct a density function that is continuous in both *time* and *cell-state.* Therefore, we can estimate cell-state densities at any desired timepoint situated between the measured instances (**Fig. 4D, Supplementary Video 1**). Thus, by leveraging the temporally related data, we enhanced the cell-state distribution of individual time points - a cell state present in preceding and following time points is likely to exist in the current time point, even if it hasn’t been directly observed.

To validate this approach, we performed leave-one-out cross-validation by comparing density computed exclusively from a timepoint with the interpolated density computed by omitting the same timepoint. The two densities are highly correlated even for timepoint E7.75 (**Supplementary Fig. 24**), which is least similar to its neighbors, providing a clear validation of our approach.

Importantly, our time-continuous approach also enables the quantification of rates of density change. By taking the derivative of the time-continuous density along the time axis, we can assess the rates of enrichment or depletion for every cell state at any time (**Fig. 4D, Supplementary Video 1**). Our analysis reveals that the initial phase of gastrulation is predominantly characterized by growth, with a nearly constant abundance of epiblast and primitive streak cells—a finding in line with prior studies noting high proliferation^45^ (**Supplementary Fig. 25A**). Following this phase, a sharp transition occurs at E7.5, where a rapid decline of epiblast and primitive streak cells signals the completion of the gastrulation process (**Supplementary Fig. 25B-C**). Finally, another transition at E8.375 marks the emergence of ectodermal and endodermal structures, accompanied by a concomitant decline in their respective progenitors (**Supplementary Fig. 25D**). These findings underscore the power and potential of employing time-continuous cell-state density modeling to provide a high-resolution depiction of the developmental process in its entirety.

The application of time-continuous cell-state densities can also offer insight into the dynamics of cell abundance along specific developmental lineages. As a case study, we chose to investigate erythropoiesis during gastrulation, given its well-understood process. Using the full gastrulation atlas and Palantir^14^ we identified cells predisposed to differentiate into erythroid lineage and derived a pseudo-temporal ordering of these cells (**Supplementary Fig 26**). Leveraging our time-continuous cell-state density function, we approximated densities along pseudo-time, which revealed a continuous progression of cells toward the erythroid state (**Fig. 4F**). Interestingly, there is a strong alignment between pseudotime and real time indicating a linear dependency in the erythroid lineage. Note that the persistent high density of early epiblast cells likely represents cells differentiating into cell types other than the erythroid lineage.

Further, the dynamics of cell-type proportion along real time can be investigated by computing the marginal of the density representation that contrasts real-time versus pseudo-time (**Fig. 4G**). This visualization allowed us to precisely pinpoint the timespan during which hematoendothelial progenitors, the earliest precursors of erythroid cells, emerge from the nascent mesoderm (**Fig. 4G**). Notably, the proportion of hematoendothelial cells remains relatively stable across time, indicating their transient presence without expansion in the cell population. In stark contrast, blood progenitor cells (Type 2) undergo a substantial increase in their proportion following their emergence, suggesting a period of accelerated cell division. Therefore, our time interpolation offers valuable insights into the progression of cell type abundances and allows for high resolution predictions of the emergence of specific cell types.

Our results showcase Mellon’s capability to provide a comprehensive, time-continuous perspective on cell-state densities during development and reprogramming.

### Mellon infers densities from single-cell chromatin data

Single-cell chromatin profiling techniques such as single-cell ATAC-seq^16^, CUT&Tag^46, 47^, and sortChIC-seq^48^ are revolutionizing the study of interplay between gene expression and chromatin landscape in disease and differentiation. We developed Mellon to be adaptable to different single-cell modalities, making it a valuable addition to the computational toolkit for these emerging techniques. Given their robust representation of cell-states, we use diffusion maps for deriving a cell-state space for density inference through Mellon. Diffusion maps rely only on distances between similar cells and thus can be constructed for most single-cell data modalities following appropriate pre-processing^21^.

To evaluate Mellon’s adaptability to scATAC-seq data, we computed diffusion maps from the ATAC modality of the T-cell depleted bone marrow dataset^21^ and applied Mellon for cell-state density inference. Similar to gene expression, Mellon reveals substantial chromatin-state density variability (**Supplementary Fig. 27A**). High- and low-density states corresponded respectively to major cell-types or checkpoints and rare transitory cells (**Supplementary Fig. 27A**). Applied to a mouse model of lung adenocarcinoma^49^, we observed extensive chromatin-state density variability amongst cells of the primary tumors, with the transition to metastasis associated with a sharp decrease in density (**Supplementary Fig. 27B**). We made similar observations with a larger-scale scRNA-seq dataset of the same mouse model (**Supplementary Fig. 27C**)^9^, consistent with previous studies which have demonstrated that metastases are seeded by small group of cells ^50^.

While diffusion maps provide desirable properties for state representation from single-cell data, Mellon’s effectiveness is not tied to their specific properties. Mellon is capable of estimating densities in any representation with a meaningful distance metric. To demonstrate this, we used Mellon to infer cell-state densities from a MIRA representation^51^ of a multimodal dataset of skin differentiation (**Supplementary Fig. 27D**). Similar to observations with single modality datasets, we observed extensive variability in densities with low-density regions corresponding to exit from the stem-cell state and specification of different lineages (**Supplementary Fig. 27D**).

We also tested Mellon’s ability to recover chromatin-state densities using single-cell histone modification data. We applied Mellon to compute densities using a single-cell sortChIC dataset of histone modifications in mouse hematopoiesis^48^ using H3K4me1, a histone modification that marks enhancers and H3K9me3, that marks heterochromatin (**Fig. 5A-D**). H3K4me1 densities demonstrated extensive heterogeneity similar to single-cell RNA and ATAC (**Fig. 5E**). On the other hand, H3K9me3 densities are relatively uniform and lower compared to H3K4me1 (**Fig. 5F**). While the lower density is likely reflective of the noise in heterochromatin marks which tend to occur in broad megabase size domains, the relative uniformity is likely reflective of the underlying biology: Active chromatin marks like H3K4me1 accurately have been shown to distinguish cell types and states whereas heterochromatin mark H3K9me3 struggles to achieve the same resolution^48^. This follows the function of H3K9me3 to aid in general repression of other cell-fates rather than to actively establish cell-type identity^48^. To quantify the difference in heterogeneity between the two marks, we subsampled cells from each mark and compared the rank of the covariance matrices (**Fig. 5D**). The covariance rank is a measure of information content where greater the rank, higher is the complexity of the system. The distribution of ranks is significantly higher for H3K4me1 compared to H3K9me3 (p-value < 1e-30, Wilcoxon rank-sum test) demonstrating a greater complexity across cell-types for the H3K4me1 histone modification.

**Figure 5:**
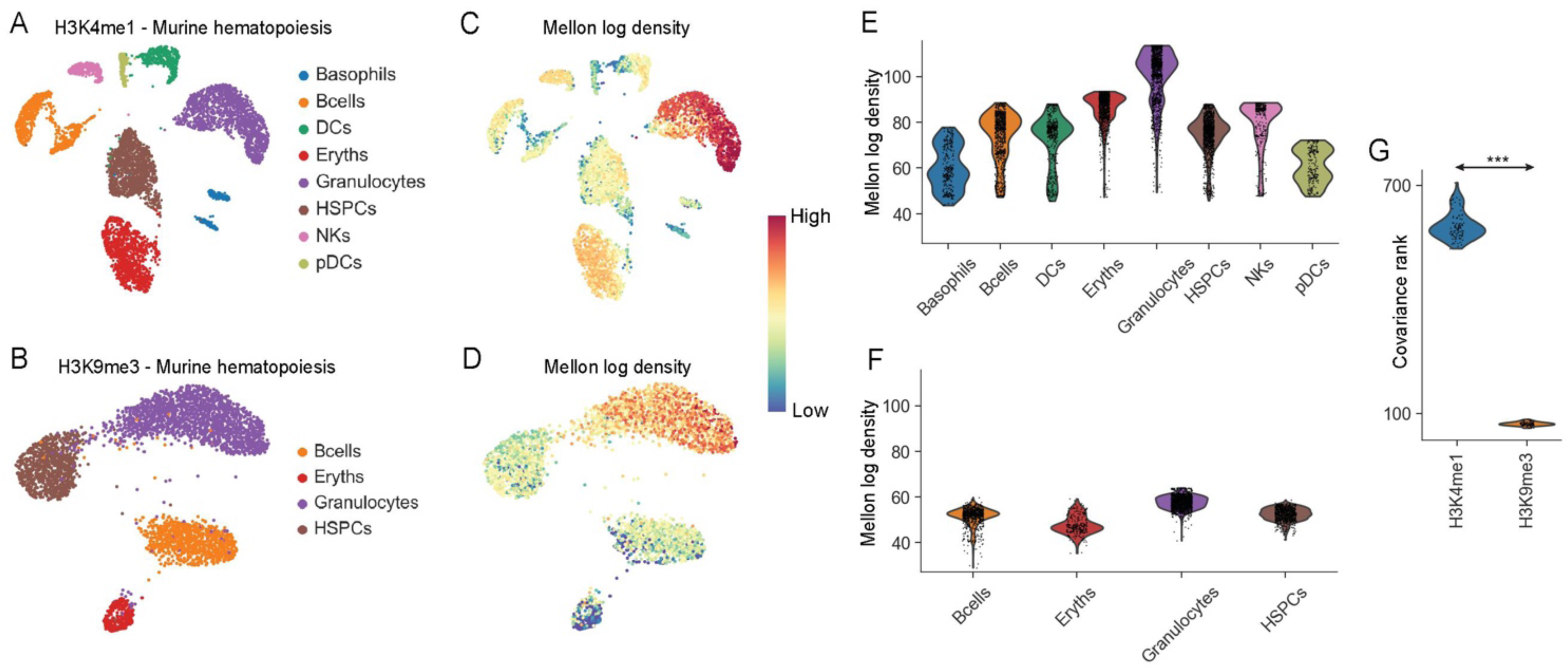
Application of Mellon density estimation to single-cell chromatin data modalities. A-B. UMAPs of H3K4me1 (A) and H3K9me3 (B) mouse bone marrow sort-ChIC dataset^48^ colored by cell-type. C-D. Same as (A-B), with UMAPs colored by Mellon log density E-F. Violin plots to compare cell-state densities among different hematopoietic cell-types. Top: H3K4me1, Bottom: H3K9me3 G. Violin plot of covariance matrix rank for each sort-ChIC dataset for 100 runs of Mellon by repeatedly subsampling 80% of the dataset. (*** p-value < 1e-30, Wilcoxon rank-sum test)

Our results demonstrate the versatility of Mellon with diverse single-cell data modalities and data representations. Mellon’s ability to robustly and accurate identify cell-state densities from single-cell chromatin data suggests a key utility in mechanistic investigations with emerging technologies that concurrently measure active and repressive modifications^52, 53^.

### Highly efficient and scalable: Mellon’s power in atlas-scale single-cell analysis

There is a growing trend towards generation of atlas-scale datasets that profile millions of cells, as well as integration of smaller datasets into large-scale data repositories^54, 55^. To enable density computation in these massive datasets, Mellon incorporates several features that enable efficient scalability: First, Mellon uses a sparse Gaussian process, leveraging landmark points to approximate the covariance matrix, facilitating the efficient handling of high-dimensional data, and reducing the computational overhead associated with large datasets. Second, Mellon requires a single computation of the covariance matrix, removing the need for continuous updates in every iteration and thus improving computational efficiency. Finally, Mellon is built on the JAX python library, which is well-known for its high-performance computing capabilities^56^. The utilization of JAX allows Mellon to optimize available hardware resources, further enhancing its scalability and computational efficiency.

Mellon’s architecture is designed to scale near linearly in time and memory requirements i.e., the runtime grows proportionally with the number of cells when the number of landmarks is kept constant (**Fig. 6**). To demonstrate the scalability, we used the T-cell depleted bone marrow (8.6k cells)^21^, CD34+ bone marrow (6.8k cells)^21^, mouse gastrulation (116k)^44^ and the iPS reprogramming dataset (250k cells)^8^ spanning datasets of different sizes and characteristics. For example, using a single CPU core and a default of 5000 landmarks, Mellon required only 100 seconds to process 10k cells and 17 minutes to process 100k cells of the iPS dataset, including the computation of diffusion maps (**Fig. 6A**). Additionally, Mellon benefits from parallelization, enabling even faster processing times (**Fig.6B**). This highlights Mellon’s efficiency in handling datasets of different sizes.

**Figure 6:**
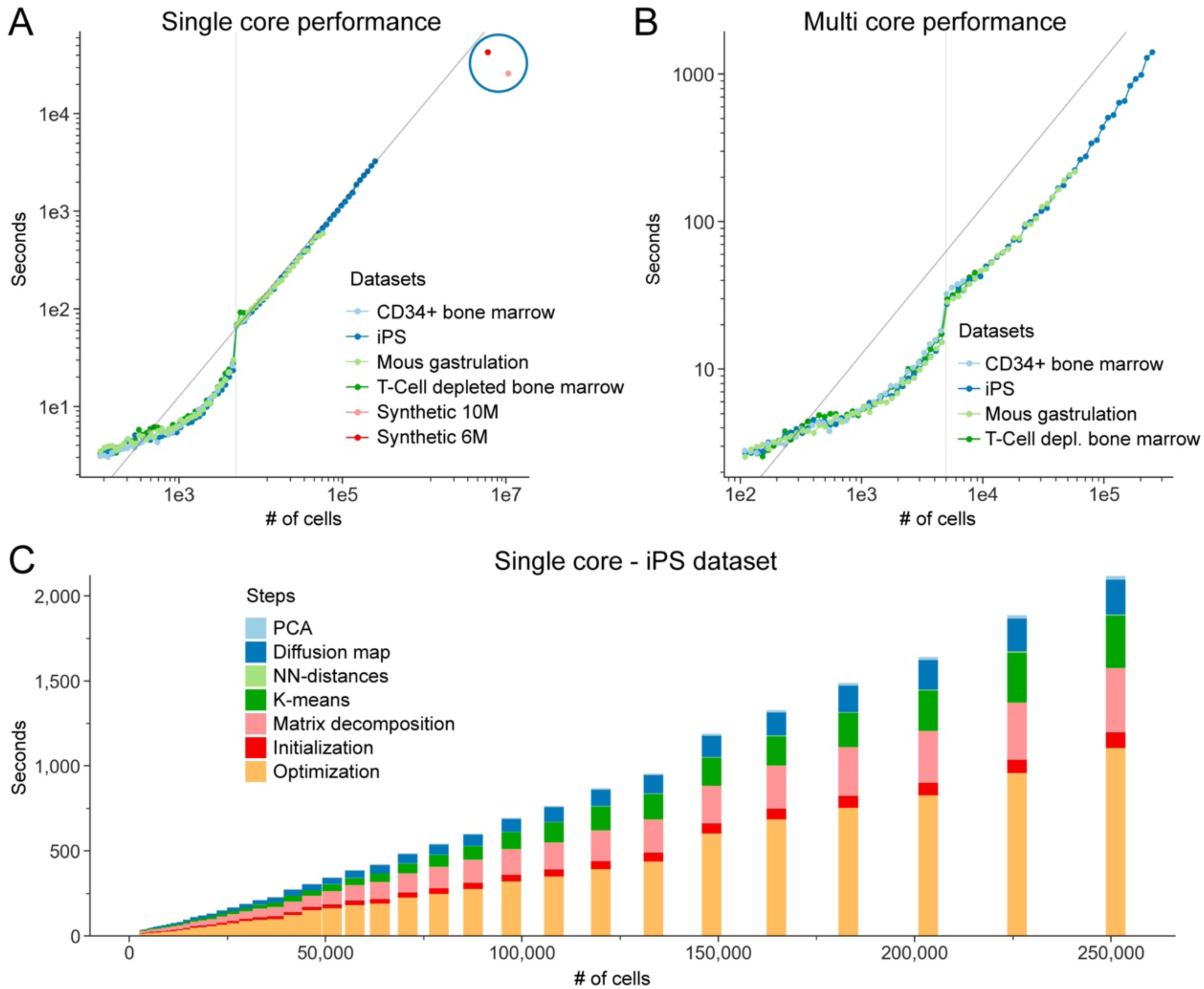
Performance benchmarking of Mellon for demonstrating its scalability and linear time complexity. A. Demonstrates the CPU time required for Mellon’s density inference on a single core across various dataset sizes from four distinct datasets. Each dataset is successively downsized by randomly removing 10% of cells. The data points in this log-log plot align closely with the diagonal line that has a slope of 1, indicating a linear relationship between the number of cells and the CPU time required, which suggests a linear time complexity of Mellon’s algorithm, particularly for large datasets. Notably, statistics for the two large synthetic datasets (6 million and 10 million cells), marked by a blue circle, fall below the diagonal. This emphasizes that a nonlinear increase in compute time does not dominate, even for these larger datasets. For these two synthetic datasets, the computation of diffusion components was omitted, and the larger dataset (10 million cells) uses only 1,000 landmarks, instead of the usual 5,000. The vertical line at 5,000 cells marks the point where the Gaussian process changes from a full process to a sparse one, demonstrating how Mellon adapts to larger datasets by computing the density based on a subset of ’landmark’ cell states. B. Same as (A) but using 36 CPU cores, showcasing the computational efficiency achieved through parallel processing. The data points, situated below the slope-1 diagonal, represent a decrease in CPU time due to the parallelization of tasks. C. Breakdown of the total single-core CPU time for the iPS dataset into individual computational stages, offering insights into the contribution of each stage to the overall density inference process.

To further evaluate Mellon’s scalability, we performed benchmarking on simulated datasets, where we utilized a single CPU core and excluded the time for diffusion map computation. Mellon demonstrated its capability to handle large-scale datasets by accurately computing densities on a dataset of ∼6 million simulated cells in less than 12 hours (**Supplementary Fig. 28A-D**). In addition, we investigated Mellon’s scalability with reduced numbers of landmarks. When the number of landmarks was decreased to 1000, Mellon maintained its accuracy while requiring less than 7 hours to accurately estimate densities for a simulated dataset of around 10 million cells (**Supplementary Fig. 28E-H**). These results highlight Mellon’s remarkable scalability to tackle the burgeoning demands of increasingly large single-cell datasets.

## Discussion

Rapid transcriptional changes that lead to rare transitory cells and thus induce differences in cell-state density have been well-documented as a fundamental property of developmental systems from plants to mammals^5^. Single-cell studies have reinforced the critical nature of rare transitory cells in diverse biological contexts such as development^42, 44^, differentiation^1^, reprogramming^8^, plasticity of tumors^10^ and metastasis^9^. However, existing approaches for estimating cell-state densities have fundamental limitations: They either rely on noisy neighborhood-based estimates around individual cells or utilize 2D dimensional embeddings that do not capture the full complexity of cell-states. Mellon addresses this gap by providing a robust and accurate framework for estimating cell-state densities from high-dimensional cell-state representations. Mellon can be applied to dissect the density landscapes not only in differentiation and development but also during reprogramming, regeneration, and disease. We extended the Mellon framework to estimate time-continuous cell-state density for temporal interpolation of time-series data. The computational efficiency of Mellon allows for rapid density computations, enabling the analysis of large-scale single-cell datasets containing hundreds of thousands of cells within minutes. Furthermore, Mellon’s flexibility supports density estimation for diverse single-cell data modalities, making it a versatile tool for investigating cell-state densities across various biological systems.

Mellon’s innovative approach involves formalizing the connection between density and nearest neighbor distances using a Poisson process and establishing a link between cell-state similarity and density through Gaussian processes. This unique combination overcomes computational challenges in high-dimensional spaces and enhances the robustness and accuracy of density estimation. The scalability of Mellon is achieved through the utilization of sparse Gaussian processes, heuristic for length-scale optimization to avoid redundant computations of the covariance matrix, and implementation using efficient JAX libraries.

Our work underscores the significance of cell-state density in understanding differentiation trajectories and the potential of Mellon to provide new insights into the regulatory mechanisms guiding cell-fate decisions. We have demonstrated the effectiveness of Mellon in estimating cell-state density using hematopoietic differentiation. Mellon’s ability to capture the heterogenous density landscapes, where high-density regions correspond to major cell-types and low-density regions represent rare transitory cells, is particularly evident. By incorporating our trajectory detection algorithm, Palantir, we have been able to observe a correlation between low-density regions and lineage specification. Further, our gene change analysis procedure helps identify gene expression changes that drive low-density transitions and can help elucidate the underlying molecular mechanisms. This was particularly insightful during our investigation into B-cell fate specification, where the detection of low-density regions played a central role in identifying the importance of enhancer priming and characterizing the regulation of the master regulator EBF1. The pattern of alternating high- and low-density regions, observed during the process of B-cell development, further highlighted the dynamic nature of differentiation. Importantly, the consistency of our density estimates across independent donor samples highlights reproducibility and reliability of Mellon. Therefore, these findings provide a strong foundation for further exploration into the intricacies of cellular differentiation.

An important consideration for estimation of cell-state density is the inherent dimensionality of the cell-state space. Mellon by default uses the dimensionality of the cell-state space i.e., number of diffusion components for density estimation, but the intrinsic dimensionality is likely substantially lower. In other words, not all diffusion components are relevant to describe any given region or point in the cell-state space. With measures of intrinsic dimensionality^57^, one can produce explicit units of density and make statements about how many more cells per volume can be expected at a given state. High-dimensionality of the state-space also presents a challenge for automatic determination of high- and low-density regions. Therefore, we compared densities with lower-dimensional projections such as pseudo-time to identify such regions. Incorporation of density as a feature for clustering algorithms or the use of local context density could lead to direct computation of such regions in the high-dimensional state-space.

We anticipate that the time-continuous cell-state densities for temporal interpolation will be a powerful addition to the computational toolkit for modeling cell-state dynamics using time-series single-cell datasets. Mellon provides capabilities to interpolate cell-state density and since the density function is differentiable, it also supports the computation of density change at all times between measured time points. Thus Mellon densities can serve as inputs for development of computational algorithms leveraging advances in optimal transport^43^ for a high-resolution characterization of cell-fate choices using time-series data.

We have demonstrated that cell-state density is a fundamental property of the differentiation landscape by observing that homeostatic density is re-established upon lung regeneration (**Fig. 2I-J**). Thus, the Mellon cell-state density function can itself serve as a phenotype of that differentiation landscape that is altered upon perturbation. Single-cell datasets in unperturbed and perturbed conditions can be jointly embedded into a common state space such as diffusion maps and density functions can be computed separately for each condition in the common space. Comparison of densities from different conditions can not only provide estimates of differential abundance at unprecedented resolution but can also be utilized to develop summary statistics that describe and quantify the nature of the perturbation across the entire differentiation landscape.

Fundamentally, the density function estimated by Mellon provides a comprehensive description of the differentiation landscape, representing the probability distribution of cells within different states. Unlike many existing approaches that rely solely on the measured cell-states and number of cells, which can introduce technical biases and impact the interpretation, Mellon’s density function reflects the inherent complexity of the biological system. As the number of measured cells increases, the density function converges in complexity, allowing for a more accurate representation of the relative abundances of all possible cell states. Mellon can be extended to support online learning, enabling the incremental refinement of the density function with new data. Monte Carlo sampling approaches can leverage Mellon’s cell-state density function to generate synthetic cell-state data, which can greatly enhance data-intensive machine learning models. By incorporating the richness of the density function, these synthetic data can augment training sets and improve the performance and robustness of downstream analyses. Further, the differentiability of Mellon’s density function opens up possibilities for utilization of partial differential equations. This enables the modeling of the differentiation process as a dynamical system and facilitates the inference of regulatory dynamics underlying cellular transitions. By integrating Mellon’s density function within differential equation frameworks, one can gain deeper insights into the regulatory mechanisms governing cellular differentiation and uncover key factors driving the dynamic processes.

## Data Availability

All datasets used in the manuscript have been previously published and the accession numbers are listed in **Supplementary Table 1**. Mellon results and cell-type metadata information for the T-cell depleted bone marrow and the mouse gastrulation data are available on Zenodo at https://doi.org/10.5281/zenodo.8118722.

## Code Availability

Mellon is available as a Python module at https://github.com/settylab/Mellon. Jupyter notebooks detailing the usage of Mellon including cell-state density estimation, gene change computation, time-continuous cell-state density estimation, and enhancer classification are available at https://mellon.readthedocs.io/en/latest/. Pipelines for running SEACells, computing gene-peak correlations, primed and lineage-specific accessibility scores are available at https://github.com/settylab/atac_metacell_utilities.

## Author Contributions

D. O. and M. S. conceived and designed the study, developed Mellon, developed additional analysis methods and statistical tests. D. O. and B. D. developed the heuristics, performed robustness analyses and implemented the framework. C. J. and M. S. performed analysis of enhancer dynamics. C. D. supported enhancer dynamics analysis. D. O., C. J. and M. S. wrote the manuscript.

## Acknowledgements

We thank members of the Setty lab for discussions and comments on the manuscript. This study was supported by National Institute of General Medical Studies grant R35 GM147125 and Brotman Baty Institute Pilot Award to MS; National Institutes of Health grant ORIP S10OD028685 to support high-performance computing at the Fred Hutchinson Cancer Research Center.

## Methods

### Mellon Algorithm

Mellon is a computational tool designed to infer cell-state densities from high-dimensional single-cell data. The objective of Mellon is to characterize the complex density landscapes of single-cell data (**Fig. 1A-B**) with density estimates that are robust even in low-density regions, while maintaining computational efficiency.

Mellon’s computational model is grounded on two core assumptions. Firstly, within the chosen representation of cell states, smaller distances between cell states signify higher biological similarity. In other words, we assume that biological dissimilarity can be effectively quantified by the Euclidean distance within this representation. Secondly, we assume that cell-to-cell density changes are smooth and continuous, meaning that cell states of high similarity are expected to have similar state densities.

The input to Mellon is a high-dimensional representation of the cell-states (e.g., Diffusion maps). The Euclidean distance between these cell-states serves as a measure of biological dissimilarity. Mellon outputs a *continuous density function* that allows evaluation of cell-state densities at single-cell resolution (**Fig. 1E-F**). The densities are computed in the high-dimensional cell-state space and visualized using low-dimensional embedding techniques such as UMAPs.

The Mellon framework contains the following major components:

- Mellon first calculates the distance to the nearest neighbor for each cell in the cell-state space, following the first assumption.
- The distances are linked to density via the *Nearest-Neighbor Distribution (***Fig. 1C**).
- Densities between highly related cell-states are connected by the *Gaussian Process* and the associated kernel function (**Fig. 1C-D**).
- A *Bayesian Model* (**Fig. 1D**) is deployed, integrating the nearest-neighbor distribution, kernel function, and Gaussian Process to compute the continuous cell-state density function (**Fig. 1E**).

We next describe each of these components in detail along with our approach to scale Mellon for large datasets.

#### Nearest-Neighbor Distribution

The core principle of Mellon relies on the relationship between nearest-neighbor distances and density, as depicted in **Supplementary Figure 3**. This connection can be formalized using a Poisson point process to define a nearest-neighbor *distribution*, which describes the probability of another cell-state existing within some distance of a reference cell-state. Intuitively, regions with a higher density of cell-states correspond to tighter nearest-neighbor distributions, while low-density regions result in broader distributions (**Fig. 1C**).

For distance 𝑟 and density 𝜌, the probability density function of the Nearest-Neighbor distribution 𝑓_NN_: ℝ^d^ → ℝ^+^ is given by

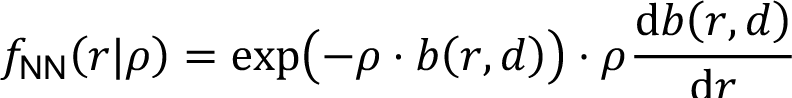

where 𝑏(𝑟, 𝑑) is the volume of a 𝑑-dimensional ball with radius 𝑟. For a cell-state 𝑥 ∈ ℝ^d^ with a nearest neighbor distance dn(𝑥), this probability density function gives rise to the following maximum likelihood estimate for density if no prior is employed, formalizing the inverse relationship between nearest neighbor distances and density as

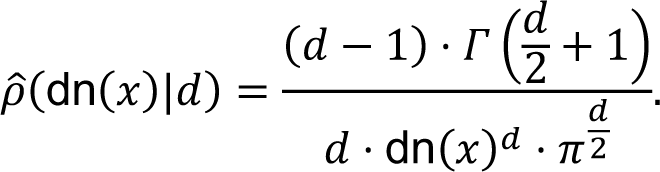

The derivation is detailed in **Supplementary Note 1**. The use of Poisson point process is facilitated by the two key assumptions of Mellon: The use of Euclidean distance in the cell-state space is a critical requirement for defining the probability density function. The second is the assumption of smoothness in cell-to-cell density changes. This is crucial as it allows us to assume that the density at a given cell-state corresponds to the average density within a sphere centered at that state, with the radius of the sphere defined by the nearest neighbor distance.

#### Gaussian Process

Building upon the foundational connection between nearest neighbor distance and density, Mellon utilizes Gaussian process (GP) priors to establish a relation between the densities of highly-similar cell-states, facilitating a continuous density function estimation. Similarity between cell-states is encoded using the covariance function of the Gaussian process. The random variable of the GP, denoted as 𝑓(𝑥), serves as the approximation of the logarithm of the cell-state density. Two properties of GPs make them ideally suitable for cell-state density estimation from single-cell data: (i) GPs can be used to describe arbitrarily complex functional spaces where the true functional form is unknown and (ii) GPs provide robust estimates even when small number of observations are available.

The GP is defined as follows:

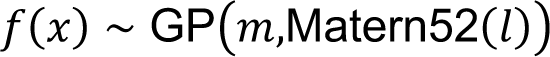

where 𝑚 and Matern52(𝑙) are the mean function and the Matern covariance function respectively. A more detailed assessment is provided in **Supplementary Note 2**.

#### Mean function

The true log-cell-state density approaches negative infinity away from any observed cell state. However, functions sampled by the Gaussian process approach the chosen mean. To approximate the true behavior of density functions we choose a very small value for the mean 𝑚 that implies a vanishingly small probability for a distant cell state. This mean function is given by the constant:

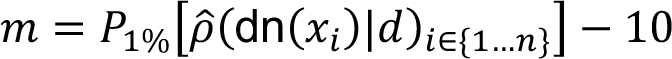

where 𝑃_1%_[⋅] is the 1^st^ percentile of the given data, 𝜌8 is the heuristic maximum likelihood estimate for density, and dn(𝑥_’_) is Nearest-Neighbor Distance of cell-state 𝑥_’_ in ℝ^#^. The choice of this mean is discussed in **Supplementary Note 3**.

#### Covariance Function and length scale

Similarities between cell-states are encoded through the GP covariance function or kernel. Specifically, the kernel function defines the covariate structure between cell-states which translates to the smoothness of the density function. Some commonly used kernels are arbitrarily smooth and allow arbitrary differentiability. Assuming such smoothness can, however, lead to unrealistic results^19^. We therefore chose to use the Matern covariance function with 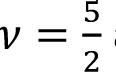 as the kernel, which is exactly twice differentiable and thus constrains the degree of smoothness of the density function. The Matern52 kernel for a pair of cell states 𝑥, 𝑦 ∈ ℝ^d^ is defined as:

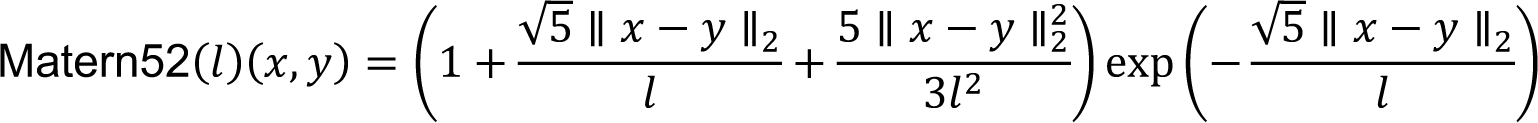

The covariate structure of the Gaussian process is governed by the length scale parameter, denoted as 𝑙, which essentially determines the radius of influence around each cell state. Conceptually, the length scale sets the reach of influence for each cell, defining the range within which other cells contribute to the local density estimate (**Supplementary Fig. 3**). In areas of lower density, fewer but more representative cells influence the density estimate, while in higher density areas, a larger number of cells contribute. This scenario gives rise to an effective number of neighbors that is density-dependent, which is a direct result of the distance-mediated impact on the local density estimate (**Supplementary Fig. 3**).

This method not only increases the reliability and robustness of density estimates, but it also enables the creation of a continuously changing density function between cell states, offering a nuanced representation of biological phenomena. Unlike the k-nearest neighbor methods for density estimation that assign an equal weight to all k neighbors irrespective of their distances, the continuous covariance function of the Gaussian process accounts for the distance between cells, smoothly adjusting the weight of their contribution. The resulting impact on the local density estimate facilitates a more precise representation of the cell-state landscape.

The ideal length scale strongly depends on the availability of data at different points of the cell-state space and encompasses a specific amount of cells needed to support a reliable density estimate of a given state. We therefore derived a heuristic for length scale as function of the mean nearest neighbor distance between cells:

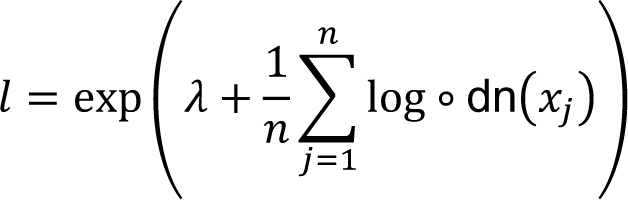

Here, 𝜆 = 3 is a heuristic value inferred from an extensive cross-analysis of multiple datasets. The derivation of the length-scale heuristic is described in **Supplementary Note 4**.

#### Sparse Gaussian Process

Gaussian process computation usually necessitates 𝑂(𝑛^0^) operations, where 𝑛 is the number of cells and thus can be prohibitively expensive for large datasets. To address this computational challenge, Mellon utilizes a sparse approximation of the GP. This approach substantially reduces the computational complexity while maintaining the versatility and expressiveness of the full GP model.

The sparse GP in Mellon is constructed using a subset of data points, referred to as “landmark cell-states,” that essentially act as inducing points. These landmark states are chosen to capture the essential structure of the cell-state space, providing a representative skeleton for the full GP model. This sparse GP approach translates to an efficient 𝑂(𝑛𝑘^$^) time complexity for inference, where 𝑘 is the number of landmark points, a substantial reduction from the cubic time complexity of the full GP.

The specifics of the sparse GP implementation play a crucial role in the overall performance of the Mellon and are described in **Supplementary Note 2.**

##### Landmark selection

The choice of landmarks, akin to the “inducing points” in a Gaussian process, is essential to ensure precise recovery of the approximated covariance structure. Previous studies have demonstrated that k-means centroids are well suited for this purpose^58^. We assessed the accuracy of this approach by comparing the inferred density derived from the landmarks against the density function inferred from a non-sparse, or “no-landmarks” version (**Supplementary Fig. 15**). This comparison showed a convergence of the landmark-based model towards the non-sparse version, thereby confirming the efficacy of the landmark selection. We therefore use k-means clustering as the default landmark selection in our algorithm and initialize it with kmeans++^59^ to ensure computational efficiency.

#### Full Bayesian Model

The full Bayesian model used in the Mellon algorithm is formally defined as follows:

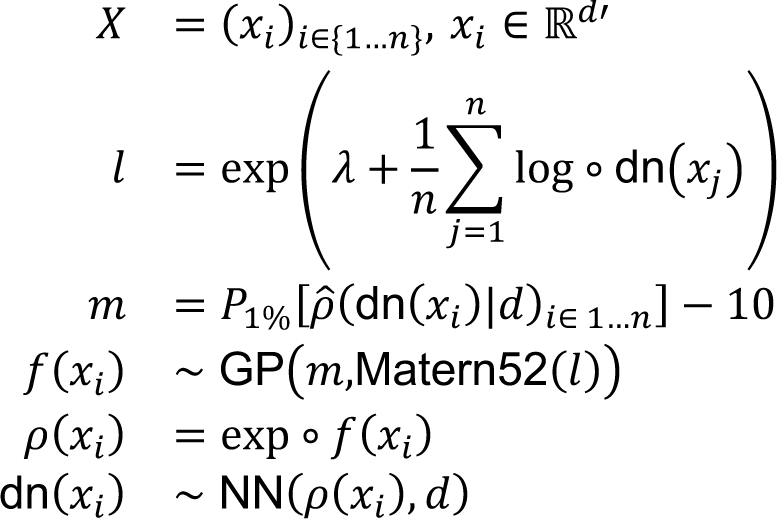

Where

- 𝑋 represents the cell states, where each cell state, 𝑥_’_, is a vector in the 𝑑′-dimensional Euclidean space. The cell states form the primary input data for the model.
- · 𝑙 is the length scale of the GP covariance function. 𝑙 is calculated from the distances to the nearest neighbors in the cell-state space. The logarithm of these distances is averaged and added to a fixed parameter 𝜆 = 3. The sum is then exponentiated to produce the length scale.
- · 𝜌8 is the heuristic maximum likelihood estimate for the density.
- _<συβ>·_ dn(𝑥_’_) is Nearest-Neighbor Distance of cell state 𝑥_’_
- · 𝑚 is the GP mean function. 𝑚 is calculated as the 1% percentile of the heuristic maximum likelihood estimates of density subtracted by a constant (10 in this case). This mean function represents the average behavior of the underlying cell-state densities.
- · Matern52 is the Matern covariance function with 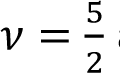 and length scale 𝑙.
- · 𝑓(𝑥_’_) is a random function generated by a sparse Gaussian process (GP), where 𝑥_’_ is the input cell-state vector. The GP is defined by the mean function 𝑚 and the Matern covariance function.

The cell-state density function 𝜌(𝑥_’_) is the random variable of interest and is calculated by exponentiating the function 𝑓(𝑥_’_). This ensures that the density is always positive. The final part of the model is the Nearest Neighbor Distance distribution NN of the Nearest Neighbor distance dn(𝑥_’_), which is calculated as a function of the cell-state density 𝜌(𝑥_’_) and the dimensionality 𝑑.

#### Initialization

An appropriate initialization 𝑦′ for the density function 𝑦 = 𝜌(𝑥_’_) can improve the convergence of the maximum a posterior estimation. We employ a regression approach to initialize density estimation using the heuristic maximum likelihood estimates of the log-density 𝜌8 (**Supplementary Note 1)**:

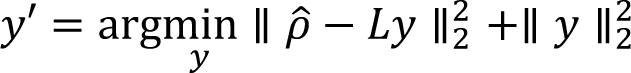

Where 𝐿 represents the transformation matrix within the Gaussian process, facilitating the conversion of the latent representation, 𝑦, into the log-density function. High values in 𝑦 are penalized through a ridge regression to simulate the additional smoothness of the true density over 𝜌8.

#### Density at single-cell resolution

The log-density function 𝑓(𝑥) is evaluated at each single cell 𝑥 ∈ ℝ^#d′^, to estimate log cell-state density at single-cell resolution. The estimated densities in cell-state space are visualized using techniques such as UMAPs for convenience. Note that the density function can be evaluated at any point in the cell-state space including states that are not measured in the dataset. Single-cell densities can be examined along pseudo-time, individual diffusion components or between interconnected clusters to identify high- and low-density regions.

#### Note on the number of landmarks for sparse Gaussian Process

The number of landmarks serves as a parameter to the sparse Gaussian Process within Mellon. It’s crucial that the number of landmarks is sufficiently large to accurately capture the intricate patterns and variability within the cell state density function. However, it is important to consider the trade-off involved: an increased number of landmarks enhances the model’s capacity to encapsulate finer details, but it also increases the computational demands.

Mellon employs a default selection of 5,000 landmarks, an empirical decision grounded in extensive testing with a multitude of datasets with different properties (**Supplementary Table 1**). Our evaluation underscores the robustness of Mellon’s density estimates across all investigated datasets, consistently demonstrating stability even when the number of landmarks is substantially altered (**Supplementary Fig. 15**).

Nevertheless, the optimal number of landmarks can be contingent on the complexity and volume of the particular dataset under examination. To assist users in selecting a representative number of landmarks, Mellon incorporates a test for approximating the rank of the covariance matrix. Should the complexity of the function appear exhausted using the existing landmark skeleton, a warning will be issued. This serves as an indication that the selected number of landmarks might be insufficient for the model to accurately capture the density function of the cell-states.

### Scalability of Mellon

The implementation of Mellon leverages modern advances in numerical computation libraries, specifically the JAX library, to enable efficient calculations and seamless differentiation. JAX is particularly suited for our purposes due to its unique capability of just-in-time (JIT) compilation using XLA (Accelerated Linear Algebra), a linear algebra compiler developed by Google^56^. This feature ensures efficient utilization of hardware resources, especially for large-scale computations and vectorized operations, which are intrinsic to our method.

Mellon’s scalability to large single-cell datasets is ensured through the use of a Sparse Gaussian Process (GP). The sparse GP allows us to approximate the full GP model, significantly reducing the computational demands while retaining the essence of GP’s expressiveness. This scalability (**Fig. 6**) makes Mellon practical for atlas-scale single-cell data sets, which often involve millions of cells.

Finally, model tractability in Mellon is achieved through the adoption of a length-scale heuristic for the GP covariance function. The covariance function is crucial in a Gaussian Process as it dictates how many nearby points in the input space influence each other in the output space. Typically, the length scale of this function is subject to inference or optimization, often involving computationally intensive iterative processes that require repeated updates of the covariance matrix and its Cholesky decomposition. In Mellon, we sidestep this computational demand by deriving an appropriate length scale with a data driven approach designed to adapt to the varying local densities in the high-dimensional cell-state space (**Supplementary Note 4**). This not only streamlines the computation but also assists in avoiding overfitting to dense regions, resulting in a smooth and accurate portrayal of cell-state density relationships.

Together, these components create a balance between computational efficiency and model expressiveness, making Mellon an effective and practical tool for cell-state density estimation from large, high-dimensional single-cell data.

#### Inference

Mellon, by default, employs the L-BFGS-B optimization algorithm to infer the maximum a posteriori (MAP) estimates of the posterior likelihood. Notably, our implementation provides direct access to the posterior distribution of the density function. This is realized through a JAX function with automatic differentiation, thus facilitating the use of any preferred inference scheme while retaining computational simplicity.

This flexibility is crucial because, in Bayesian inference, the MAP estimate can be subject to the transformation of the latent representation and might not necessarily represent the “true” underlying cell-state density. In fact, empirical evidence (**Supplementary Fig. 30)** indicates that the MAP estimate strongly coincides with the posterior mean. However, without a definitive ground truth, it is challenging to ascertain which estimate more closely resembles the true cell-state density.

In essence, Mellon’s versatile implementation provides a robust framework for density estimation that can adapt to diverse inference schemes, offering users the freedom to employ the technique best suited to their specific study.

### Cell-state Representation

Mellon utilizes diffusion components^11^, as implemented in Palantir^14^, as the representation of cell-state space. Diffusion maps have been widely used in single-cell data analysis owing to their reliable and robust representation of cell-states^12, 14^. Cellular states in phenotypic landscapes reside in substantially lower dimensions compared to measured gene expression owing to gene regulatory networks inducing a strong covariate structure amongst genes. Therefore, biological similarity between cell states is more closely linked to the distance they can traverse along the phenotypic landscape, rather than solely their direct proximity in gene expression space. Diffusion maps identify the intrinsic structure in single-cell data, mitigating noise by treating the data as realizations of a stochastic process. They not only efficiently reduce noise in single-cell data but also extract a faithful representation of the underlying cell-state manifold.

Further, the distances computed using diffusion maps, termed diffusion distances, are a measure that reflects the interconnectedness of data points along the phenotypic manifold. Importantly, diffusion distance operates along this manifold, which is constructed from the observed cell states, thereby providing a meaningful indicator of biological similarity between cells. Therefore, the use of diffusion distance in the estimation of cell-state density leads to a biologically relevant quantification of cells sharing a similar state.

Diffusion maps can be constructed for different single-cell data modalities with appropriate preprocessing. We recommend the use of PCA for RNA and SVD for ATAC and histone modification data. “Data preprocessing” section provides more details on preprocessing of single-cell datasets. Diffusion maps can also be constructed using other latent representations^36^ or multimodal representations ^51^.

#### Number of Diffusion Components

The dimensionality of the subspace where the data is represented is determined by the number of diffusion components utilized in Mellon. Mellon results are robust to the number of diffusion components indicating that pinpoint precision in their selection isn’t strictly necessary (**Supplementary Fig. 13**). However, some considerations ae important while choosing the number of diffusion components: Selecting a high number of diffusion components might lead to the inclusion of unnecessary noise within the state representation, reducing the granularity of the resulting density model. Conversely, choosing a small number of diffusion components might under-represent the complexity of the data, thus also leading to a less detailed density model. The optimal number of diffusion components is therefore largely data-specific and should be chosen to best capture the inherent structure and complexity of the cell data, without unnecessarily increasing noise or forfeiting essential information. For example, Eigen gap statistic has been previously employed to choose the number of diffusion components^14^.

### Genes Driving Low-Density Cell-State Transitions

Low-density regions representing rare transitory cells are critical for diverse biological processes. We devised a gene change analysis procedure to identify genes that drive cell-state transitions in low-density regions and thus can be used to describe the dynamic behavior of the biological system. The input is a relevant set of cells 𝑆 ⊂ {1, …, 𝑛}, such as those representing a transition of interest. These could include a branch in the cell-differentiation landscape or clusters interconnected by transitory cells. The output is a ranking of genes ordered by their change scores representing their association with the low-density regions in the selected set of cells. Top genes in this ranking can be interpreted as driving the transitions in low-density regions.

We first compute a measure of local variability of a gene for each cell-state: We compute the expression change from a cell when transitioning to each of its neighbors and normalize the change by the distance between the cells in state space to account for the magnitude of the state transition. The maximal normalized change amongst the neighbors of the cell is nominated as the local variability of the gene for the corresponding state. Formally, the local variability for gene 𝑗 in cell-state 𝑖 is defined as:

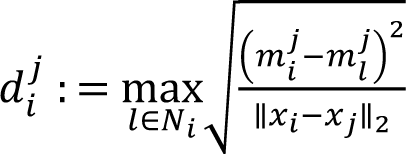

where, *m^j^_i_* denotes the MAGIC imputed expression of gene 𝑗 in cell 𝑖, 𝑁_’_ is the set of 𝑘 nearest neighbors of cell 𝑖.

We next compute, a low-density change score 𝑠. for each gene 𝑗, as the sum of the gene change rates *d^j^_i_* across the selected cells, inversely weighted by the cell-state densities, 𝜌(𝑥_’_):

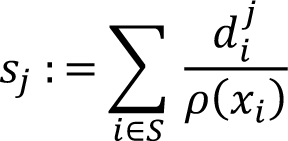

This scoring approach encapsulates the hypothesis that genes with high change score in low cell-state density regions may be driving transitions. Genes are ordered by the change score and genes with scores > 95^th^ percentile are considered to be driving low-density changes (**Supplementary Fig. 7**).

### Primed and lineage-specific accessibility scores from scATAC-seq data

Gene scores from scATAC-seq are typically computed by summarizing the accessibility of peaks in the body of the gene and its vicinity^36^. This, however, does not consider the history and temporal dynamics of peak accessibility. Enhancer priming, where open chromatin peaks are pre-established in stem cells without turning on gene expression but maintain the gene locus in an open state for lineage-specific upregulation, is an important mechanism through which stem cells encode high differentiation potential^32-^

^34^. To investigate the establishment of peak accessibility, we devised a procedure to disentangle primed and lineage-specific peaks in the context of cell-fate specification. As a result of the sparsity and noise of scATAC-seq data, our approach utilizes several abstractions and consists of the following steps:

1. Identification of peaks with accessibility strongly correlated with gene expression at metacell resolution
2. Determination of peaks with higher accessibility in the lineage under consideration compared to other lineages using differential accessibility testing between metacells
3. Classification of peaks as primed or lineage-specific based on accessibility patterns in stem cells
4. Determination of primed and lineage-specific accessibility scores for each gene at single-cell resolution.

We developed this approach to identify primed and lineage-specific peaks in the transition from hematopoietic stem cells (HSCs) to B-cell fate committed cells (proB) (**Fig. 3**). We used the monocyte and erythroid lineages as the alternative lineages to test for B-cell lineage specificity.

#### Determination of primed and lineage-specific peaks

##### Metacells and gene-peak correlations using SEACells

We used our SEACells algorithm^21^ to identify metacells from the T-cell depleted bone marrow. SEACells aggregates highly related cells into metacells overcoming the sparsity in single-cell data while retaining heterogeneity. We used the ATAC modality of the multiome data to identify metacells. We used metacells to compare the expression of a gene with the accessibility of each peak in a window of 100kb around the gene to identify the subset of peaks that significantly correlate with expression of the gene (correlation >= 0.1, p-value <= 0.1, Empirical null) (**Supplementary Fig. 19A**).

##### Peaks relevant to particular lineages

Metacells and gene-peak correlations were computed using all hematopoietic lineages in our dataset. We performed differential accessibility analysis to identify the subset of peaks with greater accessibility in the lineage under consideration. We used edgeR^60^ to perform differential accessibility with metacell counts as input. The use of metacells rather than single-cell data for differential accessibility has been demonstrated to provide better sensitivity and specificity^37^. To identify peaks that are relevant to the B-cell lineage, we compared accessibility in pro B-cell metacells and metacells of the erythroid (EryPre1) or monocyte (Monocyte) lineages and retained peaks with the accessibility fold-change log2FC > 0 in either comparison. While this ensures that the selected peaks have greater accessibility compared to other lineages, it does not exclude ubiquitously accessible peaks. We therefore excluded peaks with log2FC < 0.25 in the comparison between stem-cells (HSCs) and erythroid and monocyte lineages.

The resulting set of peaks demonstrate substantially greater accessibility in B-cell lineages compared to all other cell-types (**Supplementary Fig. 19C-D**)

##### Classification of primed and lineage-specific peaks

After identifying peaks with greater accessibility in the B-cell lineage, we assigned primed or lineage-specific status to each peak with a simple logic: A peak is annotated as primed if it is accessible in HSCs and lineage-specific if it is not. Accessibility in HSCs was determined using Poisson statistics as described in SEACells^21^. The mean of the Poisson distribution for a cell-type 𝑐 is estimated using

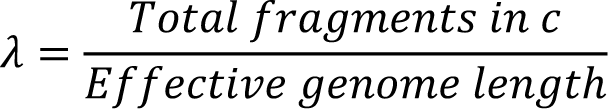

Where 𝑒𝑓𝑓𝑒𝑐𝑡𝑖𝑣𝑒 𝑔𝑒𝑛𝑜𝑚𝑒 𝑙𝑒𝑛𝑔𝑡ℎ is set to 𝑛𝑢𝑚 𝑜𝑓 𝑝𝑒𝑎𝑘𝑠 ∗ 5000. For a peak 𝑝 in cell-type 𝑐 with 𝑛 fragments, 𝜆 is used to estimate the 𝑃 value of observing more than 𝑛 fragments, and 𝑝 is considered open in 𝑐 if 𝑃 < 1𝑒 − 2.

#### Primed and lineage-specific scores

We utilized the primed and lineage-specific peaks to derive primed and lineage-specific scores for the associated genes at single-cell resolution. For each gene 𝑔 and cell 𝑖, the primed accessibility score *s^primed^_ig_* is computed as

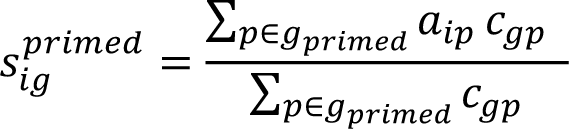

Where 𝑔_primed_ is the set of primed peaks that significantly correlate with gene 𝑔, 𝑎_’>_ is the accessibility of peak 𝑝 in cell 𝑖, and 𝑐_=>_ is the correlation between peak 𝑝 and expression of gene 𝑔 computed using metacells. Therefore, the primed score is a weighted average of the accessibility of primed peaks that correlate with the gene. The lineage-specific score *s^lin^_ig_* is computed in an analogous manner where 𝑔_4’+_ is the set of lineage-specific peaks that significantly correlate with gene 𝑔::

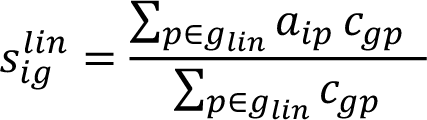

Given the sparsity of the scATAC data, we used imputed peak accessibility for computing scores. The peak counts dataset was TF-IDF normalized^61^ to preferentially weight peaks which are highly accessible in a small proportion of cells. The MAGIC algorithm^2^ was then used to perform imputation using normalized accessibility as the input.

### Data visualization

Accessibility trends along pseudo-time were computed using Mellon. Trends are visualized as a percentage of the maximum value of each trend, to allow for better comparison across genes.

#### Application to T-cell depleted bone marrow data

We applied primed and lineage-specific accessibility scores to characterize commitment of hematopoietic stem cells to B-cells using the T-cell depleted bone marrow multiome data. We used hematopoietic stem cells (HSC), hematopoietic multipotent cells (HMP), common lymphoid progenitor (CLP) and pro B-cells along the B-cell lineage to investigate the open chromatin landscape (**Fig. 3B**). The cells were chosen to span the commitment of stem cells to the B-cell lineage. The high- and low-density regions were manually assigned by comparing pseudotime and log-density of the selected subset of cells (**Fig. 3B**).

#### Primed and lineage-specific accessibility scores in B-cell specification

We applied the SEACells algorithm^21^ to identify metacells using the ATAC modality of the T-cell depleted bone marrow data. Metacells were identified using all cells, resulting in 115 metacells according to recommended heuristic for selecting the number of metacells. Peak accessibility and gene expression correlations were determined using all metacells and the subset of genes with at least 5 peaks were selected for downstream analysis (**Supplementary Fig. 19A**). We computed gene change scores using Mellon using the subset of cells that define B-cell lineage commitment. Genes in the 95^th^ percentile of gene change scores with B-cell specific upregulation in the low-density regions were used to characterize the role of enhancer priming (**Fig. 3)**. Primed and lineage-specific accessibility scores were computed for the subset of these genes with at least one lineage-specific and primed peak each.

#### In silico ChIP

We used in silico ChIP-seq^37^, a recently published approach to identify predicted targets of master regulators of B cell lineage commitment, specifically EBF1 and PAX5. Approaches like FIMO^62^ can determine enrichment scores for TF motifs in ATAC-seq peak sequences but the scores alone are not sufficiently reliable to predict TF targets. In silico ChIP-seq provides a framework for predicting TF targets by using single-cell multiome (scRNA-seq and scATAC-seq) data in addition to motif enrichment by correlating the expression of a TF to the accessibility of a peak. A combination of a high gene-peak correlation and high motif score is more indicative of potential TF binding compared to a peak with only a high motif score^37^. We used our Python adaptation of in silico ChIP-seq using the SEACells metacells as input (<u>github.com/settylab/atac-metacell-utilities</u>). FIMO^62^ was used to associate TF motifs with ATAC-seq peaks, resulting in a peak by TF matrix of scores indicating the strength of match of the TF motif in the peak sequence. In silico TF binding scores are computed as product of correlation between TF expression and peak accessibility and FIMO motif scores as follows:

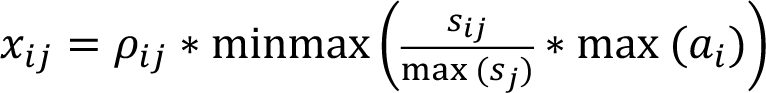

Where 𝑖 is the a ATAC-seq peak and 𝑗 is the a TF of interest, 𝜌_’._is the Spearman rank correlation coefficient of accessibility of 𝑖 and expression of 𝑗 computed across all metacells, 𝑠_’._ is the FIMO motif enrichment score for TF motif 𝑗 binding in sequence of peak 𝑖, max (𝑠.) is the maximum FIMO score for TF 𝑗 across all peaks, and 𝑎_’_ is the maximum accessibility of peak 𝑖 across all cell type metacells.

Minmax normalization is performed as follows:

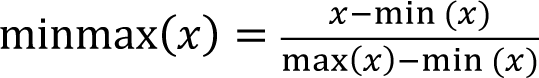

The final in silico ChIP-seq output is a peak by TF matrix, containing a value between -1 and 1 indicating how likely a TF is to bind at a given peak and whether it has a repressive (negative) or activating (positive) effect, or 0 if a peak does not meet the minimum in silico ChIP-seq score (0.15).

#### Regulation of EBF1

Peaks correlated with EBF1 expression were ordered using the procedure outlined in the section “Genes Driving Low-Density Cell-State Transitions” using imputed peak accessibility to compute accessibility change scores instead of gene change scores. In silico-ChIP was to identify the transcription factors with predicted binding sites in the top peak.

### Time-Continuous Density

Time-series single-cell datasets provide snapshots of the changing cell-state densities at discrete time intervals. Our goal is to compute a time-continuous density function to interpolate cell-state densities at any time between the measured timepoints.

We therefore incorporated a time coordinate into the Gaussian process used to generate the log density function and use the covariance of the Gaussian process to link temporally similar cell-states. Effectively the covariance function of time-continuous density has two components: (i) similarity between cells in the cell-state space and (ii) similarity between cells based on their measurement times. Similarity in cell-state space is encoded through the Matern52 kernel with the length-scale parameter as described in **Supplementary Note 5**. We now describe the Matern52 length-scale parameter for the temporal similarity component.

The length scale should be designed such that the covariance between cells from different timepoints reflects the covariance of densities between those timepoints. Therefore, we optimized the length scale to reflect the empirically observed covariance of density functions between different time points. Specifically, we employ Mellon to compute first time-point specific density functions 𝜌_I_ using only the cells from the corresponding time point 𝑡. We next evaluated these functions on *all cells from all timepoints*, and computed a correlation of cell-state density between timepoints:

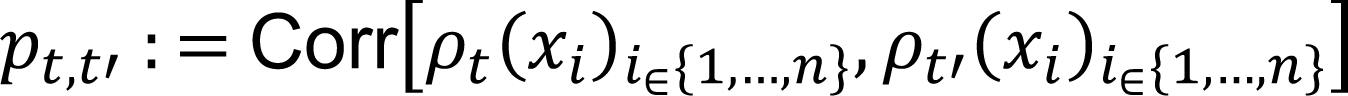

Where 𝑡 and 𝑡′ represent two time points, and Corr[⋅,⋅] denotes the Pearson correlation. This is used to derive a correlation matrix between all measured timepoints 𝑇 (**Supplementary Fig. 23A-D**):

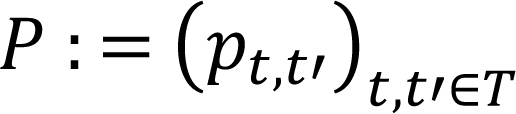

This matrix 𝑃 is then compared to the covariance matrix of time points using the Matern52 kernel. Given the isotropy of the kernel function, it maps a scalar temporal difference 𝑡 − 𝑡′ to a covariance value. The kernel-based covariance matrix is defined as:

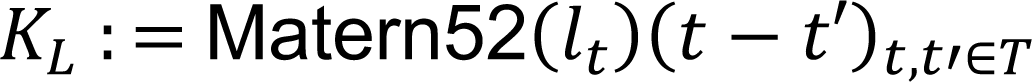

Where 𝑙_I_ is the length scale parameter for the time coordinate. We thus select the 𝑙_I_ by optimizing:

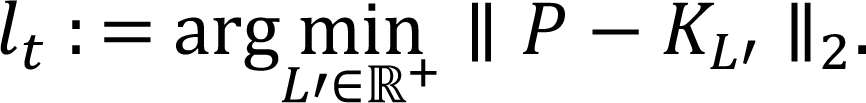

The optimized length scale is used for the Matern52 covariance kernel for the time coordinate, denoted as Matern52(𝑙_I_) (**Supplementary Fig. 23E-F**).

The resulting covariance kernel for cells 𝑖 and 𝑗, situated at their respective states 𝑥_’_, 𝑥., and measurement times 𝑡_’_, 𝑡., is then given as:

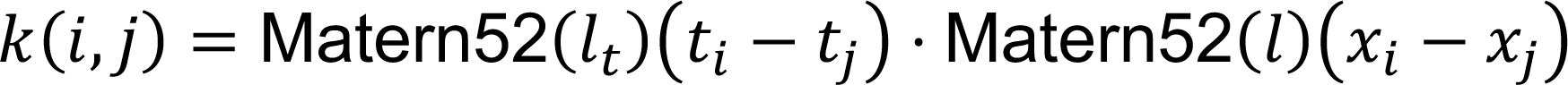

Where Matern52(𝑙) designates the Matern52 covariance kernel for cell-state coordinates and Matern52(𝑙_I_) designates the Matern52 covariance kernel for time coordinates.

This construction is easily implemented with Mellon, since it is designed to support any combination of covariance functions, each operating in distinct active dimensions – in this case, either time or cell-state coordinates.

Using this covariance function, Mellon can compute a continuous density function over time and state space using all samples, and thus can be used to interpolate cell-state densities at unmeasured timepoints. This function is also differentiable in time and state space, and the change in density over time can be determined using the first derivative (**Supplementary Video 1, Figure 25**).

#### Leave-one-out Cross Validation

We validated the effectiveness of the time-continuous density function using a leave-one-out cross-validation strategy (**Supplementary Fig. 24**). We computed a time-continuous density function after excluding cells from a particular timepoint and evaluated the densities at the excluded timepoint using this density function. We compared these densities with a time-agnostic density, which was computed exclusively using cells from the excluded time point and then evaluated across all states. Note that these two density functions were derived from mutually exclusive training datasets.

#### Density along Trajectory

Time-continuous density provides a platform to decipher the dynamics of cell-type proportions and fate choices in true temporal order. As proof of principle, we investigated the cell-type proportion dynamics along the trajectory of a particular lineage. We first used Palantir^14^ to derive fate propensities for all cells and selected the subset of cells with high propensity towards a particular fate.

In the mouse gastrulation data, we applied Palantir using all cells across all timepoints and selected cells which specify the erythroid cells (**Supplementary Fig. 26**). Palantir was also used to derive a pseudo-temporal order of progression of cells in the erythroid trajectory (**Supplementary Fig. 26**). Note that the pseudo-time order does not take measurement time into consideration and represents the potential journey of a cell through the cell-state space as it acquires erythroid fate. Further, cells measured at any timepoint can span a range of pseudo-time depending on the developmental stage.

The Palantir fate probability of cell state 𝑥_’_ reaching fate 𝐹 is represented by the function 𝑓(𝑥_’_, 𝐹). Accordingly, we define our threshold function for fate 𝐹 at pseudotime 𝑡 as:

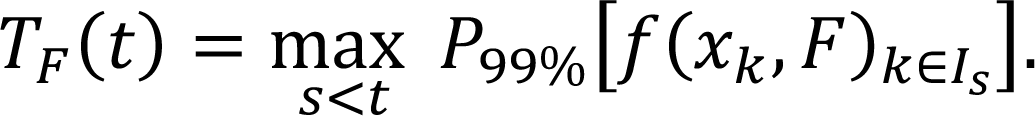

In this equation, 𝑃_PP%_ is the 99% percentile function and 𝐼_A_ is the set of all cells whose Palantir pseudotime is less than or equal to 𝑠. We then identify the subset of cells that are part of the branch leading to fate 𝐹 as:

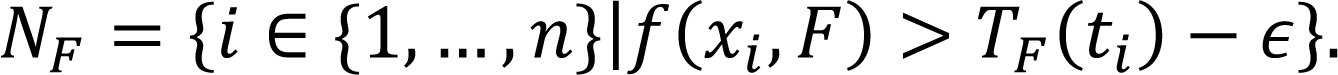

In the above equation, 𝑡_’_ is the pseudotime of the 𝑖^IS^ cell, and 𝜖 is a small chosen value (in our case, 0.01), which manages how much a cell can fall below the threshold while still being accepted as part of the branch. To simplify computation, we only calculate 𝑇_N_ for 500 specific pseudotime points along the trajectory, using the next larger pseudotime relative to 𝑡_’_ in this range to evaluate 𝑇_N_(𝑡_’_). This algorithm has been incorporated into the existing Palantir python package.

We next determined the joint cell-state density between pseudo-time and real time leveraging the time-continuous density function. We first used Gaussian process as implemented in Mellon to map pseudo-time to each coordinate of the cell-state space. This effectively generates a trajectory traversing the cell-state space by mapping the 1-dimensional pseudotime to high-dimensional cell-state space. Formally, the trajectory for each dimension 𝑚 ∈ (1, …, 𝑑′) is defined via the mean of the posterior distribution of 𝑇^7^ in the Bayesian model:

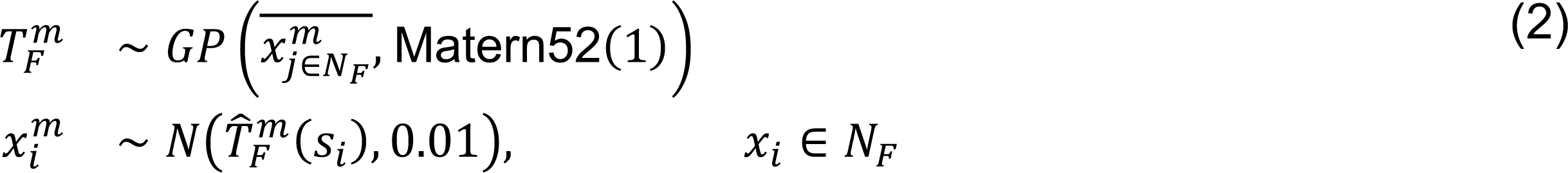

Where 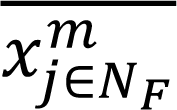 represents the average of this coordinate across all cells in branch 𝐹. The trajectory can then be denoted by

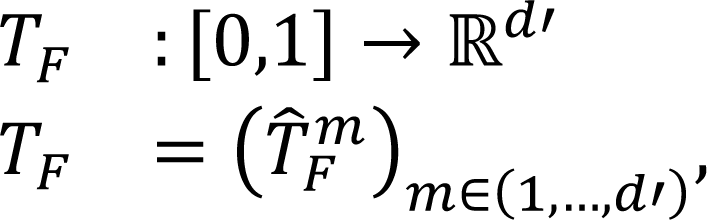

where *T̂^m^_F_* is the mean of the posterior of (2). The length scale of 1 and variance of 0.01 were selected by examination of a range of values for compatibility with cell states represented via Palantir diffusion maps.

Finally, the time-continuous density function 𝜌: ℝ^d′^ × [0,1] → ℝ^+^ can be evaluated along the trajectory to calculate joint cell-state density 𝜌(𝑇_N_(𝑠), 𝑡) for any given pseudotime 𝑠 and actual time 𝑡 (**Fig. 4F**).

#### Marginal Cell Type Proportions over Time

We used the joint cell-state density 𝜌(𝑇_N_(𝑠), 𝑡) to determine the dynamics of cell-type proportions over real time. We first assign a cell type to each section of the pseudo-temporal trajectory 𝑇_N_. This is achieved by computing a density function 𝜌_T_ for each annotated cell type 𝐻 using Mellon. The cell type annotation ℎ(𝑠), for a given pseudotime 𝑠 is then given by the largest cell type density for this point on the trajectory as follows

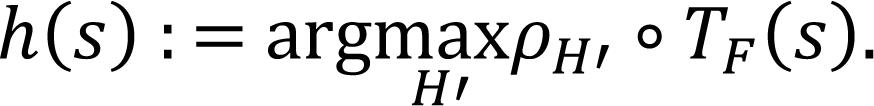

The cell-type annotation pseudotime 𝑠 can then be represented as an indicator function:

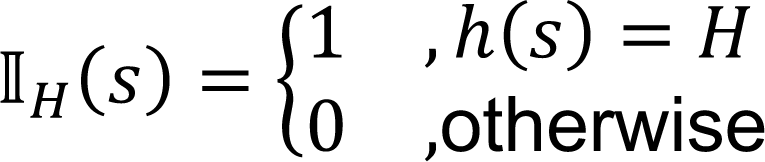

We next marginalized the joint cell-state density 𝜌(𝑇_N_(𝑠), 𝑡) over pseudo-time to determine the total mass of a cell type. Specifically, the mass of cell type 𝐻 along the trajectory of fate 𝐹 at a real time point 𝑡 is determined as

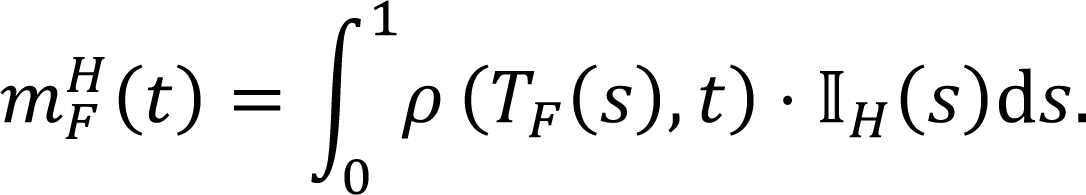

Finally, the relative proportion of cell type 𝐻 at a real time 𝑡 is given by normalizing the masses across all cell-type as follows:

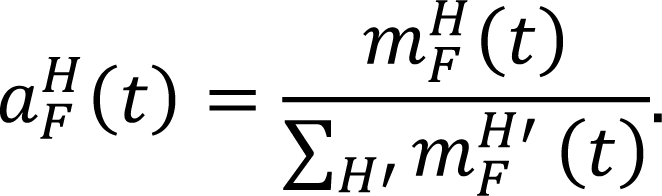

This provides a quantifiable measure of cell type proportions over time, offering valuable insights into the temporal evolution of cell types in a given biological system.

#### Application to mouse gastrulation data

We applied Mellon to determine the time-continuous cell-state density for the mouse gastrulation data^44^ across all measured time points: E6.5, E6.75, E7.0, E7.25, E7.5, E7.75, E8.0, E8.25 and E8.5. Data was preprocessed as described in section “Mouse gastrulation data in Data preprocessing”. Diffusion maps were constructed using batch corrected PCs across all cells using the Palantir package^14^. We selected 25 components, as they encapsulated all significant biological variations. Density results remained stable beyond this point with respect to the number of components (**Supplementary Fig. 13**). Time-continuous densities were computed following the procedure described above with default parameters.

Palantir^14^ was to derive pseudo-temporal order and cell-fate propensities. Palantir was run with default parameters by using an Epiblast cell as the start and manually setting the following cell-types as terminals: Cardiomyocytes, Erythroid, Endothelial, Neural crest, Brain, Notochord, Allantois, ExE endoderm. Since our goal was to identify cells with high fate propensity to erythroid lineage, a finer resolution terminal state identification was not necessary. Erythroid lineage cells were identified using Equation (1). Joint cell-state density over pseudo-time and real-time were visualized using 200 points along pseudo-time and 500 points between every pair of measured timepoints.

### Data preprocessing

#### scRNA-seq data preprocessing and analysis

The following procedure was used for preprocessing scRNA-seq data across datasets unless specified otherwise: Raw counts were normalized by dividing the counts by the total counts per cell. The normalized data was multiplied by the median of total counts across cells to avoid numerical issues and then log-transformed with a pseudocount of 0.1. Feature selection was then performed to select the top 2500 most highly variable genes, which was used as input for principal component analysis with 50 components. PCs were used as inputs for leiden clustering and UMAP visualizations. Preprocessing and analysis was performed using the scanpy^63^ package.

Diffusion maps were computed using the Palantir^14^ package with default parameters and PCs as the inputs. The diffusion kernel was also used for MAGIC^2^ gene expression imputation.

Batch correction where applicable was performed using Harmony with default parameters^64^. Batch corrected PCs if applicable were used as inputs for UMAPs, diffusion maps, and imputation.

#### T-cell depleted bone marrow single-cell multiome data

Raw gene counts, ATAC fragment files and cell metadata were downloaded from^65^.

#### RNA modality

scRNA-seq data was processed using the procedure described in section “scRNA-seq data preprocessing and analysis”, which mimics the analysis in^21^.

##### Cell-type annotation

All hematopoietic stem and progenitor cells (HSPCs) were grouped as one cell-type in the T-cell depleted bone marrow. To achieve higher granularity among the stem and progenitor cells, we integrated this data with a dataset of CD34+ bone marrow cells using Harmony^64^. This dataset is enriched for stem and progenitor cells and thus the associated cell-type information can be utilized to better resolve the cell-types within the HSPC cluster of the T-cell depleted bone marrow data. Batch corrected PCs were used for leiden clustering, and the HSPC cluster of the T-cell depleted data were assigned to different stem and progenitor cell-types based on their clustering with the CD34+ bone marrow data. Clusters associated with the B-cell trajectory were annotated using the markers described in ^66^.

##### Mellon cell-state density

Mellon was applied with default parameters using 20 diffusion components to compute cell-state density. Gene change scores, primed accessibility scores, and lineage-specific accessibility scores were computed as described above. IL7R signaling targets were downloaded from Nichenet^67^ and signature scores were computed by averaging the z-scored imputed gene expression.

#### ATAC modality

ArchR^36^ pipeline was used for analysis of the ATAC modality. In ArchR, data was normalized using IterativeLSI and SVD to determine a lower-dimensional representation of the sparse data. The first SVD component showed greater than 0.97 correlation with log library size and was excluded from downstream analysis. SVD was used as input to cluster the data with leiden and visualization using UMAPs. SVD also served as input for computing diffusion and MAGIC imputation of peak accessibilities and gene scores. Peak calling was performed within ArchR using only the nucleosome free fragments as described in ^21^.

A handful of cells which passed the RNA QC thresholds did not clear the thresholds in the ATAC modality. RNA preprocessing and analysis was repeated after excluding these cells. Mellon was applied with default parameters using 20 diffusion components to compute cell-state density of the ATAC modality.

#### Palantir trajectories

Palantir^14^ was used to infer pseudo-temporal trajectories of hematopoietic differentiation. Palantir was applied to the RNA modality using default parameters with the number of diffusion components (n=8) chosen by the Eigen gap statistic. A CD34+ hematopoietic stem cell was used as the start. Terminal cells were manually specified for erythroid, monocyte, B-cells, plasmacytoid dendritic cells. Note that the pre-pro B state of the B-cell trajectory is almost exclusively defined by cell-cycle^66^ and hence Palantir was run with pre-pro B and naïve B as the terminals. The B-cell fate probability was then computed as the sum of pre-pro B and naïve B probabilities.

Cells with increasing probability towards each lineage were selected as the lineage cells highlighted in **Fig. 1D**. B-cell lineage cells were comprised of Hematopoietic stem cells (HSCs), Hemopoietic multipotent progenitors (HMPs), Common Lymphoid progenitors (CLPs), prepro B-cells, pre B-cells, pro B-cells and Naïve B-cells. pDC lineage cells were comprised of HSCs, HMPs, Myeloid precursors, and pDCs. Erythroid lineage cells were comprised of HSCs, Megakaryocyte erythroid precursors (MEPs) and erythroid precursors. Monocyte lineage cells were comprised of HSCs, HMPs, Myeloid precursors, monocyte precursors and monocytes.

Cells involved in lineage specification (highlighted cells in **Fig. 1D**) where chosen as the subset of the lineage cells spanning from HSCs to the cell-type where the fate propensity reached 1. B-cells: HSCs, HMPs, CLPs, prepro B-cells, pro B-cells. pDCs: HSCs, HMPs, MyeloidPre, pDCs. Erythroid lineage: HSCs, MEPs. Monocytes: HSCs, HMPs, Myeloid precursors, monocyte precursors and monocytes.

#### HCA bone marrow

The processed annData was downloaded from^27^. The downloaded data was pre-batch corrected across all donors. Cell types that do not differentiate in the bone marrow such as T-cells, NK cells and plasma cells were excluded from the analysis. Following the cell filtering, each donor was analyzed separately using the steps outlined in the section “scRNA-seq data preprocessing and analysis”.

Palantir^14^ was applied separately for each donor using the same procedure that was described for the T-cell depleted bone marrow dataset. Mellon was applied with default parameters using 20 diffusion components to compute cell-state density.

#### Pancreatic development

Processed anndata was downloaded from ^17^ and the data was generated by ^29^. The pre-computed UMAPs, cell-type annotations and diffusion maps were used for analysis. Mellon was applied with default parameters to compute cell-state density.

#### In-vitro endoderm differentiation

Raw counts and cell metadata was downloaded from^30^. Wild-type cells were used for all analysis. Data analysis was performed using the steps outlined in the section “scRNA-seq data preprocessing and analysis”, batch correction was used to correct technical differences between two batches. Mellon was applied with default parameters to compute cell-state density.

#### Spatial organization of intestinal tissue

Raw counts and zone information were downloaded from^31^ and processed using the steps outlined in the section “scRNA-seq data preprocessing and analysis”. Mellon was applied with default parameters to compute cell-state density.

#### Lung regeneration

Processed anndata was downloaded from^28^. The pre-computed UMAPs, cell-type annotations and diffusion maps were used for analysis. Mellon density functions were computed for each timepoint separately and evaluated across all cells.

#### scRNA-seq of murine models of lung adenocarcinoma

Processed anndata object containing counts, visualization and cell-metadata were downloaded from ^9^. scVI^68^ was used in the publication for data integration and to derive a latent representation. scVI latent space was used as input for computing force directed layouts and diffusion maps instead of PCs like other datasets.

#### Mouse gastrulation atlas

Processed data including batch corrected principal components and cell metadata were downloaded from^44^. Batch corrected PCs were used as input for computing diffusion maps. Cells from the “mixed_gastrulation” samples were excluded since the timepoints are not well-defined. Further, ExE ectoderm, ExE endoderm and Parietal endoderm cells were excluded since their parental cells are not measured in the dataset. Given the complexity of the data, 25 diffusion components for computing time-continuous cell-state densities using Mellon.

#### iPS reprogramming dataset

Raw counts and cell metadata were downloaded from^8^. The dataset contains reprogramming in two culture conditions: Serum and 2i. Cells cultured in 2i media were used for the analysis. Highly variable genes computed in the publication were used for the analysis using the steps outlined in the section “scRNA-seq data preprocessing and analysis”. iPS data was used for robustness analysis and benchmarking performance.

#### scATAC-seq of murine models of lung adenocarcinoma

Raw peak counts and cell metadata were downloaded from^49^. Immune and stromal cells were excluded from the analysis. Following cell filtering, peak counts were normalized using TFIDF following the procedure in ^21^. SVD was to determine a lower-dimensional representation using normalized data as input. The first SVD component showed greater than 0.97 correlation with log library size and was excluded from downstream analysis. SVD was used as input for visualization using force directed layouts and diffusion maps. Mellon was applied with default parameters to compute cell-state density.

#### sortChIC data profiling histone modifications in murine hematopoiesis

Raw peak counts and cell metadata were downloaded from^48^ for all available histone modifications: H3K4me1, H3K4me3, H3K27me3, H3K9me3. Each modification was analyzed separately following the procedure described in the section “scATAC-seq of murine models of lung adenocarcinoma”: Data was normalized using TF-IDF, and then SVD was used to derive a low-dimensional representation. The first component of SVD was excluded due to high correlation with log library size and was excluded from downstream analysis. Mellon was applied with default parameters to compute cell-state density.

#### Skin differentiation Share-seq data

The processed annData was downloaded from ^51^ using the data generated by ^32^. The pre-computed UMAPs, cell-type annotations and diffusion maps were used for analysis. Note that the diffusion components were derived using the MIRA multimodal representation which uses both RNA and ATAC modalities. Mellon was applied with default parameters to compute cell-state density.

### Robustness analysis

The robustness of Mellon was evaluated by recalculating density estimations across a broad spectrum of parameter settings on multiple datasets. We carried out full density inference for an extensive range of length scales, numbers of landmarks, and numbers of diffusion components in the following scRNA-seq datasets : T-cell depleted bone marrow of human hematopoiesis (BM)^21^; CD34+ human bone marrow cells, a dataset of hematopoietic stem and precursor cells (CD34)^21^; COVID-19 atlas of peripheral blood mono nuclear cells (PBMCs) from healthy donors and critical patients (Covid)^69^; iPS reprogramming dataset (ips)^8^ and the mouse gastrulation atlas (mgast)^44^. These datasets cover a broad spectrum of systems with different complexities, cell numbers and contain discrete and continuous cell-states and cell-types. We compared the densities using Spearman correlation between results obtained from different parameter settings. As shown in **Supplementary Fig. 13-16**, Mellon results exhibited a high level of consistency in the results even when the parameters are varied orders of magnitude beyond the defaults.

We further evaluated Mellon’s robustness to down sampling the cells in the dataset. Starting with the full dataset, we serially removed 10% of cells until at least 100 cells were retained. We next computed densities for independently for each subsample by recomputing the principal components and diffusion components using only the cells in the subset. We then compared the density between all pairs of subsamples using the intersection of cells between the two samples (**Supplementary Figure 11**). The consistency is retained even when cells in the bottom 10^th^ percentile of the average density between the pair of runs are used for comparison (**Supplementary Fig. 12**).

This robustness evaluation provides empirical evidence of Mellon’s ability to perform consistently under a wide range of parameters and under the condition of subsampling, which underscores its utility for accurate density estimation from high-dimensional single-cell data.

### Simulated datasets with ground-truth densities

In order to validate the accuracy and precision of Mellon, we generated three datasets mirroring single-cell datasets of either continuum of cell-states or discrete clusters. Each dataset is accompanied by a predefined ’ground truth’ density serving as a performance benchmark for Mellon.

The datasets with continuum of cell-states were generated using a large Gaussian Mixture Model (GMM) designed to emulate a cellular differentiation tree. This tree was conceptualized as a series of velocity vectors, each connecting branching points and each being a slightly perturbed version of the vector of its parent node. For each node in this tree, a unique Gaussian was defined. The Gaussian’s covariance matrix and mean were designed to create a distribution aligning with the velocity vector. Considering the inherent low dimensionality typically exhibited by a cell-state manifold, we adjusted the principal components of these Gaussians using an exponential decay scalar. The two continuous styles mimic the structure of CD34+ bone marrow RNA-seq and T-Cell depleted bone marrow RNA-seq datasets.

The synthetic datasets representing single-cell datasets of discrete clusters was also generated using a GMM but with a different configuration. In this setup, we randomly sampled mean and covariance matrices to create an arbitrary GMM, resulting in mostly isolated clusters of simulated cells. This approach provided an alternative, contrasting framework for testing robustness of Mellon.

The GMM allowed us to easily sample simulated cell states from both dataset types and to define corresponding ground truth probability density functions. We then utilized Mellon to compute the log-density of these simulated datasets. Ground-truth densities were compared with Mellon densities using Spearman correlations. As shown in **Supplementary Fig. 4**, this comparison effectively quantified Mellon’s ability to infer cell densities from high-dimensional single-cell data, with Mellon exhibiting high consistency with the ground truth for both synthetic datasets.

See **Supplementary Note 5** for further details and parameter choices for dataset simulations.

### Comparison to density estimation approaches

A commonly used approach for density estimation with single-cell data is to calculate the reciprocal of the distance to the kth nearest neighbor, treating this value as a proxy for density^2^. While straightforward, this method tends to produce a noisy density estimation and frequently fails to capture meaningful global trends (**Supplementary Fig. 2**).

Another prevalent approach involves application of kernel density estimation (KDE) to the low-dimensional embeddings generated by tools like UMAPs or Force-Directed Layouts. While these visualization tools are powerful, their main design is not for density inference. They can produce unstable embeddings, and when KDE is applied, the instability in the embeddings directly translates into the density inference, resulting in less reliable outputs. Furthermore, the high compression involved in generating these low-dimensional representations means that they cannot capture all the relevant biological variability inherent in the data. Consequently, these methods often fail to depict all the nuanced details of the underlying cell-state density function (**Supplementary Fig. 2**).

### Efficient Pseudotime Trend Computation with Mellon

The versatility of Mellon extends beyond density inference, showcasing its robust capability in the swift computation of gene trends, defined as continuous, smooth functions that trace the trajectory of gene expression over pseudotime. Our Gaussian process (GP) regression-centric design not only serves as the backbone for Mellon’s primary application but also efficiently caters to general GP regression, due to scalable features such as the fixed length scale for the Matern52 covariance kernel and landmarks for Sparse Gaussian process regression.

Gaussian processes shine in their adeptness at handling high-noise scenarios, for instance, non-imputed gene expression values. This strength enables Mellon to generate smooth gene trends from a selected cellular branch’s temporal ordering using unimputed gene expression values, effectively capturing the dynamics of gene expression as cellular differentiation unfolds (**Supplementary Fig. 21**).

Mellon’s implementation harnesses the power of the JAX library’s vectorization capabilities and low-dimensional latent representations of functions within the GP framework, enabling efficient gene trend computations across a substantial quantity of genes. In tests using a 36-core CPU, Mellon was able to generate gene trends for up to 10,000 genes and 1,500 cells at 500 pseudotime points in about one second. This efficient computation allows high-throughput exploration of gene expression dynamics during cellular differentiation from large-scale single-cell datasets.

