## Supplementary Figures for "Quantifying Cell-State Densities in Single-Cell Phenotypic Landscapes using Mellon"

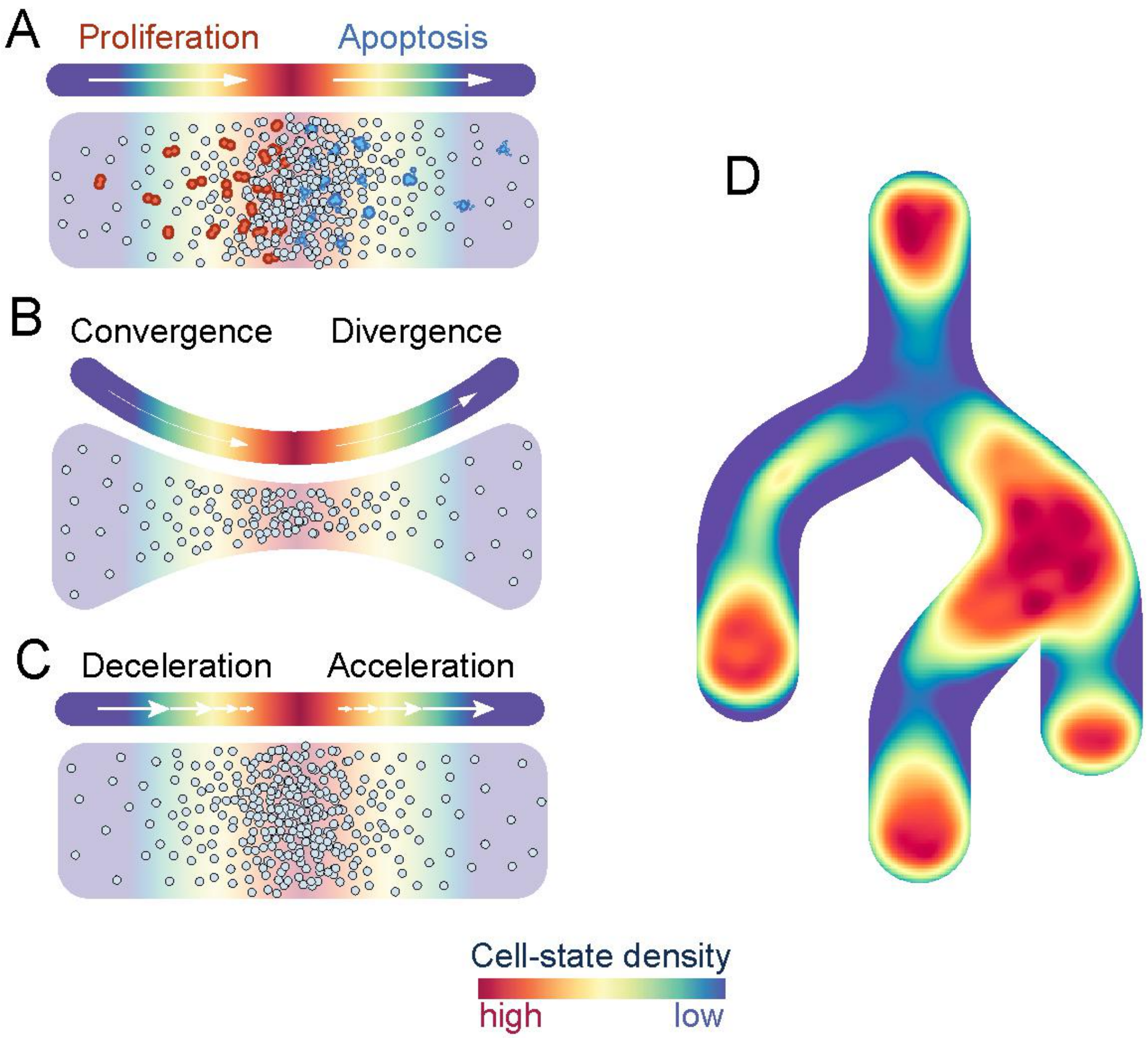

**Supplementary Figure 1**: **Schematic representation of cell differentiation dynamics impacting cell-state density**

A. Proliferation and apoptosis directly impact cell-state density, respectively increasing or decreasing it.

B. Convergence and divergence during cell-state progression along the differentiation trajectory. Convergence forces cell states into a more confined state space, thereby increasing density, whereas divergence spreads out cell states, resulting in a decrease in density.

C. The pace of state changes also influences cell-state density. Deceleration of state changes can result in a concentration of cell states, thereby increasing density. Conversely, acceleration of state changes tends to disperse cell states, decreasing density. In gene expression, acceleration is induced by rapid transcriptional changes.

D. Schematic of the toy dataset 1 to illustrate the Mellon continuous density function.

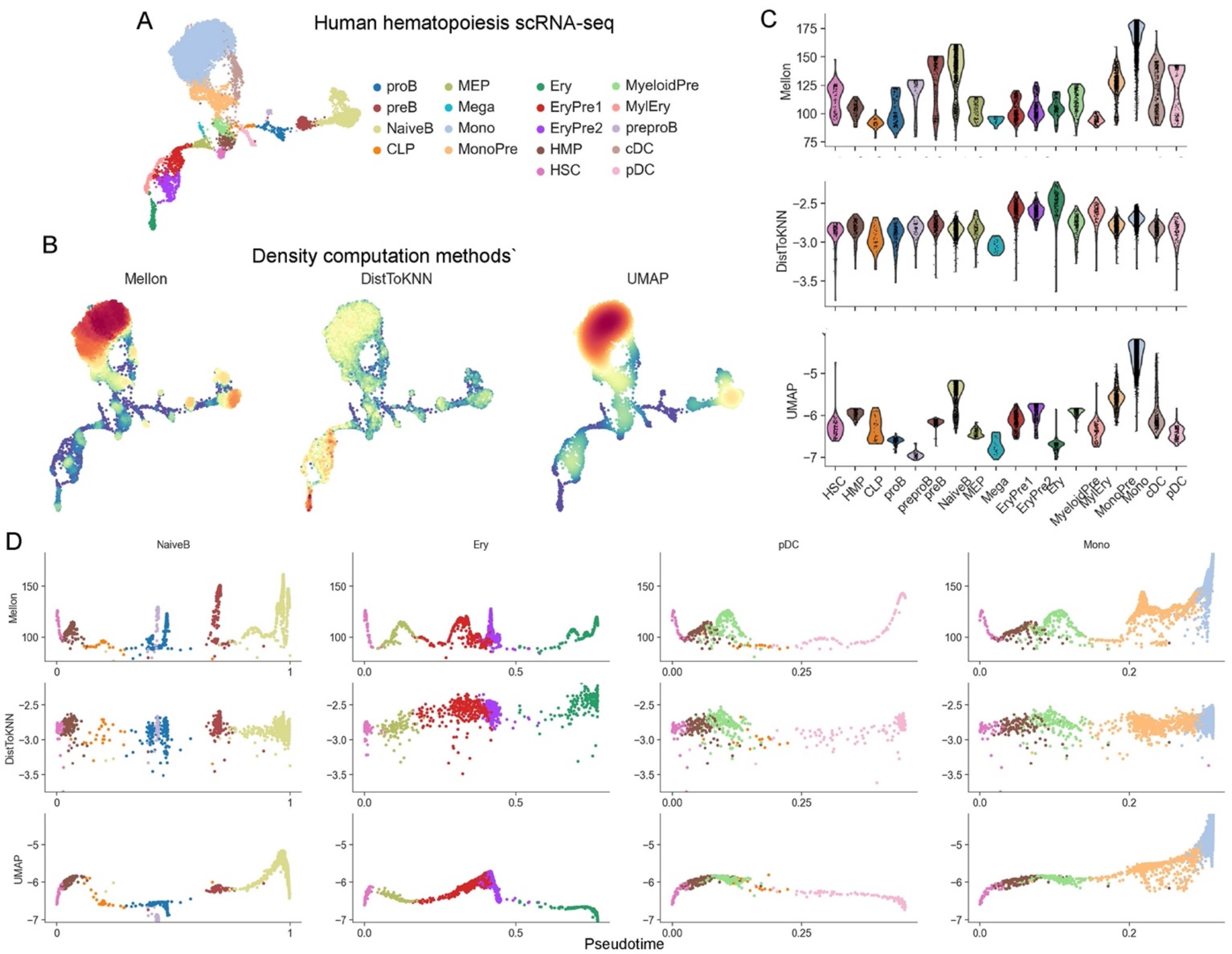

**Supplementary Figure 2: Comparison of cell-state density estimation approaches**

A. UMAP of scRNA-seq dataset of T-cell depleted bone marrow dataset, colored by cell-type.

B. UMAPs colored by Mellon density (left), density computed as inverse of distance to kth nearest neighbor (middle) and density computed using UMAP coordinates (right). K =15

C. Violin plots to compare cell-state densities among different hematopoietic cell-types. Arrowheads indicate example cell-types with high variability in density. Top: Mellon, Middle: Inverse of distance to kth nearest neighbor, Bottom: UMAP densities. Mellon densities are most consistent with expected landscape of human hematopoiesis.

D. Plots comparing Palantir pseudotime to log-density for different hematopoietic lineages. Top row: Mellon, Middle row: Inverse of distance to kth nearest neighbor. Bottom row: UMAP densities. Mellon provides the most robust and interpretable density estimates with clear separation of high- and low-density regions.

Log densities are shown for all comparisons.

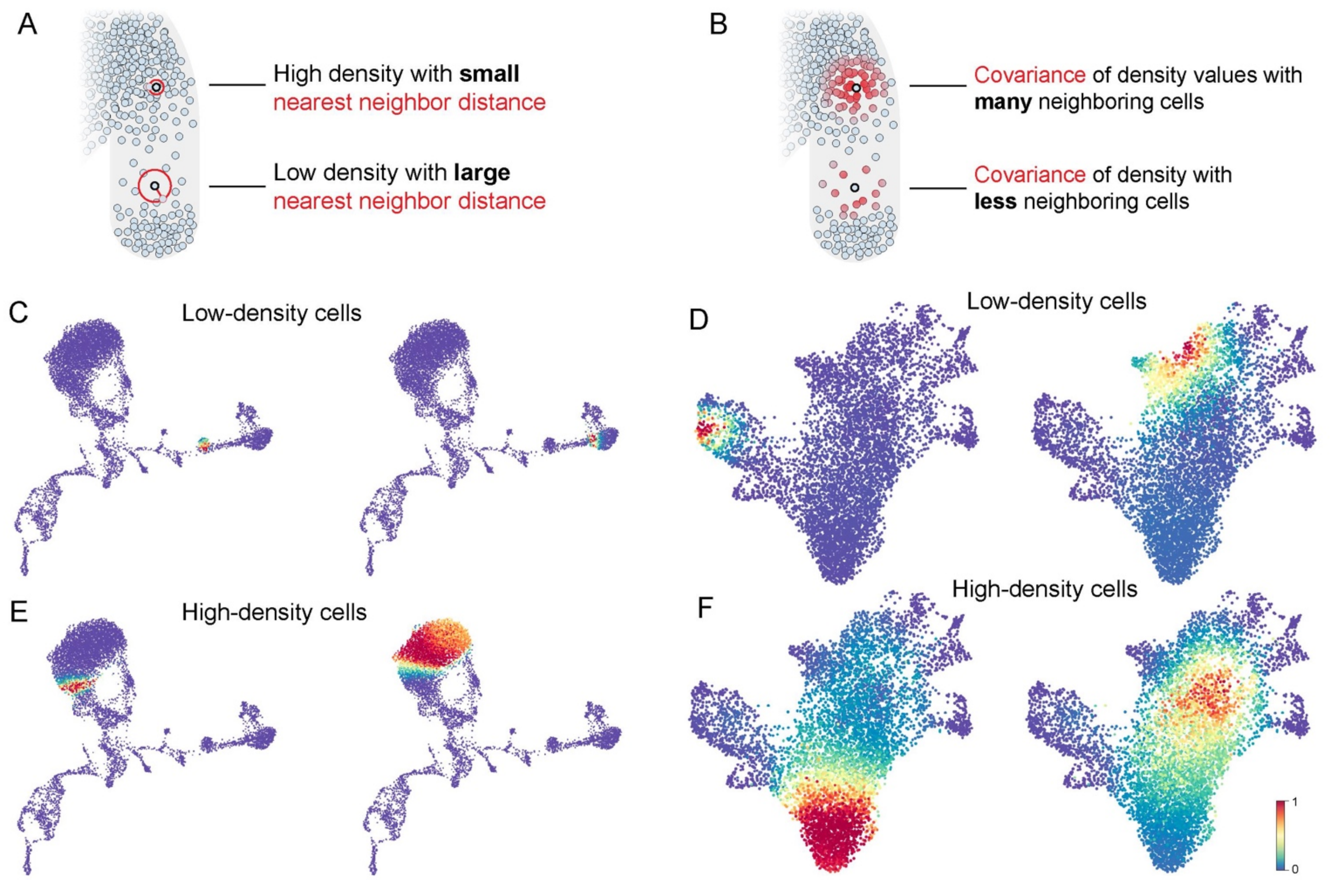

**Supplementary Figure 3: Illustration of how the Covariance Kernel of the Gaussian Process links cell-state density inference across related cells.**

A. Inset of the toy dataset in **Fig. 1B**, showing nearest neighbor distance of representative cells in high- and low- density regions

B. Representative cells displaying the covariance function (more intense red indicates higher covariance). This gradient demonstrates the degree of covariance with neighboring cells: a cell located in a high-density region shows high covariance of density with many neighbors, whereas a cell in a low-density region exhibits strong covariance with fewer cells.

C. UMAPs of the T-cell depleted bone marrow dataset colored by covariance to randomly selected cells in low-density regions. Covariance between all pairs of cells serve as input to Gaussian Process.

D. Same as (C), for the CD34+ bone marrow data.

E. UMAPs of the T-cell depleted bone marrow dataset colored by covariance to randomly selected cells in high-density regions.

F. Same as (E), for the CD34+ bone marrow data.

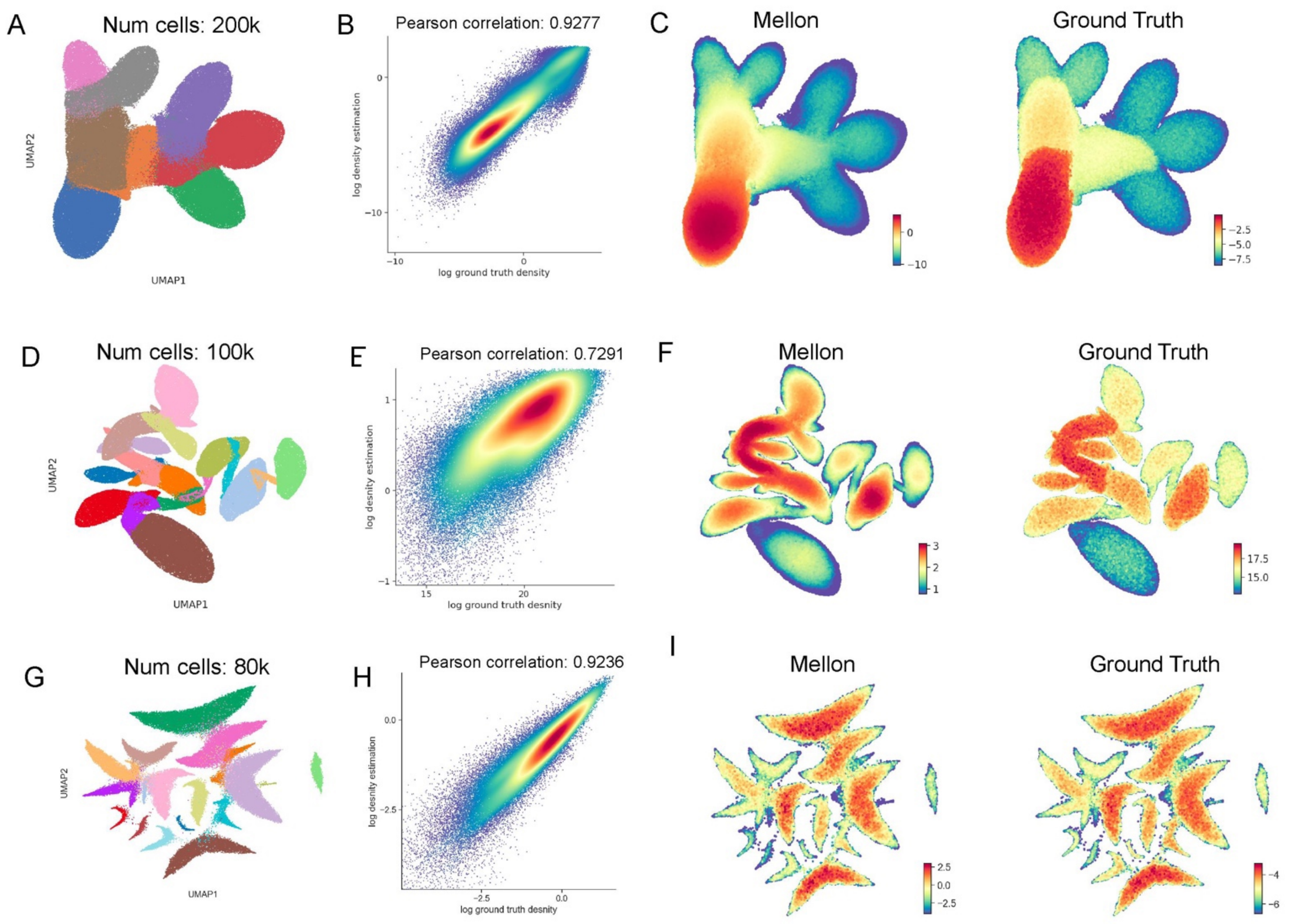

**Supplementary Figure 4: Validation of Mellon density estimation using simulated datasets with known ground truth.**

A. UMAP of simulated data colored according to differentiation tree nodes

B. Correlation plots between known ground truth log-density (x-axis) and Mellon-inferred density (y-axis) across all simulated cells. Each point represents a simulated cell.

C. MAPs colored by Mellon-inferred density (left) and ground truth log-density (right). Density values falling below the 20th percentile are projected to the 20th percentile for visualization

D-F. Same as A-C for a second simulated dataset. Cells in (D) are colored by differentiation tree nodes

G-I. Same as A-C for a third simulated dataset. Cells in (G) are colored by clusters.

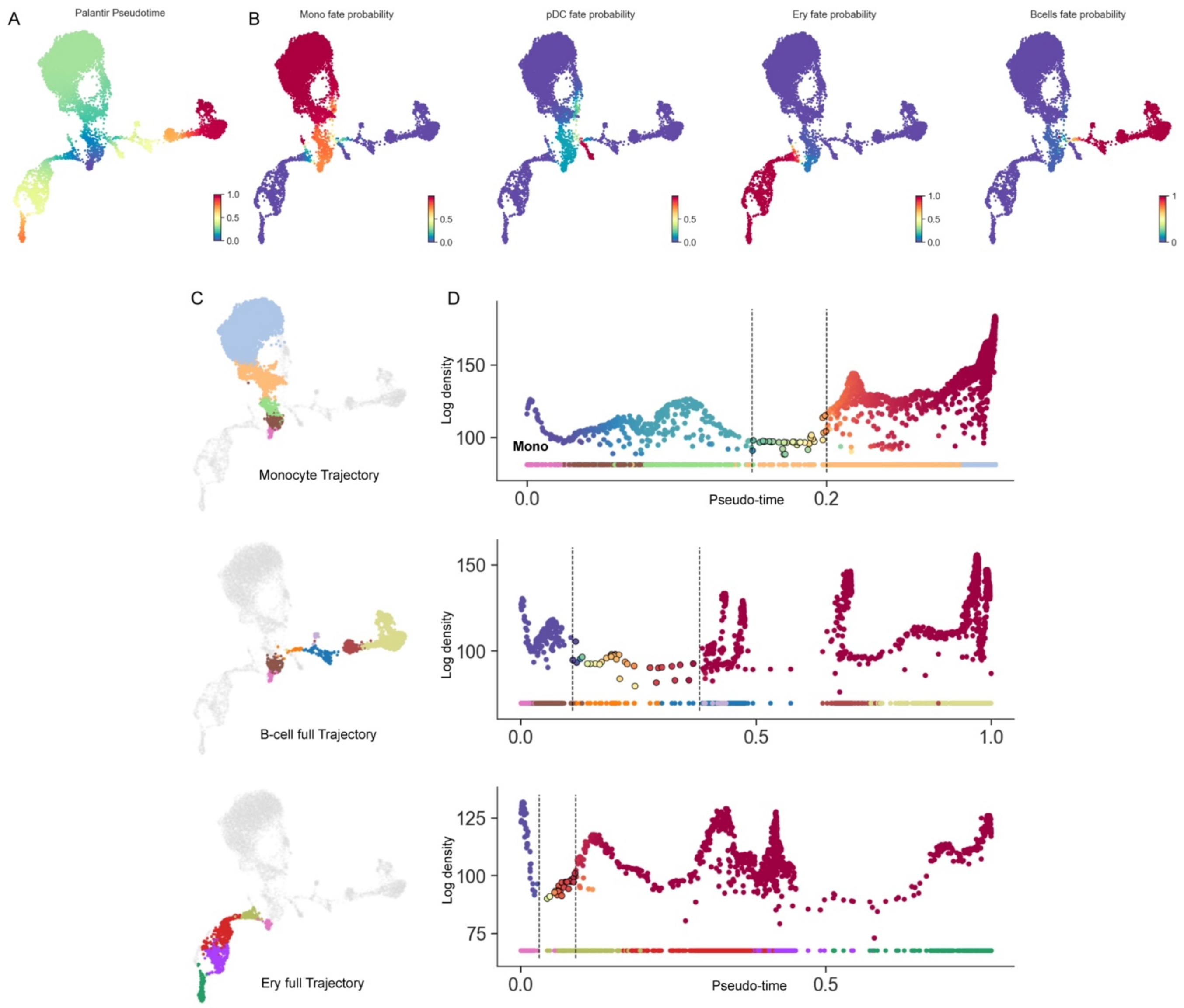

**Supplementary Figure 5: Cell-fate trajectories in the T-cell depleted bone marrow scRNA-seq dataset using Palantir.**

A. UMAP with cells colored by Palantir pseudotime.

B. UMAPs with cells colored by the fate propensities towards terminal cell states : monocytes (Mono), plasmacytoid dendritic cells (pDC), erythroid cells (Ery), and B-cells.

C. UMAP with cells of the monocyte lineage highlighted.

D. Plots comparing pseudotime and log-density with cells colored by monocyte fate probability. Vertical regions separate high and low-density regions and were chosen manually. Cells in the low-density region representing fate specification are highlighted. B-cell and erythroid trajectories are same as **Fig. 2D-E** with all cells of the trajectory.

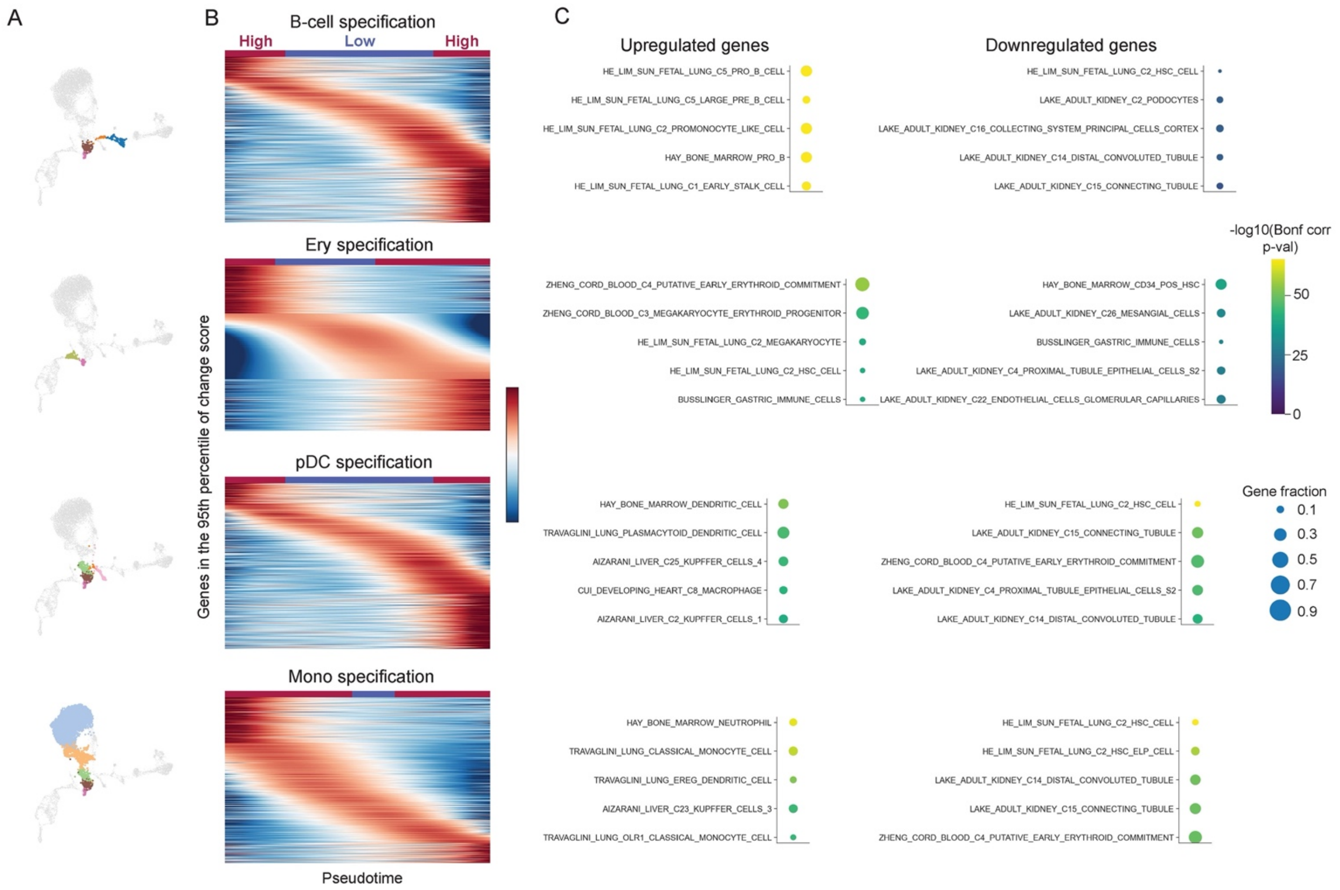

**Supplementary Figure 6: Genes driving low-density state transitions**

A. UMAPs highlighting the cells spanning hematopoietic stem-cells to fate committed cells along each lineage.

B. Heatmaps showing the gene expression dynamics along pseudotime for genes in to the 95$^{th}$ percentile of change scores associated with the respective lineage. High and low-density regions were assigned manually by comparing Mellon density with Palantir pseudotime (**Fig. 2E**).

C. Gene ontology results of up and downregulated genes along each lineage. Genes with higher expression in the first high-density region in (A) were nominated as downregulated and the rest as upregulated. Bonferroni corrected p-values were determined using hypergeometric tests using the c8 gene set from MSigDB[1].

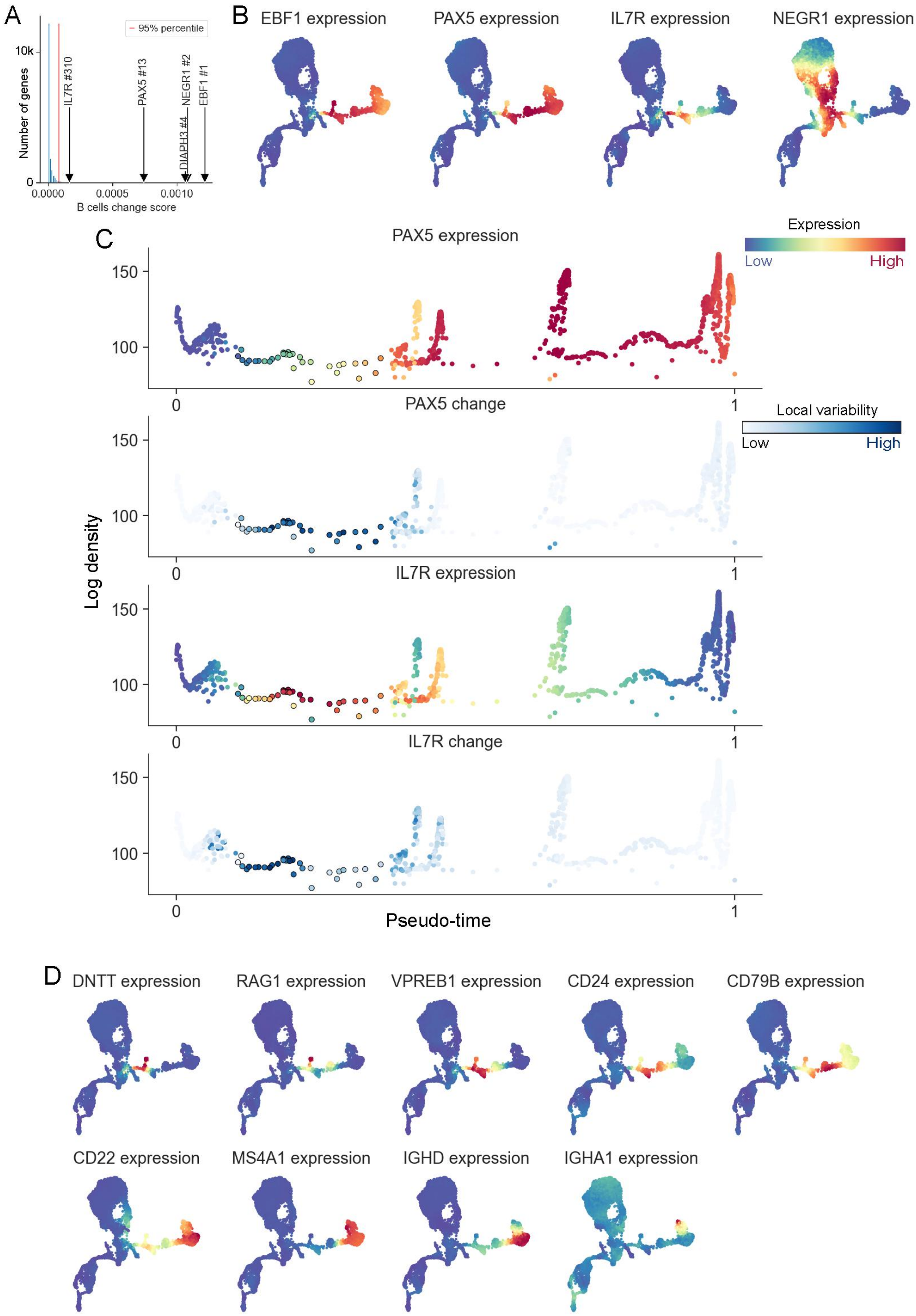
A
95% percentile
10k
0
Number of genes
IL7R #310
PAX5 #13
DIAPH3 #4
NEGR1 #2
EBF1 #1
0.0000
0.0005
0.0010
B cells change score
B
EBF1 expression
PAX5 expression
IL7R expression
NEGR1 expression
C
PAX5 expression
Expression
Low
High
PAX5 change
Local variability
Low
High
IL7R expression
IL7R change
Log density
150
100
0
1
Pseudo-time
D
DNTT expression
RAG1 expression
VPREB1 expression
CD24 expression
CD79B expression
CD22 expression
MS4A1 expression
IGHD expression
IGHA1 expression

**Supplementary Figure 7: Gene expression dynamics along B-cell differentiation.**

A. Histogram showing the distribution of gene change scores for the low-density cell-states encompassing hematopoietic stem cells through pro B-cells. EBF1 has the highest change score.

B. UMAPs colored by expression of key B-cell regulators

C. Plots comparing Palantir pseudo-time with Mellon log density along B-cell trajectory with cells colored by PAX5 expression, PAX5 expression change, IL7R expression and IL7R expression change. Cells involved in B-cell fate specification are highlighted.

D. UMAPs colored by expression of B-cell checkpoint markers. MAGIC imputed expression is used for visualization.

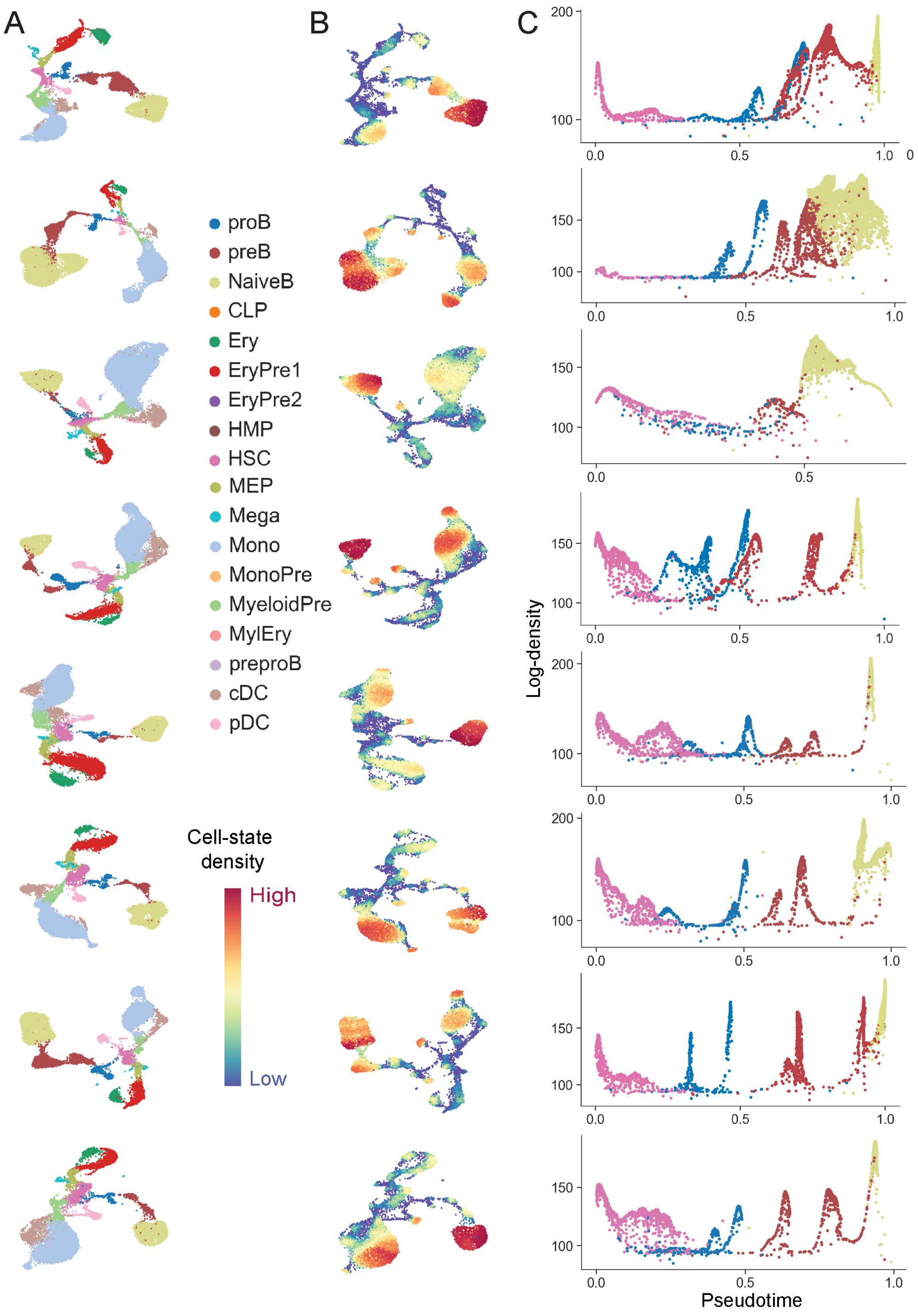
A
B
C
proB
preB
NaiveB
CLP
Ery
EryPre1
EryPre2
HMP
HSC
MEP
Mega
Mono
MonoPre
MyeloidPre
MylEry
preproB
cDC
pDC
Cell-state density
High
Low
Log-density
Pseudotime

**Supplementary Figure 8: Mellon densities are robust across independent bone marrow donors**
A. UMAPs colored by cell-types for seven different bone marrow donors from the Human Cell Atlas[2]. T-cells were excluded from this analysis
B. UMAPs colored by Mellon log-density
C. Plots comparing pseudotime with log density across different donors for the B-cell trajectory.

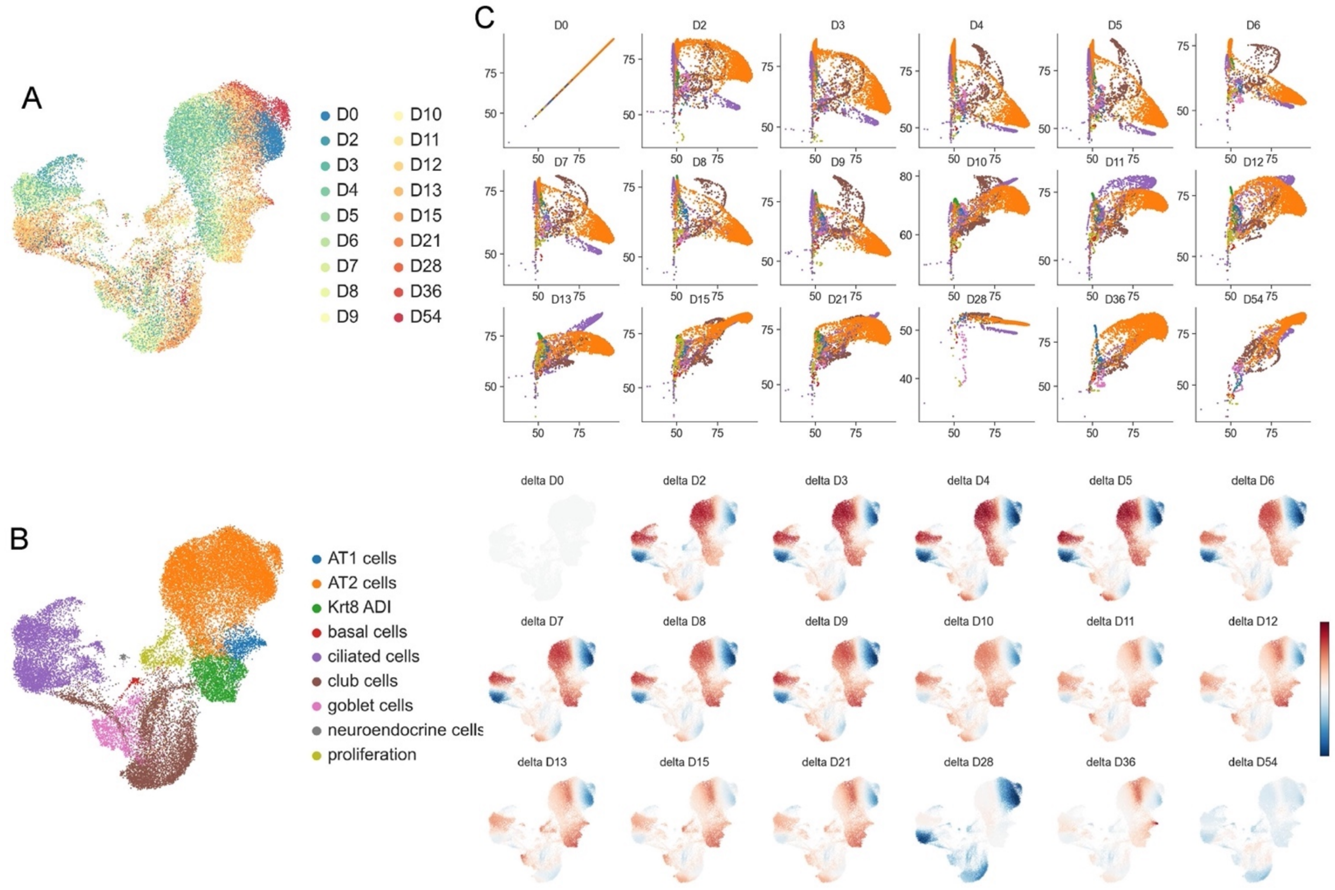

**Supplementary Figure 9: Cell-state density is a property of the homeostatic system**

A. UMAP of a scRNA-seq dataset of lung regeneration[3] with cells colored by time point of measurement. D0 is prior to injury and the subsequent timepoints show recovery from injury induced by bleomycin.

B. Same as (A), colored by cell-types.

C. Plots comparing log-density of D0 with the specified timepoints. Subset of cells from D0 and the corresponding timepoint were used for this comparison.

D. UMAPs colored by the difference of densities between D0 and the specified timepoint.

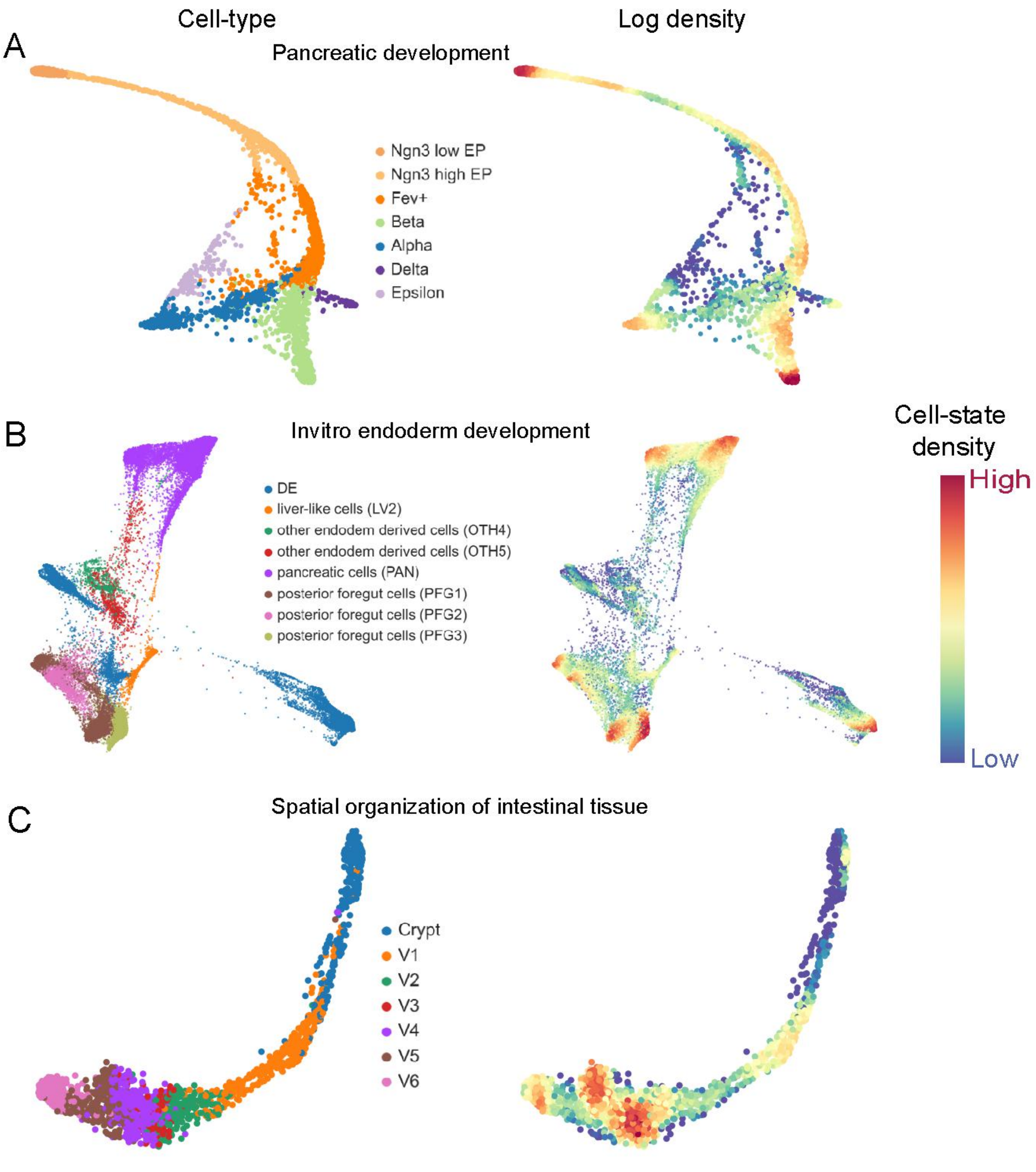

**Supplementary Figure 10: Mellon characterizes cell-state densities in diverse differentiation systems**

A. UMAPs colored by cell-types (left) and Mellon log density (right) for the scRNA-seq dataset of murine pancreatic development at E15.5[4].

B. Same as (A) for scRNA-seq dataset of invitro human endoderm differentiation[5].

C. Same as (A) for scRNA-seq dataset of scRNA-seq dataset describing the spatial organization of murine intestinal tissue[6].

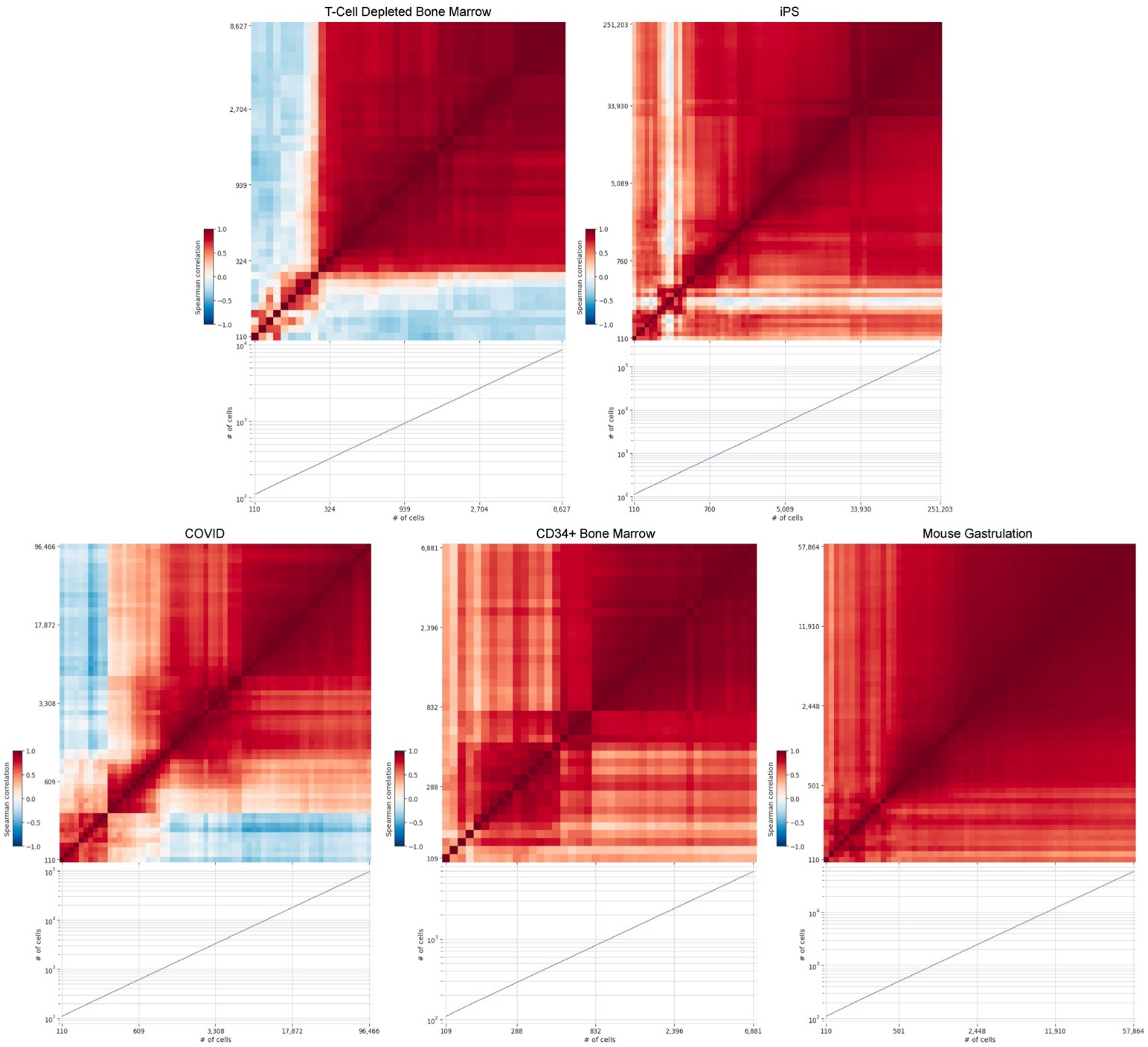

**Supplementary Figure 11: Mellon is robust to subsampling of real datasets.**
Heatmaps displaying Spearman correlation between density estimations derived from hierarchically subsampled versions of five datasets. The marginal plot at the bottom denotes the number of cells retained for the dataset in each column. The right-most column and the topmost row represent the density estimates from complete datasets. Datasets are progressively subsampled from right to left, with each subsequent subsample reducing the number of cells by 10%. All colormaps are equivalently scaled from -1 to 1, representing the full range of Spearman correlation values. This systematic reduction and corresponding correlation analysis illustrate the robustness of Mellon's density estimates to the size of the input dataset.

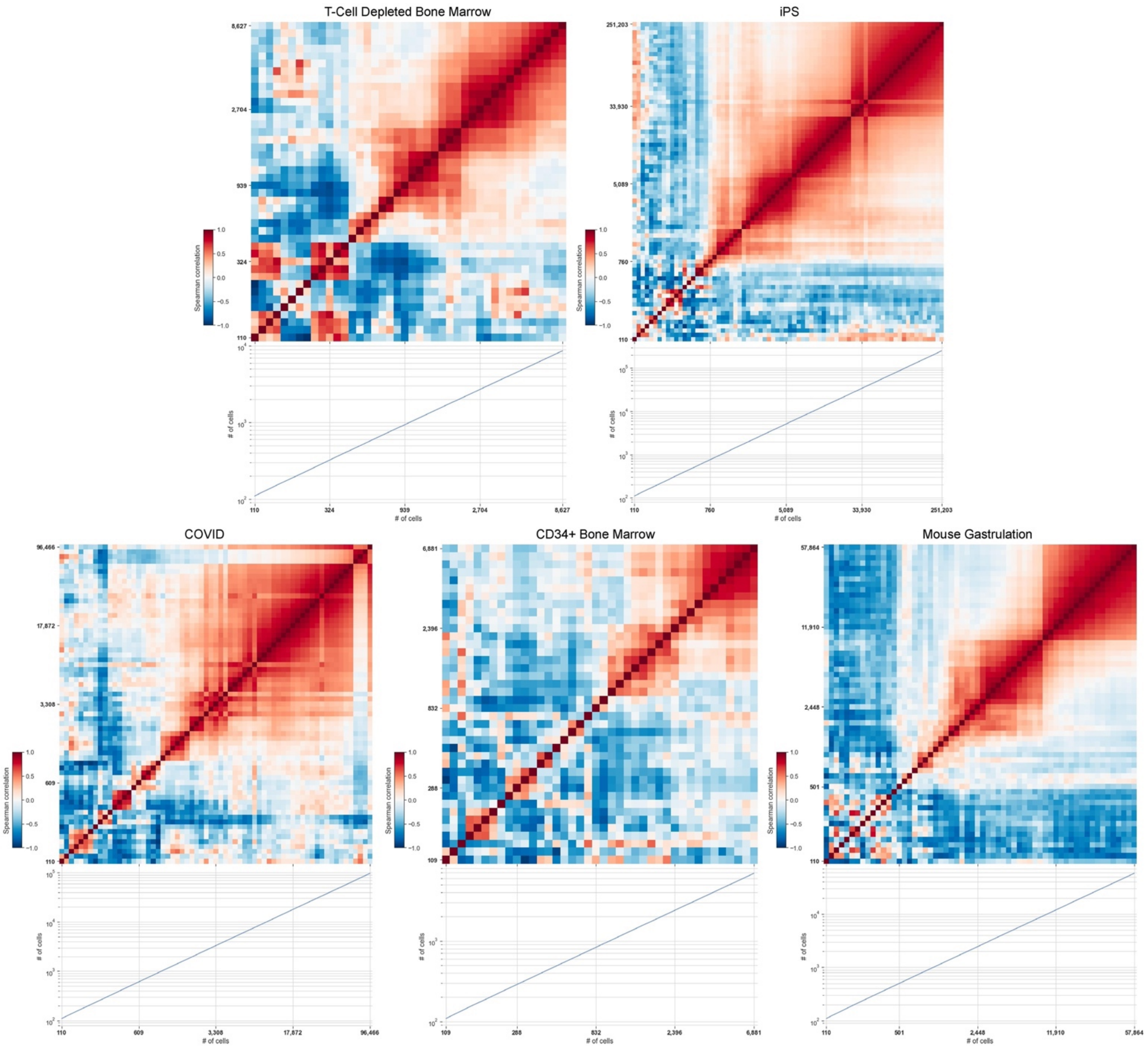

**Supplementary Figure 12: Mellon produces robust densities within lower-density regions.** Same as **Supplementary Figure 11,** with Spearman correlations computed for the subset of cells in the bottom 10% of density, determined by averaging the densities of the two estimates being compared.

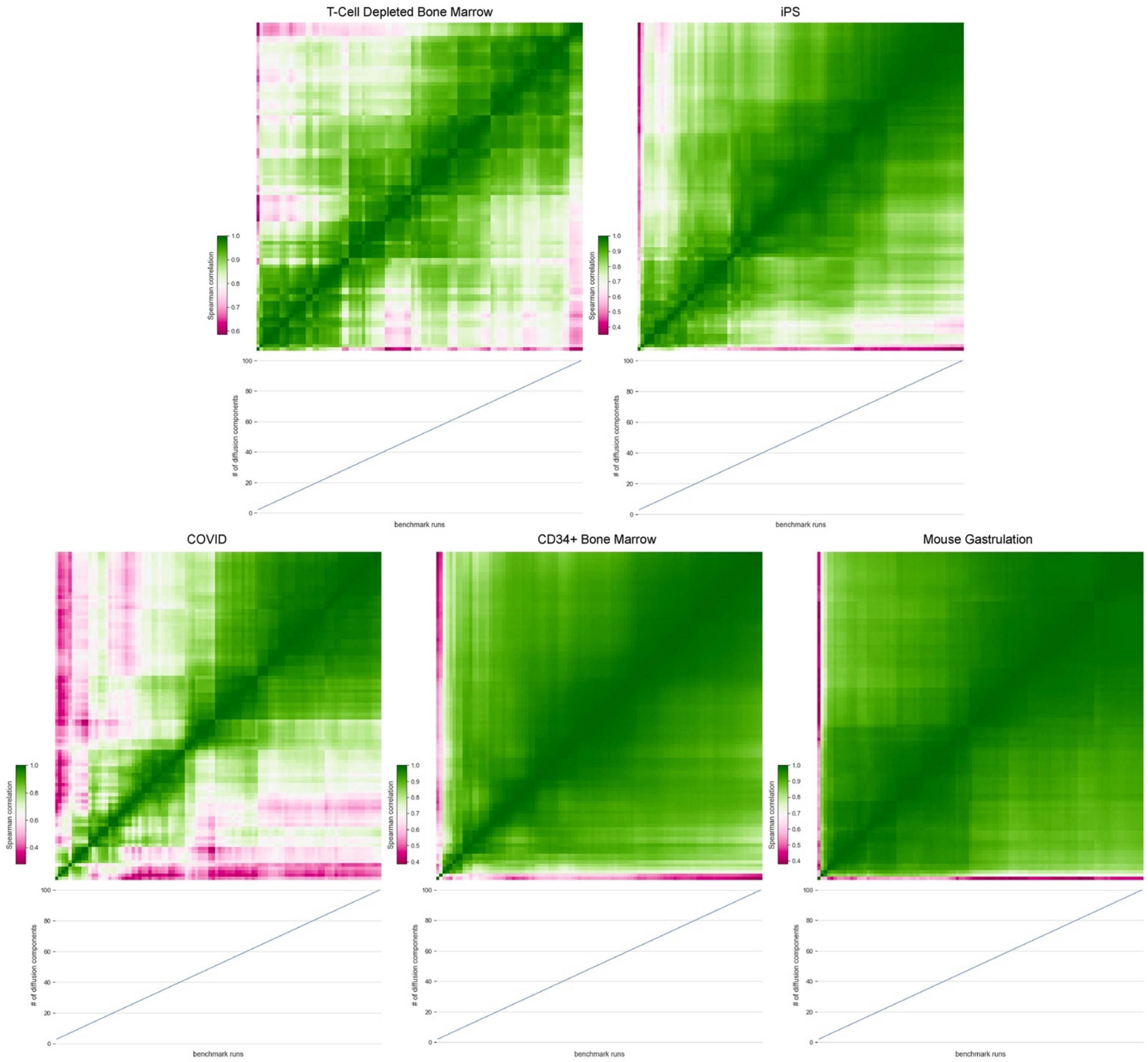

**Supplementary Figure 13: Mellon densities are robust to varying number of diffusion components.**

Heatmaps displaying Spearman correlation between density estimates, each generated using a different number of diffusion components in the cell-state representation for the five datasets. The number of diffusion components applied for each column's density estimate is depicted in a marginal plot at the bottom. The number of diffusion components increases from left to right. Each heatmap employs an individual color-scale, optimized to display the range of correlation values within that specific dataset. Despite variations in the number of diffusion components used, the consistently high correlation values demonstrate Mellon's ability to maintain robust density inference across different dimensionalities of cell-state representations.

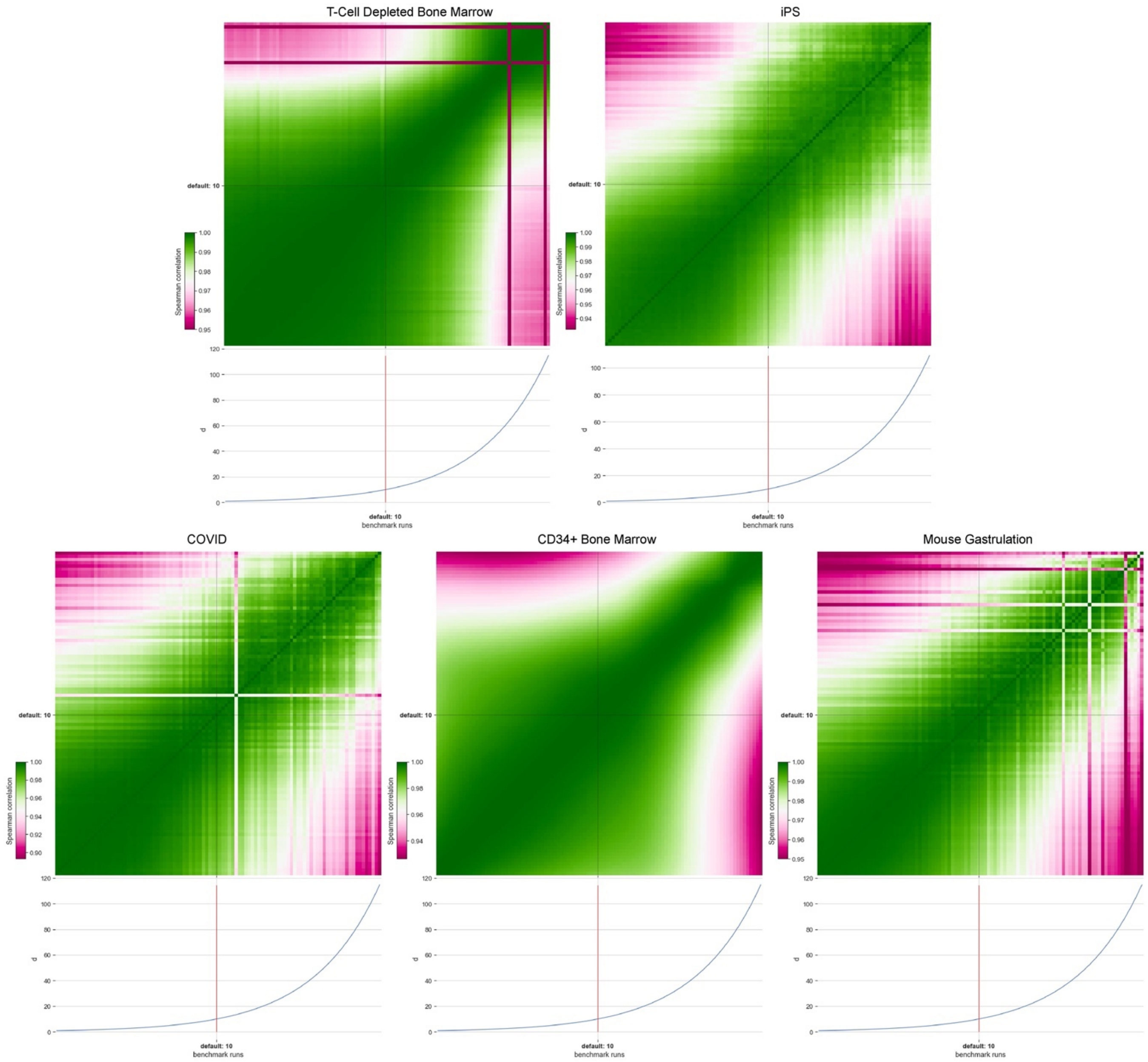

**Supplementary Figure 14: Robustness of Mellon densities to variation in the parameter 'd'.** Heatmaps present the Spearman correlation between density estimates, each produced with a different value for the parameter 'd' in the cell-state representation for the five datasets. The value of 'd' applied for each column's density estimate is represented in a marginal plot at the bottom, increasing from left to right. Every heatmap utilizes an individual color scale, tailored to highlight the range of correlation values within that specific dataset. Regardless of the changes in the 'd' parameter value, the consistent high correlation values attest to Mellon's robustness in density inference across different settings of this parameter.

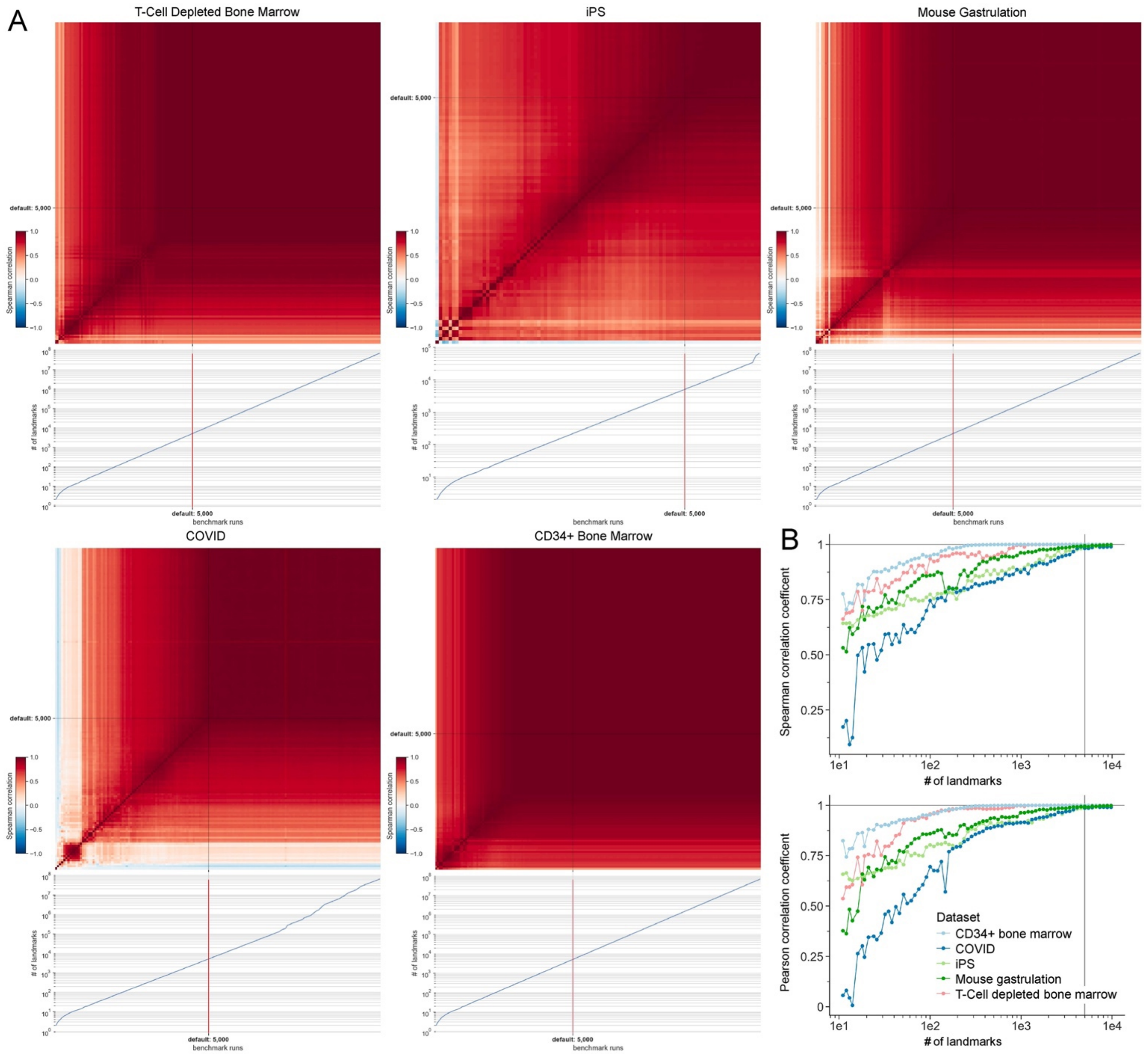

**Supplementary Figure 15: Mellon densities are robust to varying number of landmarks.**
A. Heatmaps displaying Spearman correlation between density estimates derived using different numbers of landmarks for five datasets. The number of landmarks used for each column's density estimate is indicated by a marginal plot at the bottom. The right-most column and topmost row correspond to the maximum number of landmarks tested, which is typically substantially more than the default of 5,000 landmarks. This default value is indicated by a vertical line traversing both the heatmap column and the associated marginal plot. As the number of landmarks decreases from right to left, the resulting correlations provide insights into the robustness of Mellon's density inference as a function of landmark quantity. The colormaps are uniform across all heatmaps, ranging from -1 to 1, covering the entire possible range of Spearman correlation values. High correlation values adjacent to the top-right corner

signify that Mellon's density inference maintains strong accuracy, even with varying numbers of landmarks.

B. Spearman and Pearson correlation coefficients between density estimates derived using varying numbers of landmarks and the landmark-free full Gaussian process for each dataset. In the case of the iPSC dataset, a 65k landmark inference is used as the reference, as the full Gaussian process requires too much memory. A vertical line marks the default value of 5,000 landmarks, while a horizontal line denotes a perfect Pearson correlation coefficient of 1.

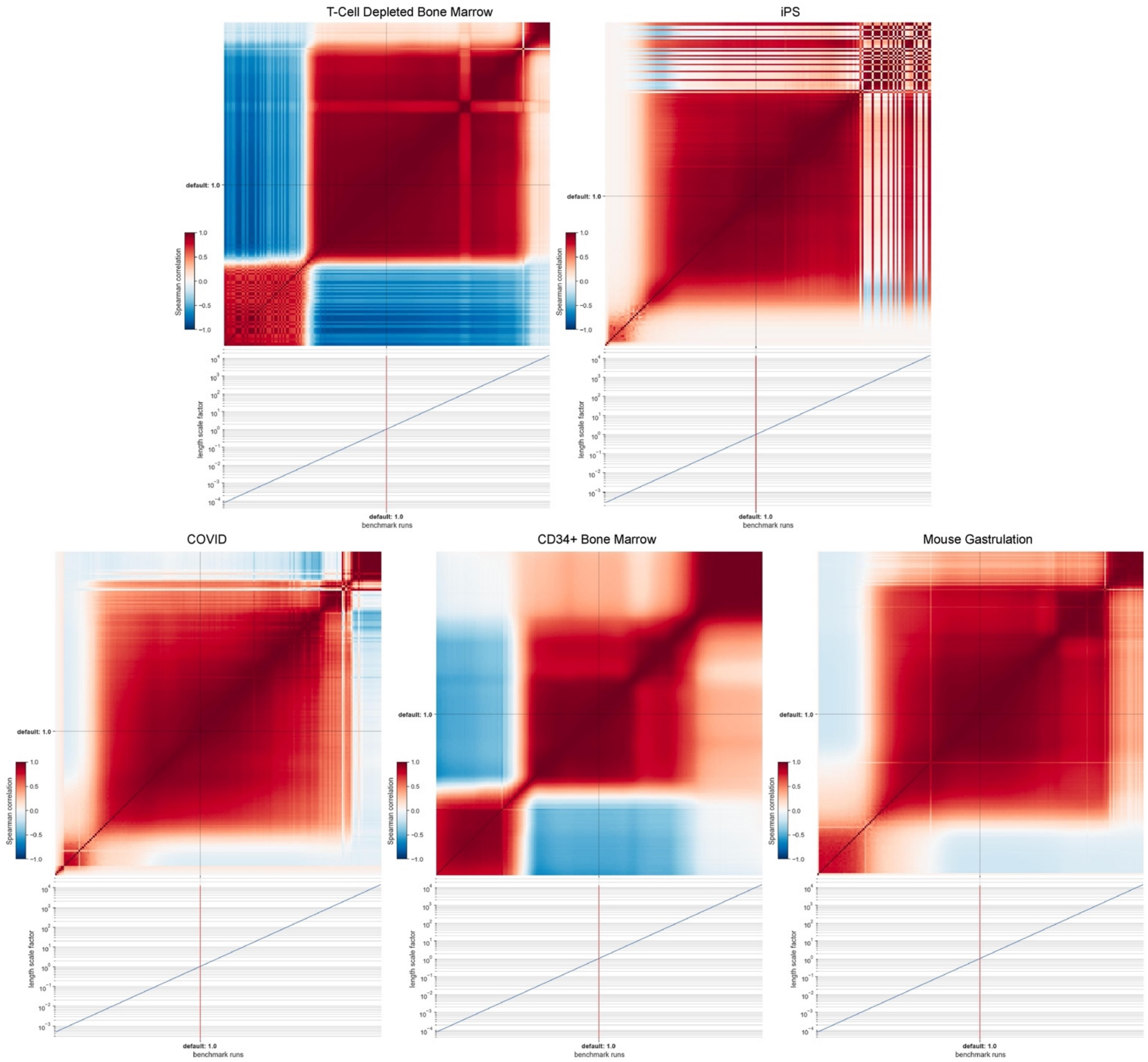

**Supplementary Figure 16: Mellon densities are robust to varying the length scale factor for the covariance function.**

Heatmaps displaying Spearman correlation between density estimates obtained using different length scale factors for five datasets. The length scale factor employed for each column's density estimate is denoted in a marginal plot at the bottom. The length scale factor varies from smaller to larger values from right to left. The default value of 1, corresponding to the utilization of the length scale as derived from the heuristic (**Methods**), is marked by a vertical line running through both the heatmap column and the marginal plot. The colormaps are consistent across all heatmaps, spanning from -1 to 1, encapsulating the full potential range of Spearman correlation values. High correlation values around the default value suggest that Mellon's density inference remains stable even when the length scale factor deviates slightly from the heuristic-derived value.

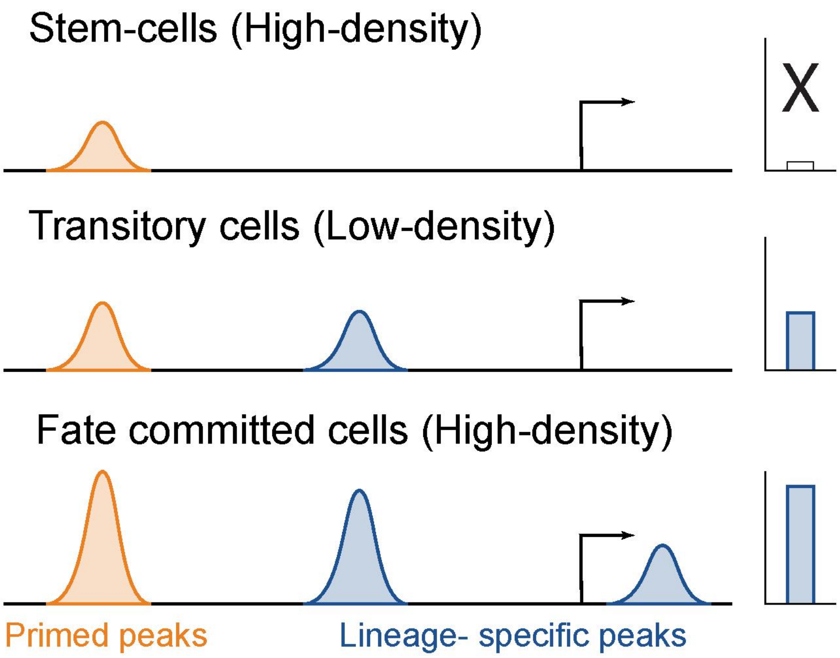

**Supplementary Figure 17**

Schematic illustrating the role of primed and lineage-specific peaks in achieving rapid transcriptional changes. Rows represent gene loci in different cell-states. Top: Primed peaks are pre-established in stem cells without turning on gene expression. Expression of the gene is upregulated with the emergence of lineage specific peaks (Middle, bottom).

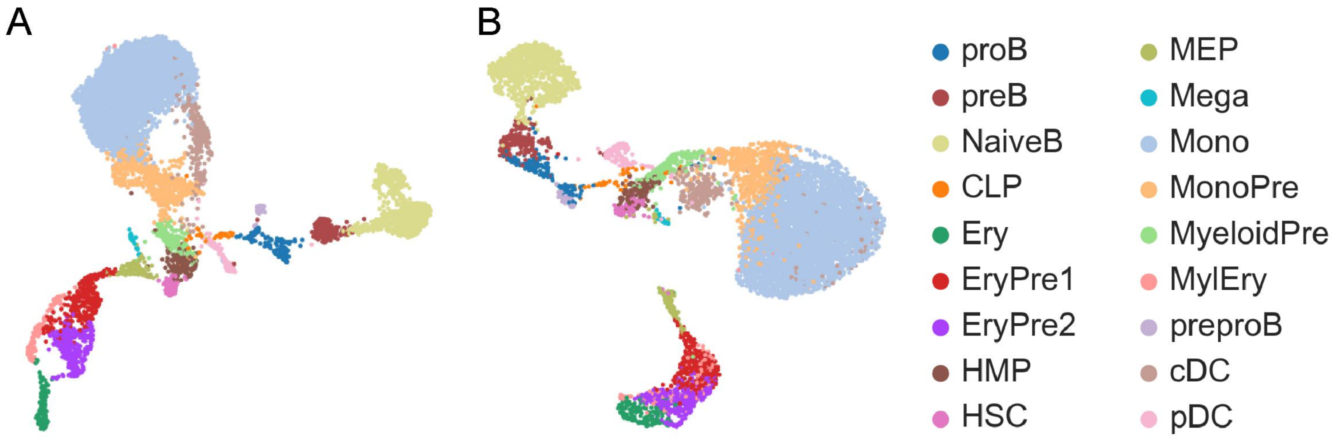

**Supplementary Figure 18: T cell depleted bone marrow multiome dataset.**

A. Cell-type annotations for all cells in the T cell depleted bone marrow dataset (8627 cells) plotted on the RNA UMAP embedding.

B. Identical cell type annotations to A for all cells in the dataset, plotted on the ATAC UMAP embedding.

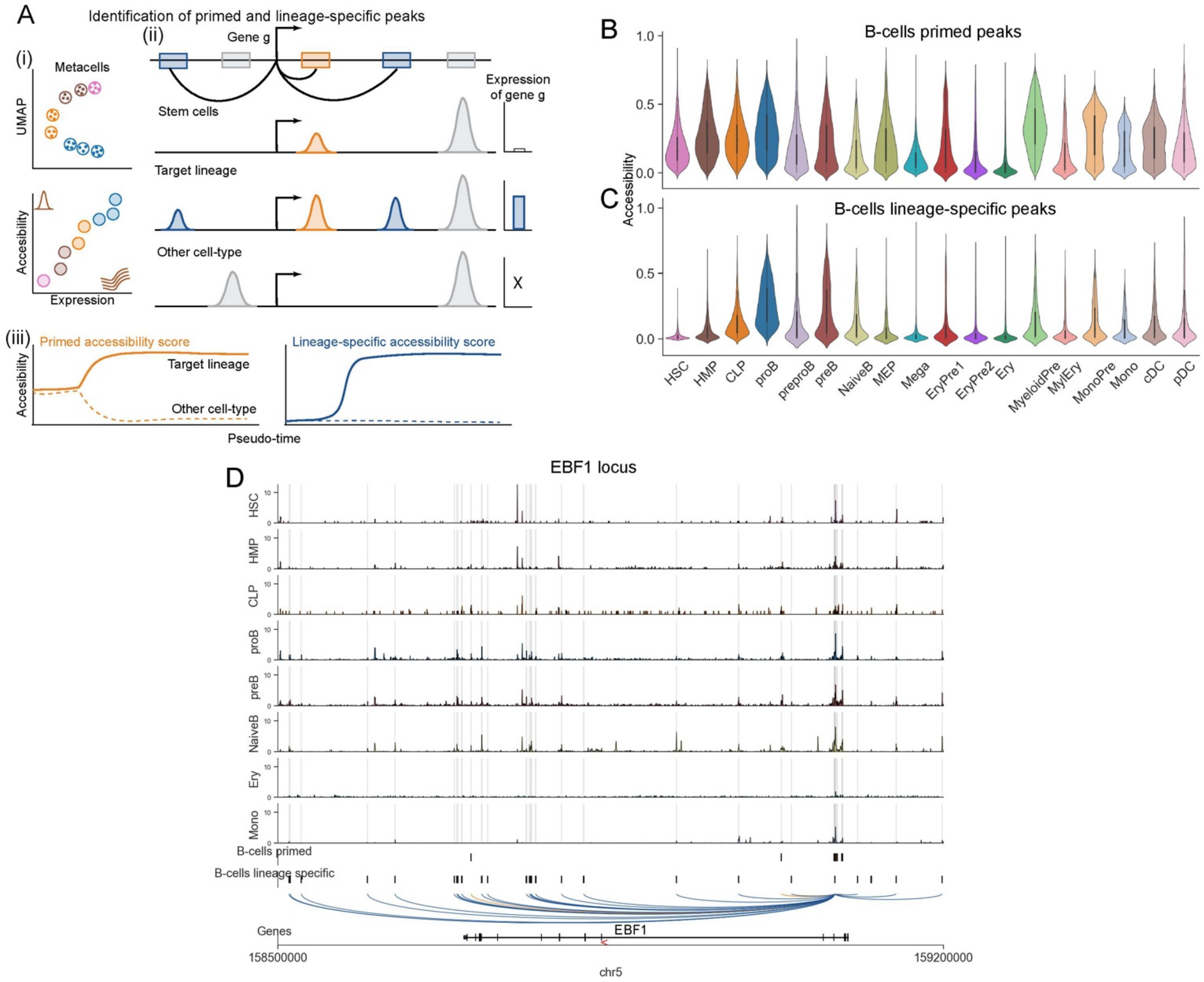

**Supplementary Figure 19: B-cell primed and lineage specific peaks.**

A. Schematic illustrating the approach to identify primed and lineage specific peaks. (i) Metacells are used to identify peaks with accessibility that significantly correlate with gene expression. (ii) Correlated peaks with greater accessibility in the target lineage compared to other cell-types are nominated as primed if they are accessible in stem cells (Orange) and lineage-specific other-wise (Blue). (iii) Accessibility of peaks associated with each gene are summarized to derive primed and lineage-specific scores. Dynamics in other lineages are shown as dotted lines.

A. Plots showing the number of peaks significantly correlated with each gene. The correlations were computed using SEACells[7] metacells.

C. Violin plots determined using averaged imputed accessibility for B-cell primed peaks.

D. Same as (C), for B-cell lineage specific peaks.

E. Coverage plots highlighting the EBF1 correlated peaks. Primed peaks are in orange and lineage specific peaks are in blue.

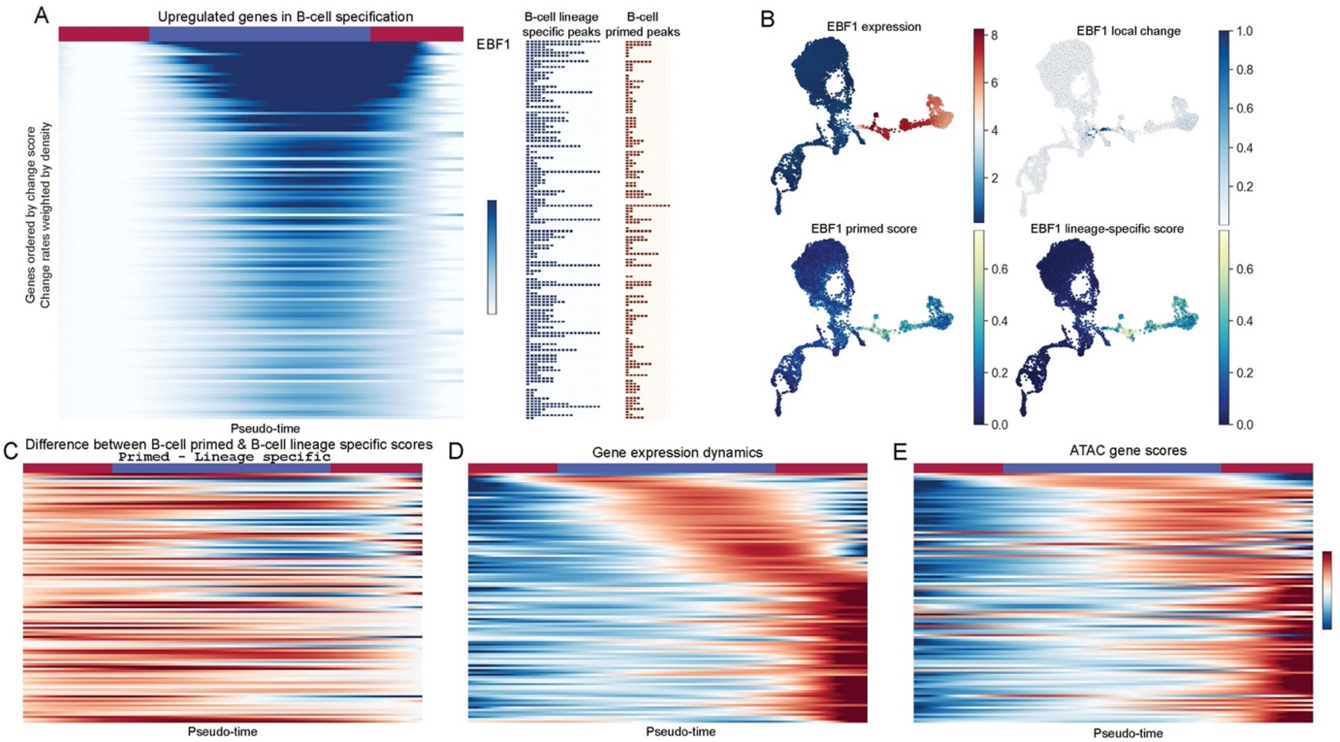

**Supplementary Figure 20: B-cell primed and lineage specific peaks.**

A. Left: Heatmap showing the trend of local expression variability during the trajectory of B-cell specification (**Fig. 3B**). Upregulated genes in the top 5$^{th}$ percentile of B-cell specification gene change scores are shown. Genes are ordered by change scores. EBF1 has the highest change score.

Middle: Plots depicting the number of B-cell lineage specific peaks associated with each gene.

Right: Plots depicting the number of B-cell primed peaks associated with each gene.

B. UMAPs colored by EBF1 MAGIC imputed expression, local variability, primed accessibility score and lineage-specific accessibility scores (**Related to Fig. 3C**).

C. Heatmap of the difference in normalized B-cell primed and lineage-specific accessibility trends for genes associated with density change in B-cell specification (**Related to Fig. 3E-F**). Stronger red signal at the start of the trajectory indicates primed enhancers are associated with the gene.

D. Heatmap of gene expression dynamics along B-cell specification. Genes are colored by expression along trajectory.

E. Heatmap of ArchR[8] gene scores derived from the ATAC modality. Genes are in the same order as (D).

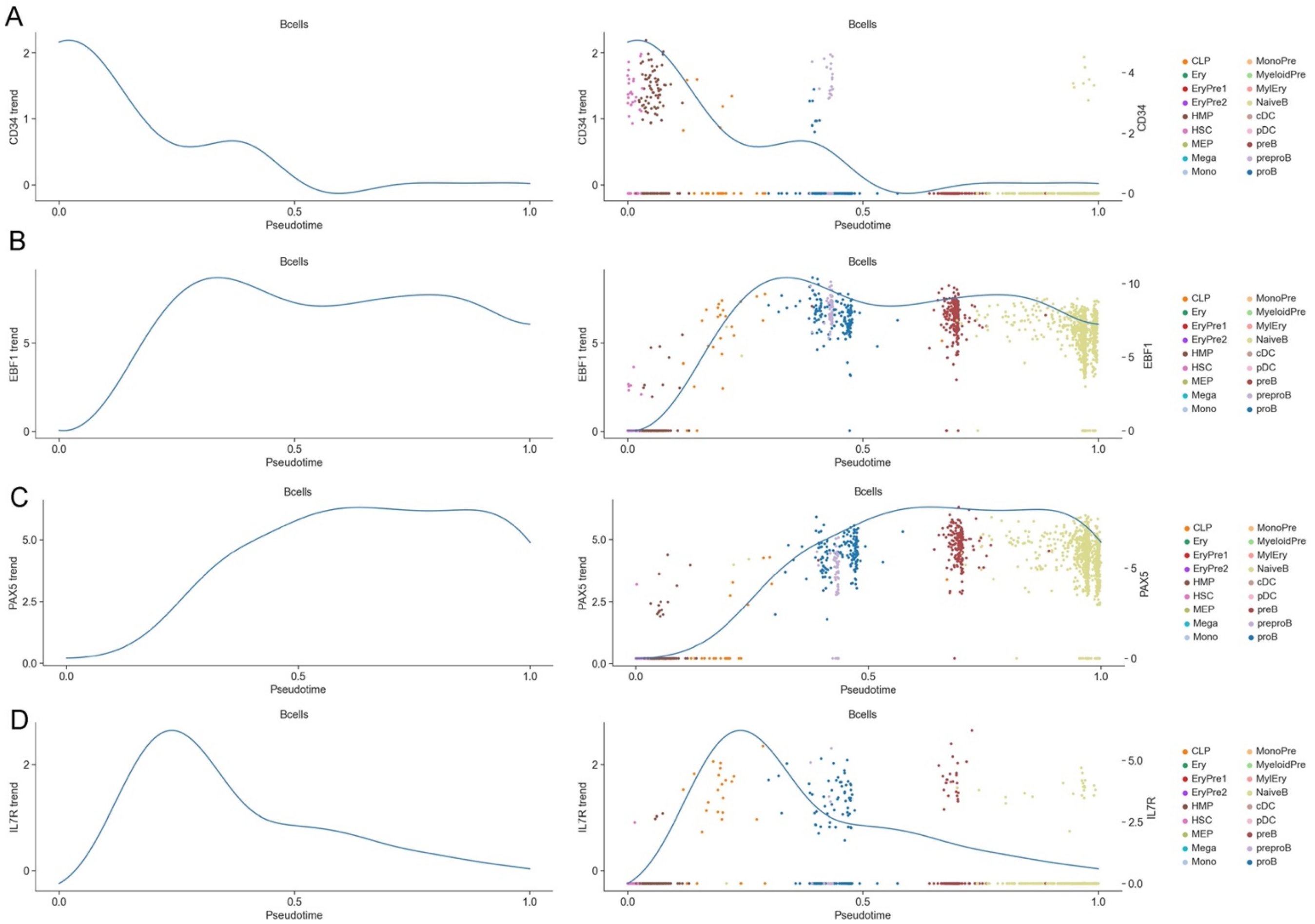

**Supplementary Figure 21: Gene trend computation using Mellon**

A. Left: Expression trend for CD34 computed along B-cell lineage using Mellon trend estimator. Right: Expression trend for CD34 along with the normalized, log-transformed data used for computing the trend. B – D. Same as (A) for EBF1 (B), PAX5 (C) and IL7R (D).

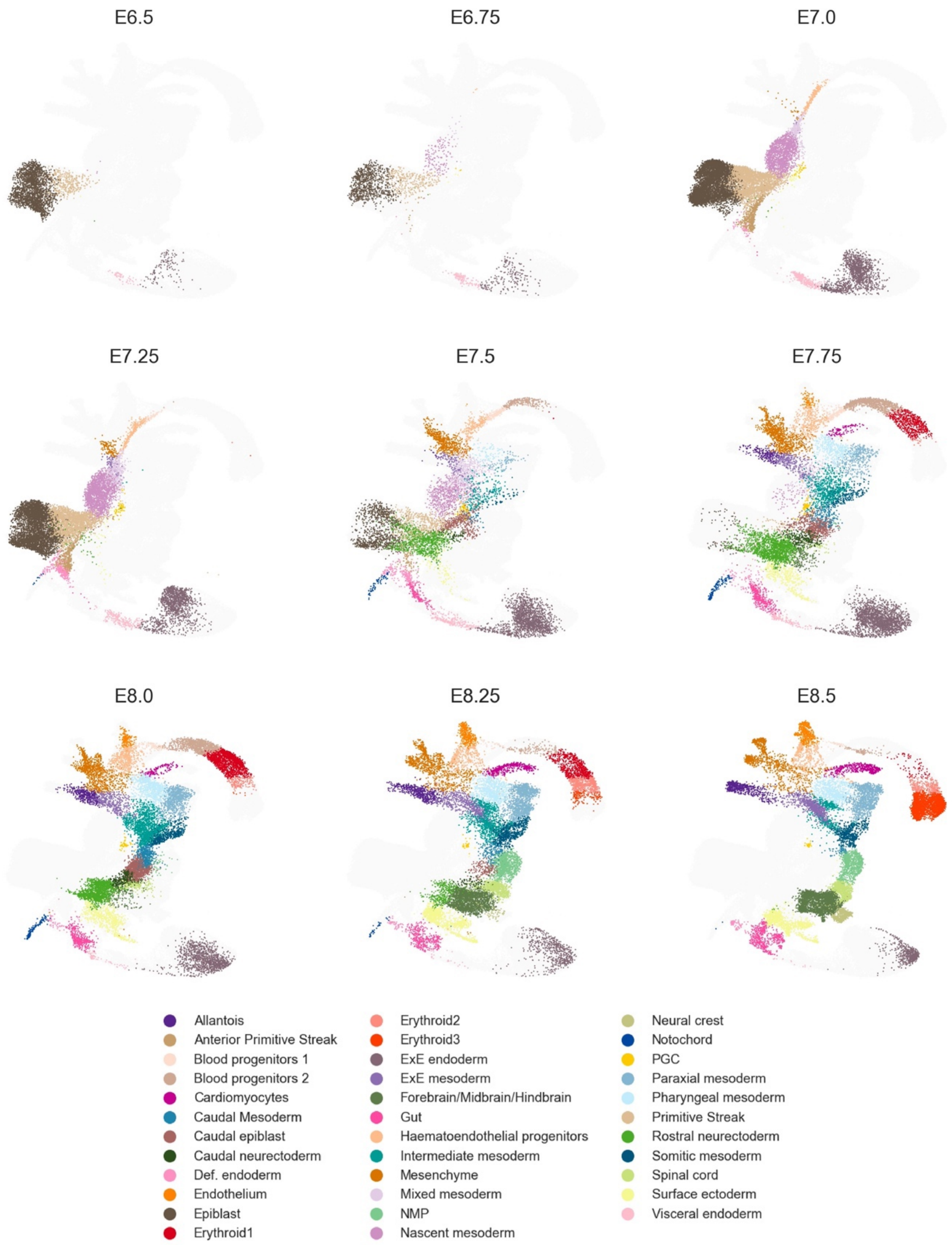

**Supplementary Figure 22: Cell-type composition in mouse gastrulation scRNA-seq dataset.** UMAPs colored by cell-types of cells per timepoint for the mouse gastrulation atlas[9]. Cells from other time points are displayed in a faint grey to maintain context.

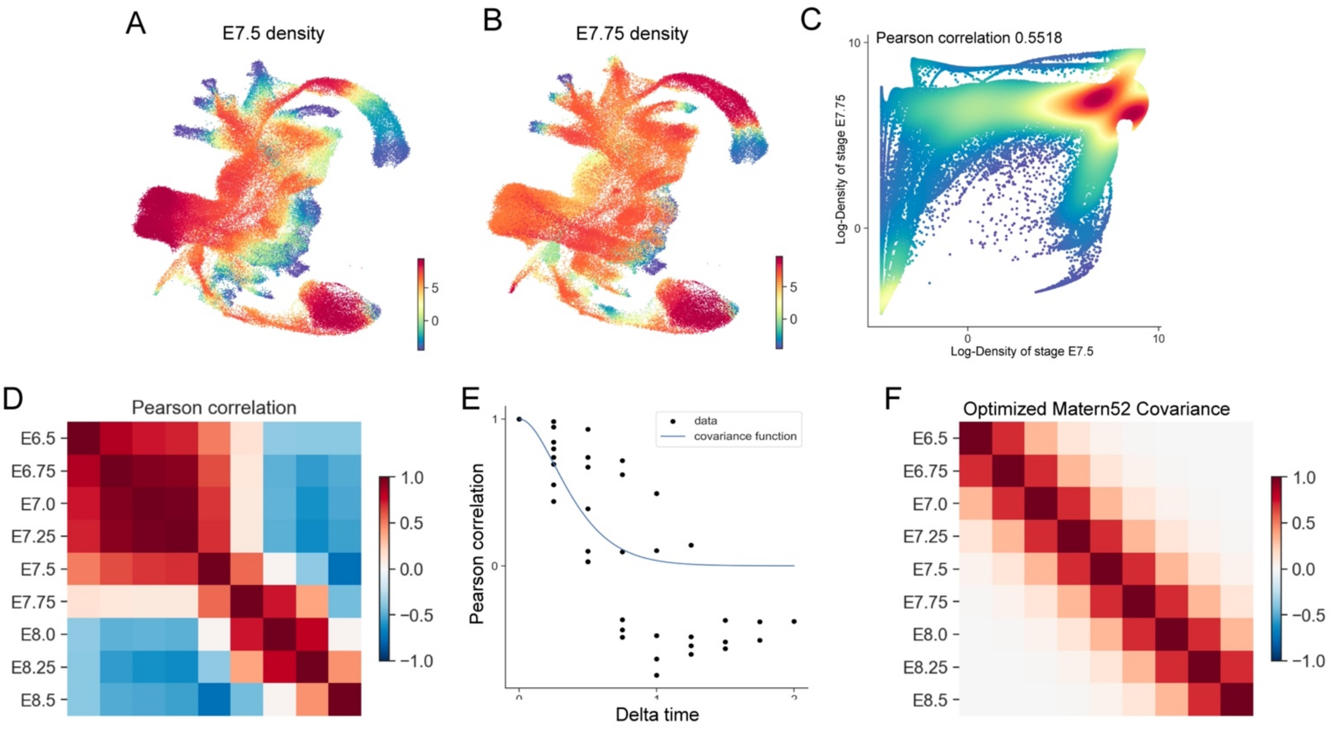

**Supplementary Figure 23: Cell-state density correlation across timepoints during mouse gastrulation.**

A. UMAP of all cells in the mouse gastrulation dataset colored by Mellon-derived density for cells at E7.5.

B. Same as (A), showing density for cells at E7.75.

C. Correlation plot comparing densities at stages E7.5 (x-axis) and E7.75 (y-axis).

D. Pearson correlation matrix illustrating density correlation between all combinations of time points.

E. Plot of Pearson correlation for all time point pairs (y-axis) against their temporal difference (x-axis), represented as black dots. The covariance of the optimized Matern52 length scale over time distances is displayed as a blue line.

F. The covariance matrix produced by the Matern52 kernel (from E) based on the temporal differences between sample pairs.

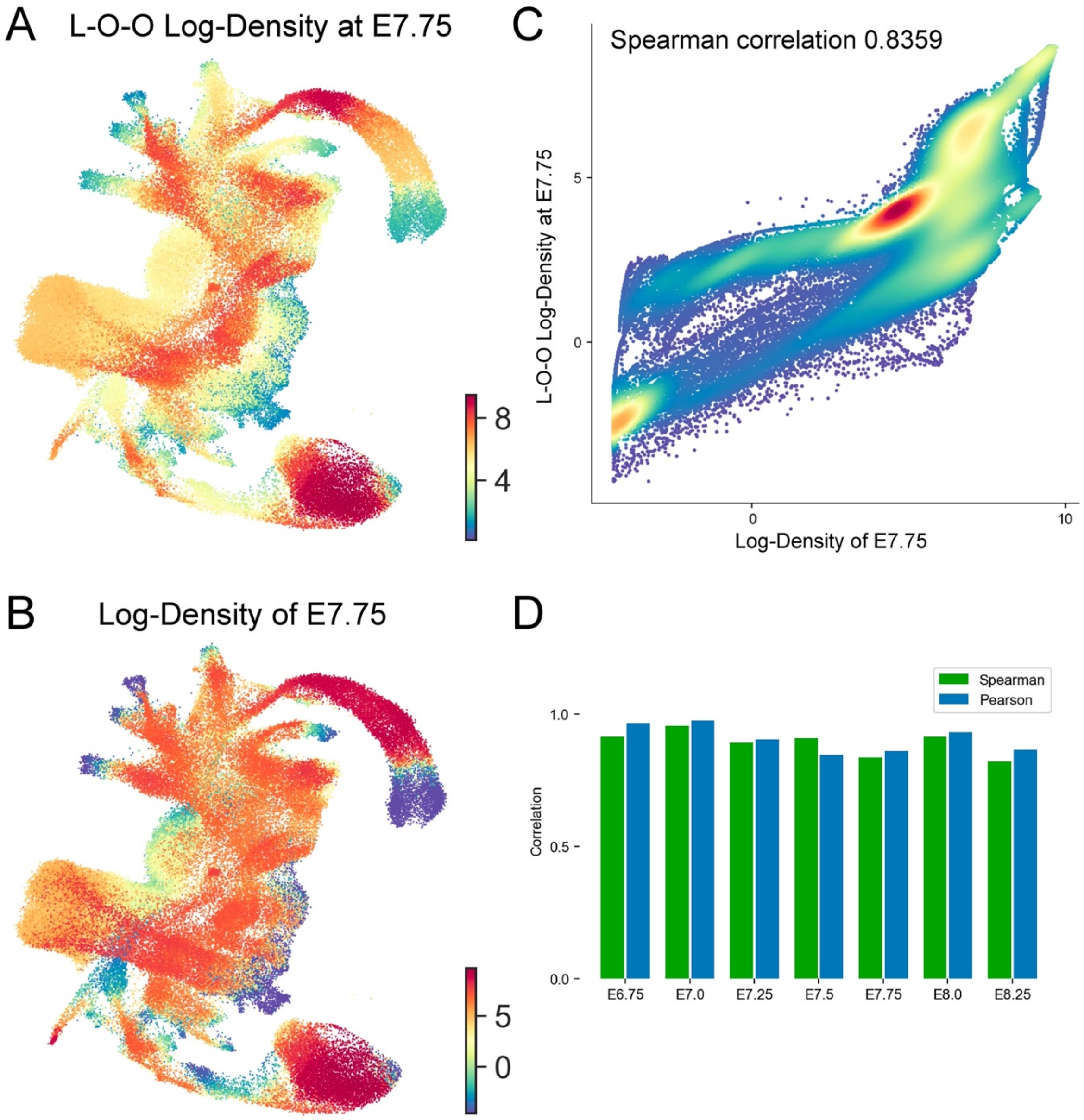

**Supplementary Figure 24: Evaluation of Mellon's time-continuous density via leave-one-out cross-validation.**

A. A representative slice at time point E7.75 from Mellon's time-continuous density, inferred excluding E7.75 cells.

B. UMAP colored by Mellon-derived density using only cells from timepoint E7.75.

C. Correlation plot comparing the density at E7.75 (from (B)) and the time-continuous density slice from the leave-one-out method (from (A)).

D. Bar plot indicating both Pearson and Spearman correlations for all leave-one-out tests; each test omits one timepoint and compares the resulting time-slice at this omitted timepoint to the density of cells specifically from this timepoint.

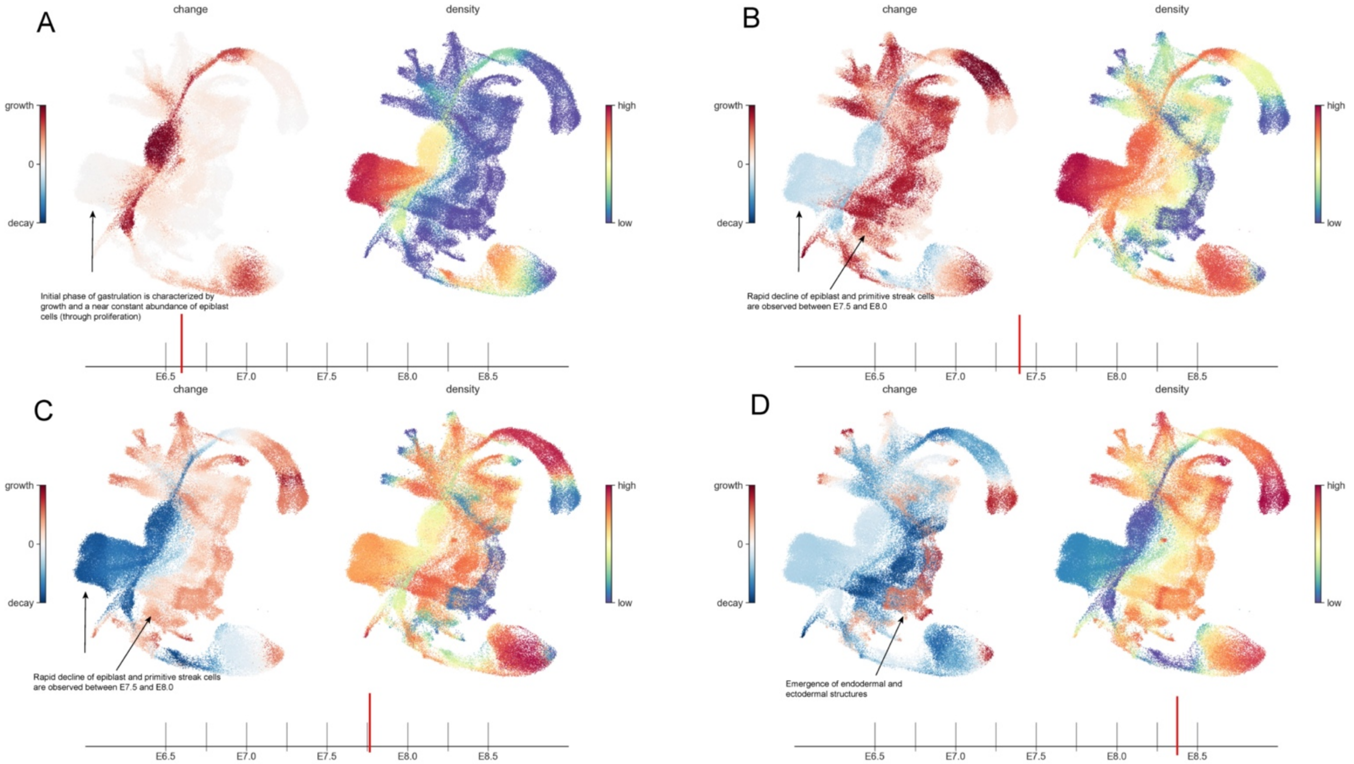

**Supplementary Figure 25**

Select frames from the animation presented in **Supplementary Video 1**, illustrating the time-continuous density and its derivative at four randomly chosen time points. Each frame is divided into a right and left panel, with the right panel showcasing the cell density across all cell states for the specified time point, and the left panel exhibiting the time derivative of this density. Red regions denote cell enrichment over time, while blue regions indicate cell depletion. The time point corresponding to each frame is highlighted on the timeline shown below the panels.

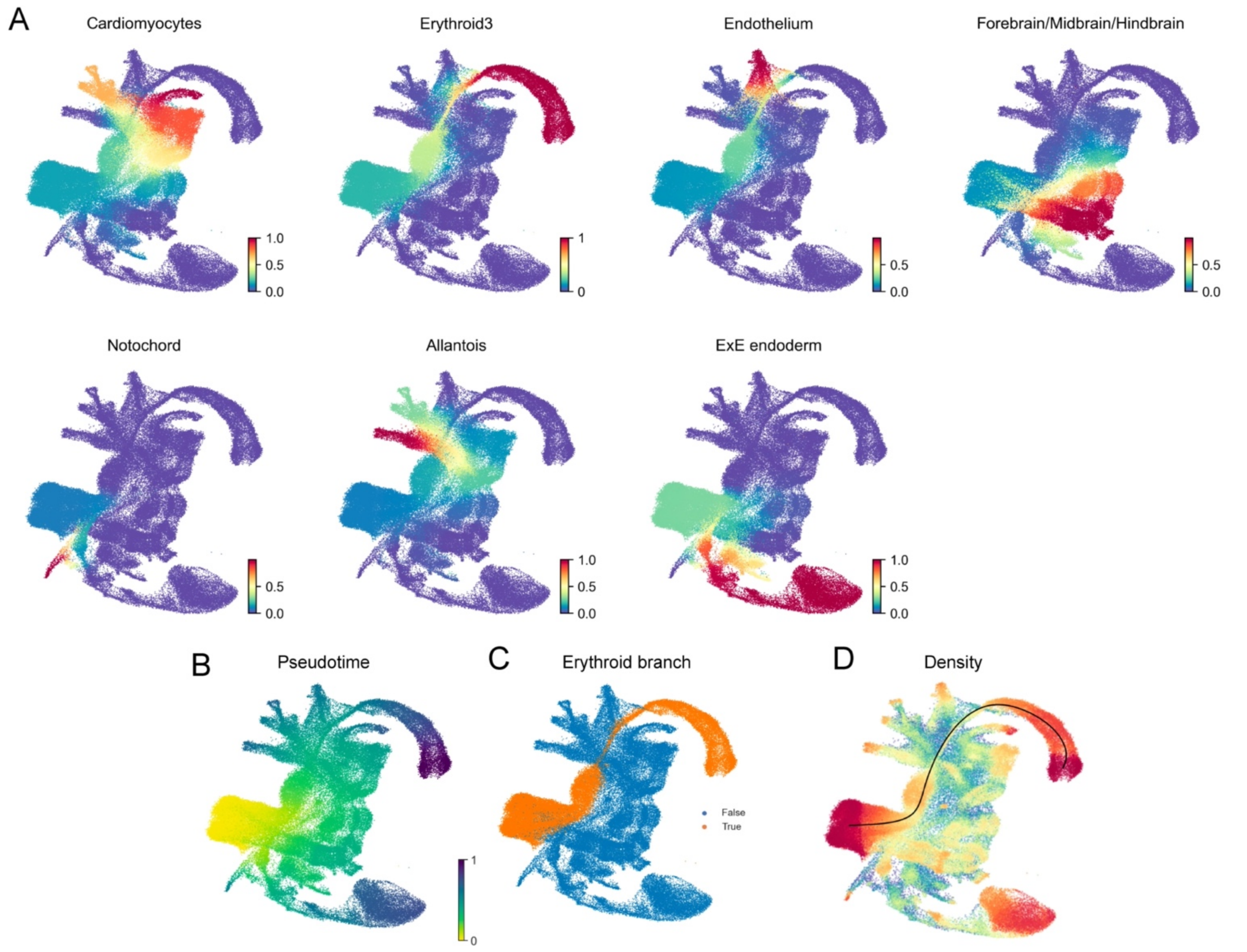

**Supplementary Figure 26: Depiction of the selected Erythroid developmental trajectory within the mouse gastrulation dataset.**

UMAPs colored by fate probabilities for terminal cell states (A) and pseudotime (B) as determined by Palantir.

C. Cells chosen to represent the Erythroid branch, extending from early epiblasts to erythroid cells.

D. UMAP colored by time-agnostic density across the UMAP. Erythroid trajectory is represented by a black line.

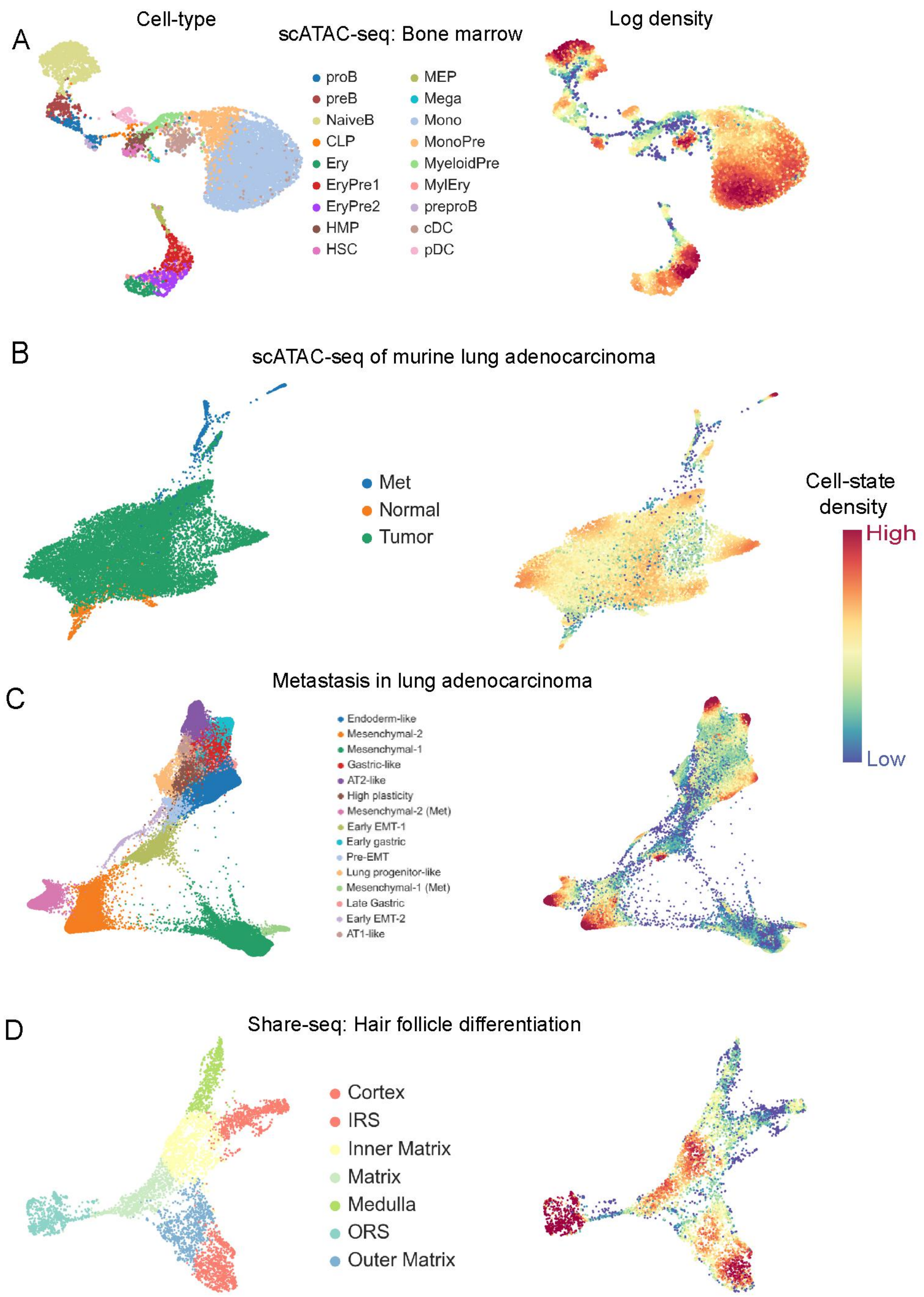

**Supplementary Figure 27: Mellon generalizes to different single-cell data and cell-state representations**

A. UMAPs colored by cell-type (left) and density (right) for scATAC-seq modality of T-cell depleted bone marrow dataset[7].

B Force directed layout of the mouse lung adenocarcinoma scATAC-seq dataset[10] colored by cell-type (left) and Mellon density (right).

C. Same as (A), for the scRNA-seq data of mouse models of lung adenocarcinoma[11]. UMAPs and diffusion maps for density estimation were computed using scVI[12] latent space.

D. Same as (A), for SHARE-seq data of mouse skin differentiation[13]. UMAPs and diffusion maps for density estimation were computed using MIRA[14] multimodal representation.

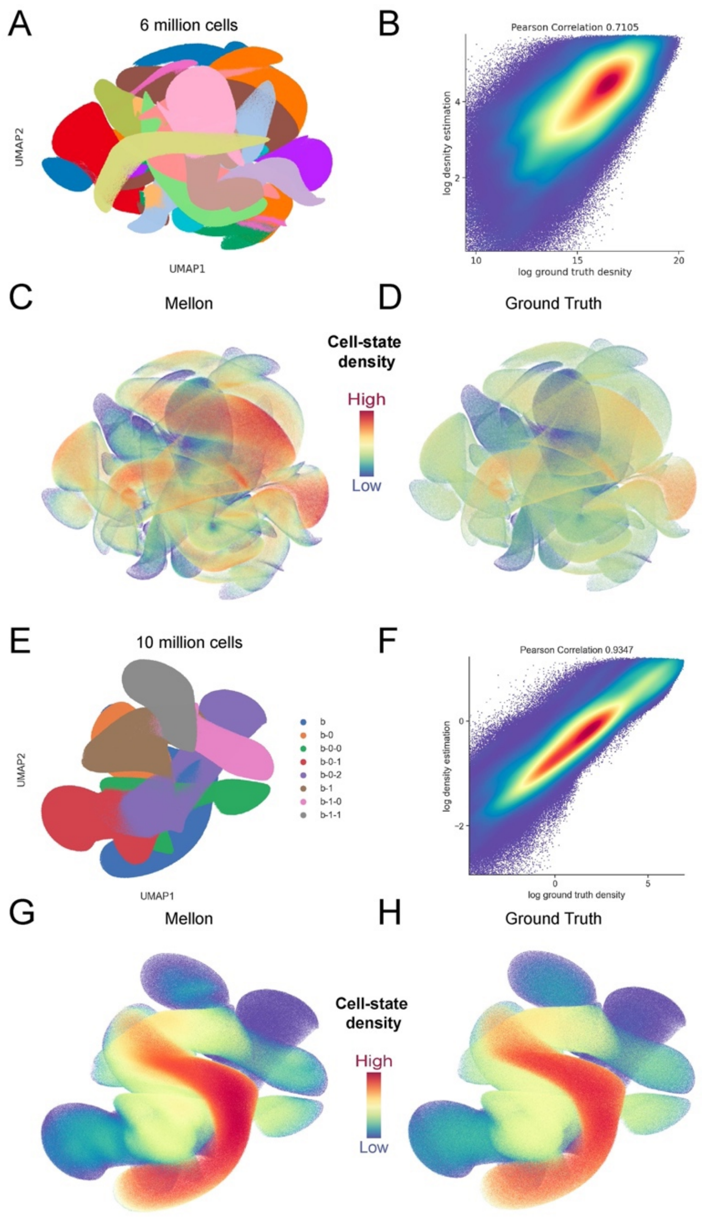

**Supplementary Figure 28: Illustration of Mellon's scalability utilizing large, simulated datasets.** (A-D) Density inference in a dataset with 6 million cells, structured based on a differentiation tree, using the default 5000 landmarks. (E-H) Density inference in a larger dataset with 10 million cells, structured based on a differentiation tree, using a reduced number of 2000 landmarks for improved scalability.

Each dataset features four subplots, organized akin to **Supplementary Figure 4**. (A and E) UMAPs of the simulated data display color coding according to the differentiation tree nodes. (B and F) Correlation plots demonstrate the relationship between known ground truth log-density (x-axis) and Mellon-inferred density (y-axis) across all simulated cells. Each dot represents an individual simulated cell. (C and G) UMAPs are colored based on Mellon-inferred density. (D and H) UMAPs are colored according to ground truth log-density. Density values below the 20th percentile in C, D, G, and H are adjusted to the 20th percentile level to enhance visualization clarity.

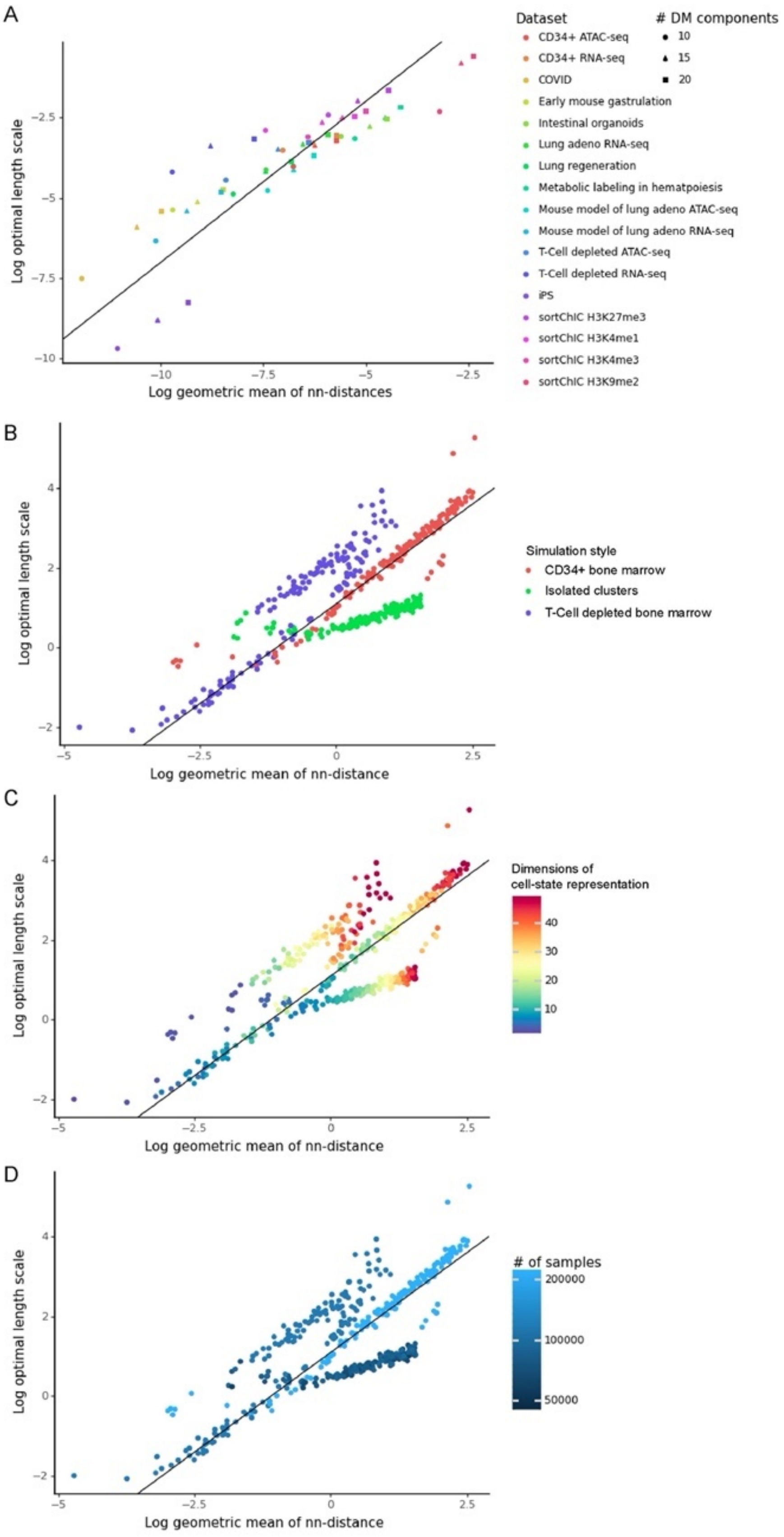
A
B
C
D
Dataset
### DM components
10
15
20
CD34+ ATAC-seq
CD34+ RNA-seq
COVID
Early mouse gastrulation
Intestinal organoids
Lung adeno RNA-seq
Lung regeneration
Metabolic labeling in hematpoiesis
Mouse model of lung adeno ATAC-seq
Mouse model of lung adeno RNA-seq
T-Cell depleted ATAC-seq
T-Cell depleted RNA-seq
iPS
sortChIC H3K27me3
sortChIC H3K4me1
sortChIC H3K4me3
sortChIC H3K9me2
Log optimal length scale
Log geometric mean of nn-distances
Simulation style
CD34+ bone marrow
Isolated clusters
T-Cell depleted bone marrow
Log geometric mean of nn-distance
Dimensions of
cell-state representation
### of samples
200000
100000
50000

**Supplementary Figure 29: Mellon length scale heuristic.**

A. Plot depicting datasets with varying numbers of diffusion components used for cell-state representation (**Supplementary Table 1**). Optimal length scale (y-axis) is chosen through the maximum a posteriori estimate of the Bayesian model employed in Mellon's density inference, where the length scale is considered a free parameter. Regression line depicts the fit used to derive the length scale heuristic for any dataset.

B-D. Scatter plots displaying the same relation for 597 simulated datasets (**Supplementary Note 5**). The optimal length scale on the y-axis is chosen by maximizing the Spearman correlation of the Mellon-inferred density with the ground truth density. The x-axis shows the geometric mean of nearest neighbor distances across all cells in the respective datasets. (B) Datasets colored by the simulation style as per **Supplementary Note 5**. (C) Datasets colored by dimension of the cell-state space. (D) Datasets colored by the number of cells in the dataset. Regression line is derived from (A), reinforcing the relationship between nearest neighbor distances and the optimal length scale.

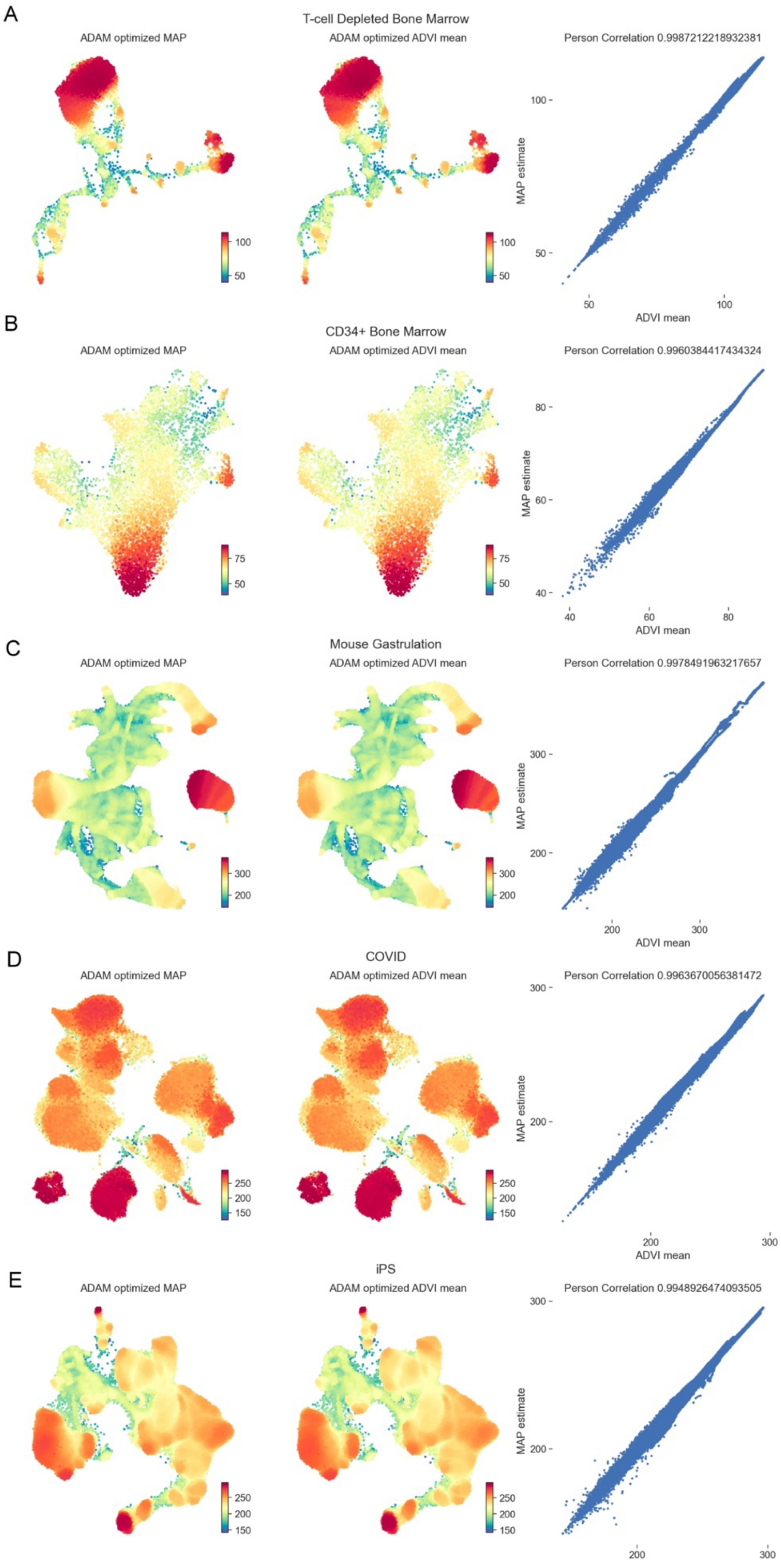
A
T-cell Depleted Bone Marrow
ADAM optimized MAP
ADAM optimized ADVI mean
Person Correlation 0.9987212218932381
MAP estimate
ADVI mean
B
CD34+ Bone Marrow
ADAM optimized MAP
ADAM optimized ADVI mean
Person Correlation 0.9960384417434324
MAP estimate
ADVI mean
C
Mouse Gastrulation
ADAM optimized MAP
ADAM optimized ADVI mean
Person Correlation 0.9978491963217657
MAP estimate
ADVI mean
D
COVID
ADAM optimized MAP
ADAM optimized ADVI mean
Person Correlation 0.9963670056381472
MAP estimate
ADVI mean
E
iPS
ADAM optimized MAP
ADAM optimized ADVI mean
Person Correlation 0.9948926474093505
MAP estimate

**Supplementary Figure 30: Consistency of MAP estimate with the ADVI mean of the posterior distribution of the density in Mellon's Bayesian model for density inference.**

Results are shown for five datasets in A-E

Left: MAP estimate of log-density as inferred by the ADAM stochastic optimizer.

Middle: Mean of the posterior distribution for the density function, inferred by ADVI using the ADAM stochastic optimizer.

Right: Correlation plots between the ADVI mean (x-axis) and the MAP (y-axis), highlighting the congruence between the two methods.

MAP: Maximum a Posteriori; ADVI: Automatic Differentiation Variational Inference.

Note that all results (ADVI mean, ADAM-optimized MAP, and L-BFGS-B optimized MAP) are derived from Mellon's inbuilt functionality.

**Supplementary Video 1: Animation demonstrating the progression of time-continuous density across the entire range of time covered by the dataset.**

The video displays a UMAP plot where the right panel presents the density for a specific time point evaluated across all cells, and the left panel depicts the time derivative of this density. Red signifies an enrichment of cells over time, while blue indicates depletion. A red vertical line on the timeline at the bottom marks the current time for the displayed frame, while black vertical lines denote the measured timepoints.
