## Supplementary Notes for "Quantifying Cell-State Densities in Single-Cell Phenotypic Landscapes using Mellon"

### Supplementary Note 1

### Nearest Neighbor Distribution

The Nearest Neighbor Distribution (NND) provides the basis for the density estimation in our model. For a homogeneous Poisson point process on $\mathbb{R}^{d'}$ with a constant density $\rho$, the cumulative probability density function $D : \mathbb{R}^+ \to [0, 1]$ of distances to the nearest data point from any point $r$ in $\mathbb{R}^{d'}$ is given by:

$$D(r) = 1 - \exp(-\rho \cdot b(r, d)),$$

where $b(r, d)$ is the volume of the $d$-dimensional ball with radius $r$. The corresponding probability density function (pdf) $f_{\text{NN}}\colon \mathbb{R}^+ \to \mathbb{R}^+$ is then given by:

$$f_{\text{NN}}(r|\rho) = \frac{\mathrm{d}D}{\mathrm{d}r} = \exp(-\rho \cdot b(r, d)) \cdot \rho \frac{\mathrm{d}b(r, d)}{\mathrm{d}r}.$$

Taking into account the relationship

$$\frac{\mathrm{d}b(r, d)}{\mathrm{d}r} = \frac{d \cdot r^{d-1} \cdot \pi^{\frac{d}{2}}}{\Gamma(\frac{d}{2} + 1)},$$

the pdf for the nearest neighbor distribution $\text{NN}(\rho, d)$ with fixed density $\rho$ can be expressed as:

$$\begin{aligned} \log \circ f_{\text{NN}}(r|\rho) =& \log(\rho) - \frac{\rho \cdot r^d \cdot \pi^{\frac{d}{2}}}{\Gamma(\frac{d}{2} + 1)} + \log(d) + \frac{d}{2} \log(\pi) \\ &+ (d - 1) \cdot \log(r) - \log \circ \Gamma \left( \frac{d}{2} + 1 \right). \end{aligned} \tag{1.1}$$

In our model, the density $\rho$ is not constant, but is instead a function $\rho\colon \mathbb{R}^{d'} \to \mathbb{R}^+$. Consequently, to apply the above reasoning, we must assume that the density at any point $x \in \mathbb{R}^{d'}$ approximates the average density within a ball $b(x, \text{dn}(x))$, where $\text{dn}(x)$ is the distance to the nearest neighbor of $x$. This assumption allows us to connect local density with the nearest neighbor distance distribution within our model's framework.

The pdf leads to a maximum likelihood point estimate for a given $\text{dn}(x)$ through the equation:

$$\frac{\mathrm{d} \log \circ f_{\text{NN}}(r|\rho)}{\mathrm{d}r} = 0,$$

resulting in:

$$\hat{\rho}(\text{dn}(x)|d) = \frac{(d - 1) \cdot \Gamma(\frac{d}{2} + 1)}{d \cdot \text{dn}(x)^d \cdot \pi^{\frac{d}{2}}}, \tag{1.2}$$

which serves as the heuristic maximum likelihood estimate for the density at a point $x$, when no prior for the density function is employed.

In our model, we set $d := d'$, a simplification made under the assumption that cell-states can vary in all directions with equal likelihood. However, a more precise definition of $d$ would reflect the intrinsic dimension of the local subspace of state variability. This is typically lower than the dimensionality of the embedding space, as cell states are restricted to the phenotypic manifold, meaning that the local subspace of variability is the tangent space. However, it's worth mentioning that our analysis, as demonstrated in **Supplementary Figure 14**, shows that the density inference remains stable with respect to Spearman correlation even when this parameter is varied. As a result, any variations in this parameter don't significantly influence the conclusions drawn from relative density changes within the dataset.

### Supplementary Note 2

### Gaussian Process

In order to sample from $\mathrm{GP}(m, k_l)$ for some kernel function $k_l\colon \mathbb{R}^{d'} \times \mathbb{R}^{d'} \to \mathbb{R}$ we employ a sparse gaussian process. The term

$$f \sim \mathrm{GP}(m, k_l)$$

in the model translates to

$$f \sim \mathcal{N}(m, \Sigma)$$

where $\Sigma$ is the covariance matrix induced by the kernel function $\Sigma_{i,j} := k_l(x_i, x_j)$. While Mellon accepts any covariance function, we chose the Matern kernel with $\nu = \frac{5}{2}$ as default:

$$k_l(x_i, x_j) = \mathrm{Matern}_l \frac{5}{2}(x_i, x_j) = \left(1 + \frac{\sqrt{5}h}{l} + \frac{5h^2}{3l^2}\right) \exp\left(-\frac{\sqrt{5}h}{l}\right)$$

where $h = \|x_i - x_j\|_2$. To sample from this distribution we draw samples from

$$y \sim \mathcal{N}(0, \mathbb{I}) \tag{2.1}$$

where $\mathbb{I} \in \mathbb{R}^{k \times k}$ is the identity matrix and transform $y$ with

$$yL + m \sim \mathcal{N}(m, \Sigma)$$

for some matrix $L \in \mathbb{R}^{k \times d}$. To speed up the inference we employ a dimensional reduction where $k < d$ and $yL + m \sim \mathcal{N}(m, \Sigma')$ such that $\Sigma' \approx \Sigma$. This means

$$LL^T = \Sigma' \tag{2.2}$$

and the rank of $\Sigma'$ is at most $k$. To construct the desired $L$, we utilize a set of landmark points, also known as inducing points, to formulate the distribution of $f$ as conditional on $y$. A detailed explanation of this approach is provided below.

#### 2.1 Using Landmark Cell States in Sparse GP

To reduce computational demands we implement a Sparse Gaussian process using inducing points that we refer to as landmark cell states. Let $f' \in \mathbb{R}$ be the values of some landmark cells $x' \in \mathbb{R}^{d'}$ then

$$\begin{bmatrix} f - m \\ f' - m \end{bmatrix} \sim \mathcal{N}\left(0, \begin{bmatrix} \Sigma_{f,f} & \Sigma_{f,f'} \\ \Sigma_{f',f} & \Sigma_{f',f'} \end{bmatrix}\right) = p(f, f') = p(f|f') \cdot p(f')$$

which gives rise to the conditional distribution

$$p(f - m | f' - m) = \mathcal{N}(\Sigma_{f,f'}\Sigma_{f',f'}^{-1}(f' - m), \Sigma_{f,f} - \Sigma_{f,f'}\Sigma_{f',f'}^{-1}\Sigma_{f',f}).$$

As our interest lies solely in the maximum a posteriori estimate of $f$, we will disregard the covariance of this conditional distribution and instead focus on computing its mean with respect to $f'$. After the Cholesky decomposition of $\Sigma_{f',f'} = UU^T$, and (2.1) we have

$$f' - m = Uy \sim N(0, \Sigma_{f',f'})$$
$$\Sigma_{f',f}U^{-1}y \sim \mathcal{N}(0, \Sigma').$$

This gives us $L := \Sigma_{f',f}U^{-1}$ with a rank that is limited by the number of landmarks $k$. Note that

$$LL^T = \Sigma' = \Sigma_{f,f'}\Sigma_{f',f'}^{-1}\Sigma_{f',f}. \tag{2.3}$$

The time complexity to compute $L$ and the subsequent inference is $o(nk^2)$ where $n$ is the number of cells.

### Supplementary Note 3

### The Gaussian Process Mean Function

The random variable of the Gaussian process used in our model serves as an approximation of the logarithm of the density of cell states. We know that the true log-cell-state density approaches negative infinity away from any observed cell state. However, functions sampled by the Gaussian process approach the chosen mean. To approximate the true behavior of density functions we choose a very small value for the mean $m$ that implies vanishingly small probability for a distant cell state. To find a reasonably low value we compute the expected distribution of density values for a given dataset through the maximum likelihood log-density estimations (MLDE) for each individual cell (1.2). While using the minimum observed MLDE value could serve as a reasonably small value, we opted against this to avoid dependence on individual cells and risk instability of the density inference. Instead, we chose the 1% quantile of MLDE values

$$q_{1\%} := P_{1\%}\left[\hat{\rho}(\mathrm{dn}(x_i)|d)_{i\in\{1\ldots n\}}\right]$$

and subtracted a blanket 10 from this value which translates to a factor $4.5 \cdot 10^{-5}$ to the low density value in linear scale. This mean

$$m = q_{1\%} - 10$$

is as a "very low" log-density value and serves as a minimum, the inferred log-density converges to away from any observed cell state.

If convergence towards negative infinity is required one can transform the inferred log-density function $f$ through

$$f'(x_i) := \log\left[\exp \circ f(x_i) - \exp(m)\right].$$

### Supplementary Note 4

### The Covariance Length Scale

Updating the length scale $l$ throughout the inference is an expansive process since it requires updating the covariance matrices $\Sigma_{\cdot,\cdot}$ and there decompositions. Therefore, we investigated the dependency between (i) the mean $n^{th}$ nearest neighbor distance between cells, and (ii) the maximum a posterior length scale, which regard as optimum acorss multiple datasets.

To find the maximum a posteriori estimation of the length scale, we ran the density inference for a grid of values ranging from the smallest nearest neighbor distance in the sample and the prism diagonal of the dataset which serves as a upper bound for the distance between any two cells. The distance between neighboring length scales in the grid is at most 20% of the larger one. We show in the **Supplementary Fig. 16** that the density result remains stable within a factor 2 of the best optimal length scale. Hence the chosen resolution is guaranteed to include a length scale that produces an optimal result.

The correlation plots in **Supplementary Fig. 29** show that the maximum a posterior length scale $l'$ strongly correlates with the geometric mean nearest neighbor distance

$$\overline{\mathrm{dn}} = \exp\left(\frac{1}{n}\sum_{j=1}^{n} \log \circ \mathrm{dn}(x_j)\right)$$

across all tested datasets. We leveraged this insight through a regression

$$l' \approx \exp(\lambda) \cdot \overline{\mathrm{dn}} =: l\left(\overline{\mathrm{dn}}\right)$$

and arrived at $\lambda = 3$. This regression is intentionally without an intercept term to allow scale invariance of our density inference, and because the chosen length scale should approach 0 as $\overline{\mathrm{dn}}$ converges towards 0.

### Supplementary Note 5

### Synthetic Datasets with Known Ground Truth

To assess the accuracy of Mellon, we generated synthetic datasets exhibiting complexity similar to that of real-world single-cell datasets. For this purpose, Gaussian Mixture Models (GMMs) were employed, owing to their ability to emulate intricate structures often seen in actual single-cell data.

Three styles of simulation were adopted. The first approach generates random Gaussians, resulting primarily in isolated clusters when sampled. The other two styles are designed to replicate the architecture of biological differentiation systems, akin to our T-Cell depleted bone marrow dataset and the CD34+ bone marrow dataset. These styles implement a tree structure, as detailed in the subsequent sections.

#### 5.1 Constructing Gaussian Mixture Models

The 'cluster-style' synthetic datasets are created from a random ensemble of mean vectors and covariance matrices. The mean vectors are sampled from a standard normal distribution, while the covariance matrices are generated using the sklearn's make_spd_matrix function. This function employs uniform sampling and singular value decomposition to create symmetric, positive-definite matrices.

For the other two simulation styles, the structure of the GMM is carefully tailored to resemble a cellular differentiation tree. This also involves devising a mean vector and a covariance matrix for each Gaussian component of the model. Here, a tree is defined as a tuple containing an integer and a list of subtrees $(n_i, [T, ...])$, with each subtree $T$ being similarly structured.

Each node $i$ in the tree corresponds to a Gaussian, for which we calculate a mean $\mu_i$ and a covariance matrix $\Sigma_i$. These parameters are derived from a base point $b_i \in \mathbb{R}^{d'}$ and a velocity vector $v_i \in \mathbb{R}^{d'}$. The base point and velocity vector play crucial roles in shaping the underlying structure of the differentiation tree, thus influencing the complexity and characteristics of the simulated datasets.

#### 5.2 Generation of Base and Velocity Series

We used a sequential procedure to generate the base points (representing node locations) and velocity vectors (indicating the direction of simulated differentiation). These vectors define the structure of the differentiation tree:

$$\begin{aligned} \epsilon_i &\sim N(0, \sigma_i), & \epsilon_i &\in \mathbb{R}^{d'}, \\ v_i &= v_{P(i)} + \epsilon_i, & v_0 &= \mathbf{1}, \\ b_i &= b_{P(i)} + v_i, & b_0 &= \mathbf{0}. \end{aligned}$$

In the above expressions, $P(i) < i$ represents the parent node of node $i$, with 0 being the root node. We define $\sigma_i$ as $\|v_{P(i)}\|_2 \cdot \gamma$, where $\gamma$ is a positive real number reflecting the average curvature of the simulated differentiation trajectory. For datasets replicating T-Cell depleted bone marrow and CD34+ bone marrow, we chose $\gamma = 0.2$ and $\gamma = 0.5$, respectively.

#### 5.3 Formulation of the Covariance Matrix and Mean

To formulate a Gaussian distribution for each node $i$, we initiate with the velocity vector $v_i$ and construct an orthonormal basis via comprehensive QR-factorization. This operation yields a set of unit vectors $(\hat{u}_i^j)_{j\in(1,\dots,d')}$, which stand as the principal components of the Gaussian.

Given that the intrinsic dimensionality of a cell-state manifold is generally low, we scaled the Gaussians' principal components with a scalar that decays exponentially. This step is represented by the following equation:

$$u_i^j := \hat{u}_i^j \cdot \left[0.5 + \exp\left(\frac{-j \cdot \lambda}{2}\right) \cdot \|v\|_2\right].$$

We selected the variance decay rate, $\lambda$, as 1.5 for datasets that mimic T-Cell depleted bone marrow and 1 for datasets that resemble CD34+ bone marrow. Each Gaussian at node $i$ isdefined by its covariance matrix and mean, as outlined below:

$$\Sigma_i := (u_i^j)_{j\in(1,\ldots,d')} \cdot (u_i^j)^T_{j\in(1,\ldots,d')}$$
$$\mu_i := b_i + \frac{v_i}{2}.$$

#### 5.4 Ground Truth Density and Data Sampling

We sampled $n_i$ cells from the Gaussian distribution $N(\mu_i, \Sigma_i)$ for each node $i$. These samples together form the simulated dataset $X$. The ground truth density was defined as:

$$f(x) := \frac{1}{\sum_i n_i} \sum_i n_i f(x|\mu_i, \Sigma_i).$$

#### 5.5 Density Inference with Mellon and Performance Evaluation

Mellon was employed to infer the log-density for the simulated dataset $X$. We then compared this inferred log-density with the log-transform of the ground truth density $\log \circ f(x)$. The Spearman correlation was used to evaluate the agreement between these two for all $x \in X$. This comparison helped us evaluate Mellon's precision and reliability in estimating cell densities from high-dimensional single-cell data. [1]
